# An engine for systematic discovery of cause-effect relationships between brain structure and function

**DOI:** 10.1101/2025.04.05.647237

**Authors:** Andrea I. Luppi, Filip Milisav, Laura E. Suarez, Golia Shafiei, Jakub Vohryzek, Yonatan Sanz Perl, Hana Ali, Fernando E. Rosas, Pedro A. M. Mediano, Bratislav Misic, Gustavo Deco, Morten Kringelbach

## Abstract

Characterising how perturbations of brain architecture influence brain function is essential to understand the origins of brain dysfunction, and devise potential avenues of treatment. Here we introduce a computational engine for systematic causal discovery of the functional consequences of altering network architecture and local biophysics in the brain. We integrate multimodal anatomical and functional neuroimaging to implement over 2, 000 *in-silico* brains, and provide mechanistic insight into the functional consequences of local lesions, global wiring, and empirically-derived maps of regional cytoarchitecture and chemoarchitecture. We comprehensively assess how each manipulation of brain macrostructure reshapes spatial and temporal signal coordination, information dynamics, and functional hierarchy—as well as spontaneous co-activation of meta-analytic cognitive circuits, and *>* 6, 000 dimensions of local neural dynamics. Our computational model systematically identifies which features of brain architecture have overlapping or antagonistic causal influence over each dimension of brain function, and how functional properties are traded off against each other across disorders and neuromodulation. We find that regions’ functional vulnerability to lesions *in silico* recapitulates their vulnerability to neurodevelopmental and psychiatric *in vivo*, along a core-periphery organisation. We provide convergent evidence that the brain’s wiring diagram is finely tuned to favour the hierarchical integration of information. Notably, our model successfully recapitulates known empirical results that have not been modelled before, including desynchronisation and flattening of the brain’s functional hierarchy induced by psychedelic 5*HT*_2*A*_ agonists. To catalyse future discoveries, we make this resource freely available to the neuroscience community through an interactive website (https://systematic-causal-mapping.up.railway.app/), where users can interrogate our systematic database of simulated cause-effect relationships. Altogether, we provide a powerful computational engine to predict the functional consequences of experimental or clinical interventions, and drive neuroscientific hypothesis-generation.

## INTRODUCTION

A central goal of neuroscience is to obtain a mechanistic understanding of how coordinated brain activity emerges from the architecture of the human brain. Characterising how interventions on brain structure will influence brain function is also imperative for clinical and translational neuroscience, to identify successful avenues of treatment. However, a major obstacle to this endeavour is the brain’s heterogeneity and causal degeneracy. At the microscale, individual brain regions exhibit distinctive profiles of cyto-and chemo-architecture, [1– 12], shaping the biophysical and computational properties of local circuitry [13–23]. At the macroscale, specialised brain regions interact over a complex network of anatomical connections: the structural connectome [13, 16, 24–27]. Additionally, causal relations between brain structure and function are many-to-many. The same structural scaffold supports a diversity of functional configurations, as brain regions form and dissolve coalitions to execute different cognitive operations [13, 16, 24, 28–34]. Conversely, non-overlapping focal lesions can produce overlapping functional symptoms [35–38]. Thus, there are both convergent and divergent relationships between brain structure and function [39– 41].

Overcoming the challenge of causal degeneracy requires developing a systematic mapping between changes in brain macrostructure (regional biophysics, inter-regional wiring) and their consequences on brain function. However, avenues for experimental intervention on brain architecture are severely limited *in vivo*— especially for global properties such as the network topology of the connectome. In humans we are restricted to pharmacology, or naturally occurring variation in brain structure due to lesions and surgery, development, or neurocognitive disorders. Some manipulations remain even beyond the greater experimental accessibility of animal models.

Computational models of brain activity are ideally suited for mechanistic investigation of the links between brain architecture and function. They provide complete access for both manipulation and recording beyond what is biologically possible—including rewiring the entire connectivity of the brain into any desired configuration [42–53]. However, individual studies rarely use the same model and focus on hand-picked structural and functional properties of interest. This lack of methodological standardisation limits the field’s capacity to compare, generalise, and synthesise results from the modelling literature as a whole.

To overcome these challenges, here we execute a comprehensive structure-to-function causal mapping *in silico*, characterising how brain regions act, interact, and relate to each other along multiple dimensions. To ensure a standardised baseline for comparison, we use the same implementation throughout: a dynamic mean-field model (DMF) of excitatory and inhibitory populations, representing cortical regions and coupled according to the wiring diagram of the human connectome reconstructed from *in vivo* tractography (Fig. 1a) [63, 64]. Being amenable to separate manipulations of regional excitation, inhibition, and wiring, the DMF model provides an ideal balance of biophysical realism and computational tractability. This computational tractability enables us to systematically manipulate network connectivity and local biophysics across over 2, 000 ‘in silico brains’, and then map the effects of each intervention onto the most comprehensive battery of functional readouts to date (Fig. 1a). Specifically, we consider nearly 100 distinct manipulations of brain structure, which include:

**Figure 1.**
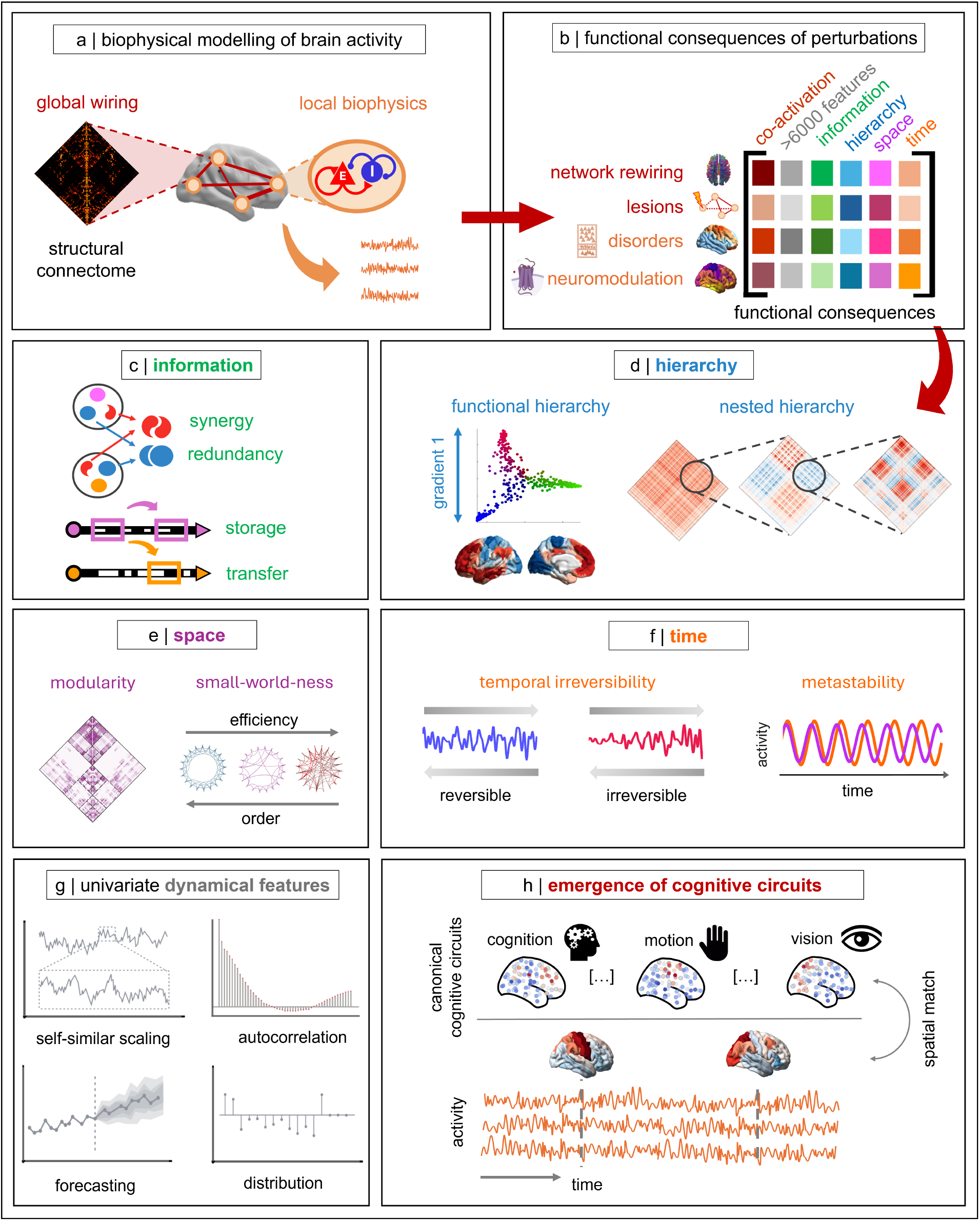
Systematic causal mapping of structural perturbations to their functional consequences with computational brain models. (**a**) The biophysical mean-field model comprises local excitatory and inhibitory neural masses, representing brain regions, coupled according to the empirical wiring of the human brain reconstructed from diffusion tractography. (**b**) The model’s local and global properties are systematically perturbed: (1) by rewiring or scaling the global connectivity; (2) by lesioning the local connectivity of each region; (3) by tuning each region’s recurrent excitation according to empirical maps of disorder-related cortical atrophy; (4) by modulating regional excitation-inhibition balance according to PET-derived maps of neurotransmitter receptor density. Each perturbation is systematically mapped onto informational, hierarchical, spatial, and temporal aspects of global functional organisation, as well as thousands of individual dynamical features, and the expression of meta-analytic brain patterns recapitulating fundamental cognitive operations. (**c**) Example of information measures: redundancy, synergy, information storage, and information transfer [54] (**d**) Example measures of hierarchical organisation: the range of the principal gradient of functional connectivity (FC) is used as proxy for the unimodal-transmodal functional processing hierarchy [55]; and the FC can be divided into a hierarchy of nested modules [56] (**e**) Example measures of spatial signal coordination: network modularity, and network small-world organisation due to the presence of shortcuts between local clusters. (**f**) Example temporal measures: temporal (ir)reversibility of the signal [57], and metastability (here quantified as the temporal variance of the global synchrony) [58, 59]. (**g**) Representative examples from our comprehensive sampling of univariate dynamical features. Features are obtained from the *highly comparative time-series analysis* toolbox (`hctsa`) covering the broad literature on time-series analysis in neuroscience

1. increasing or decreasing the global effective coupling between all regions;
2. rewiring the global network topology into specific configurations;
3. lesioning the local connectivity of each cortical region;
4. implementing changes in cortical cytoarchitecture from cortical thickness reflecting 11 neurological, psychiatric, and neurodevelopmental diagnostic categories across over 17 000 patients [65–67]
5. implementing ‘virtual pharmacology’ by tuning regional excitation and inhibition according to 15 maps of neurotransmitter receptor density, obtained from *in vivo* positron emission tomography (PET) scans in *>* 1 200 participants [2]

On the functional side, we map each macrostructural manipulation onto four key dimensions of global brain functional organisation:

1. Information dynamics (integrated information; synergy; redundancy; transfer entropy; and active information storage) [54, 68] (Fig. 1b)
2. Temporal signal coordination (synchrony, metastability, intrinsic timescale, and temporal irreversibility) [57–59, 69–71] (Fig. 1c);
3. Spatial signal coordination (prevalence of anticorrelations; modularity and small-world propensity of the functional connectome) [72–74] (Fig. 1d)
4. Functional hierarchical organisation (local-global propagation of intrinsic “ignition” events; principal gradient of functional connectivity; and nested modular organisation) [55, 56, 75](Fig. 1e).

We further complement these global read-outs by:

[start=5]Assessing thousands of univariate time-series features of brain dynamics [60, 61] (Fig. 1f); Quantifying the brain’s spontaneous self-organisation into macroscale functional circuits associated with fundamental cognitive operations, defined using the NeuroSynth meta-analytic engine [62, 76] (Fig. 1g).

This systematic structure-to-function mapping establishes a resource for computational ‘reverse inference’. Researchers will be able to consult our comprehensive look-up table of over 500 000 combinations of structural interventions and functional read-outs, to identify which causal manipulations would be more or less consistent with an empirically observed effect, overcoming the need to develop ad-hoc models for each new empirical observation. This resource is freely available to the neuroscience community through our interactive website (https://systematic-causal-mapping.up.railway.app/). At the same time, this work provides a computational engine to predict the functional consequences of experimental or clinical interventions, and drive hypothesis-generation. Altogether, we integrate multimodal neuroimaging data (functional MRI, diffusion tractography, cortical morphometry, PET) to systematically discover which aspects of brain structure have overlapping or antagonistic causal influence over each dimension of brain function, and how functional properties are traded off against each other.

## RESULTS

### Overview: Computational modelling to integrate multimodal brain structure and function

We use biophysical modelling to provide mechanistic insight into the functional consequences of manipulating the brain’s local and global wiring, cytoarchitecture, and chemoarchitecture, across an extensive battery of readouts: regional activity, inter-regional interactions, and the emergence of cognitively relevant circuits. Network-based brain models generate biologically plausible simulations of brain activity by combining two key ingredients: (i) a mathematical representation of the local biophysics of each brain region; and (ii) a wiring diagram of the anatomical connectivity between regions [44, 50, 64]. Models can vary widely in terms of complexity and biological realism. Specifically, here we simulate BOLD signals for 68 anatomically-defined regions of the Desikan-Killiany cortical atlas [77], using a dynamic mean-field (DMF) model of excitatory and inhibitory populations [18, 63, 64]. A cortical model ensures that the same model can be used to integrate and compare all our datasets, some of which are only available for the cortex. Regions are then coupled according to the empirical structural connectivity between regions of the human brain, obtained as a consensus connectome from 100 individuals of the Human Connectome Project [78], reconstructed from *in vivo* diffusion MRI tractography (*Methods*).

The DMF model has a free parameter, known as ‘global coupling’ and denoted by *G*, which scales all weights in the structural connectivity to account for the effectiveness of signal transmission between brain regions. To tune the model, we identify the value of *G* where the model produces the most realistic brain dynamics. In the literature, there are many criteria for identifying the best-fitting value of the global effective coupling; for example, maximising metastability or the similarity between empirical and simulated FC [23, 75, 79–83]. To avoid the issue of circular analysis, here we explicitly choose not to use as fitting criterion any of the functional read-outs defined above, whose use we reserve for model assessment. Instead, we follow a well established procedure [18, 64, 84–86], minimising the Kolmogorov-Smirnov distance between the model’s functional connectivity dynamics (FCD; the correlation between instantaneous functional connectivity measured at different points in time) and the group-wise FCD obtained from fMRI BOLD signals of 100 individuals from the Human Connectome Project (see *Methods* for details). This approach captures both spatial and temporal features of the data, better differentiating between optimal and suboptimal values of *G* than alternative fitting methods [18]. Since the model with *G* = 1.6 produces dynamics that most faithfully recapitulate the empirical spatiotemporal dynamics of human BOLD signals (Fig. S1), all our analyses are performed starting from a model tuned with *G* = 1.6.

### Functional consequences of regional biophysics

#### Role of neurotransmitter systems: ‘Virtual pharmacology’ from gradients of receptor expression

A prominent advantage of the dynamic mean-field model is that it can be enriched with regionally heterogeneous excitatory and inhibitory dynamics. What are the functional consequences of heterogeneous modulations of local biophysics? This question bears clear clinical relevance for our mechanistic understanding of pharmacological interventions. Pharmacology is a cornerstone of modern clinical practice, being routinely and widely used to treat psychiatric and other mental health conditions. Many pharmacological agents reshape cognition and behaviour by engaging the brain’s rich array of neurotransmitter receptors, reshaping local biophysics [87–89]. It is therefore of great value to understand the functional consequences of engaging different neurotransmitter systems.

However, even when a drug’s molecular targets are relatively specific and well-characterised, it can be challenging to translate from microscale effects of engaging a specific receptor *in vitro*, to the drug’s macroscale effects on the human brain *in vivo*. One of the many reasons is that receptor expression is not uniform across the brain, but rather each region is characterised by a unique profile of receptor expression [1–3]. Computational modelling can contribute to bridging this gap between *in vitro* and *in vivo*. For example, biophysical whole-brain models with heterogeneous dynamics based on the empirical distribution of specific receptors across the cortex can recapitulate the effects of serotonergic, dopaminergic, nicotinic, or GABA-ergic agents on macroscale brain activity [18, 19, 80, 81, 90–93].

Here, we adopt the same approach for our ‘virtual pharmacology’. We systematically characterise how diverse aspects of brain function are reshaped by engaging each of 15 excitatory and inhibitory neurotransmitter receptors across 8 neurotransmitter systems, quantified from *in vivo* PET across over 1200 human volunteers [2] (Fig. 2a). Following the approach of [18], the effect of engaging an excitatory receptor (*mGluR*_5_, *NMDA, α*_4_*β*_2_, 5*HT*_2*A*_, 5*HT*_4_, 5*HT*_6_ *D*_1_, *M*_1_) is modelled by increasing the excitatory gain parameter of each region, according to that region’s receptor density (normalised between 0 and 1). Likewise, the effect of engaging an inhibitory receptor (*GABA*_*A*_, *D*_2_, *MOR, CB*_1_, *H*_3_, 5*HT*_1*A*_, 5*HT*_1*B*_) is modelled by increasing the inhibitory gain parameter of each region, according to that region’s receptor density (normalised between 0 and 1) - as per [91]. Note that no additional parameter tuning is performed, but rather we directly use the normalised values for each map.

**Figure 2.**
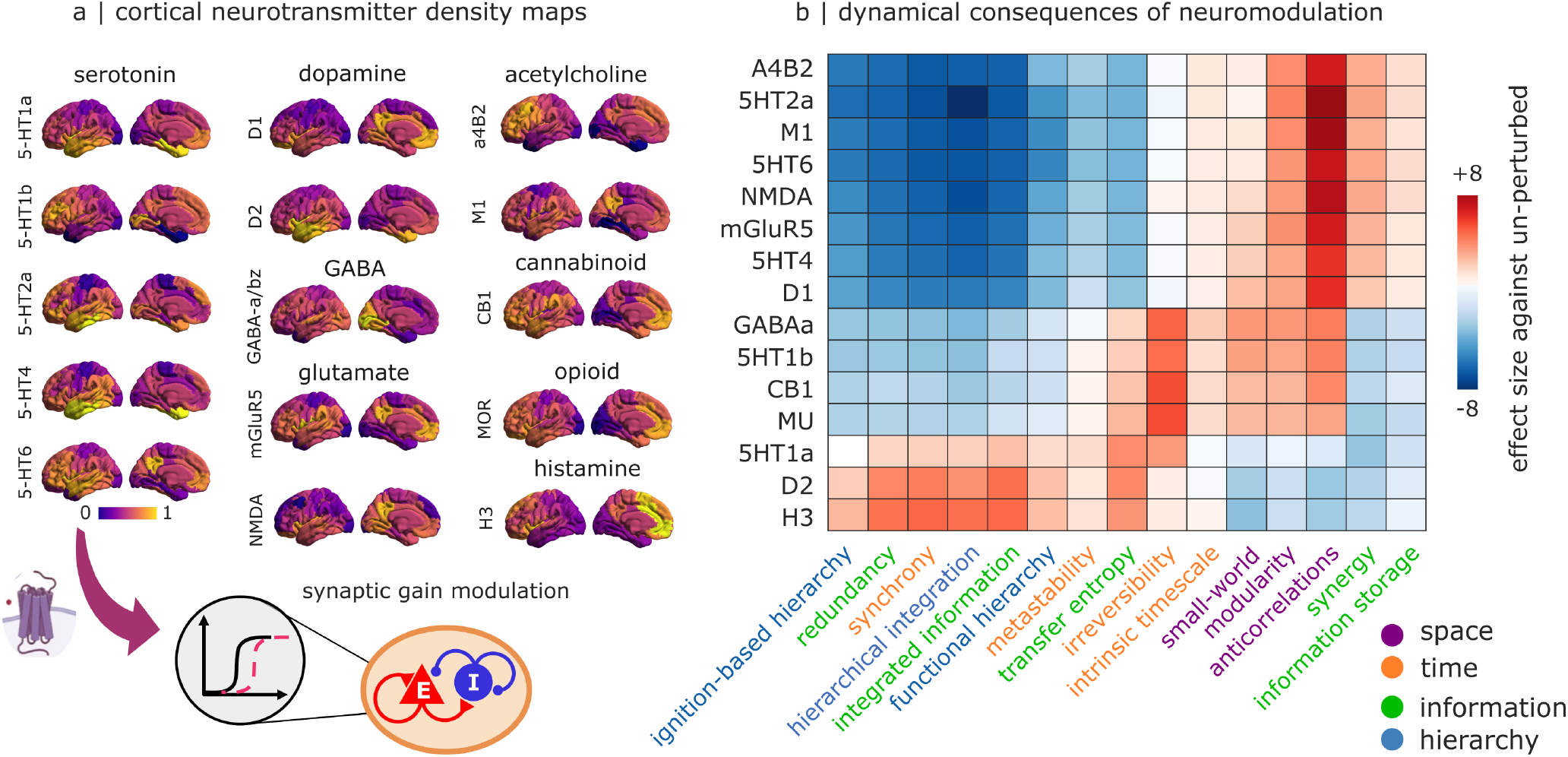
Functional consequences of engaging neurotransmitter systems. | **(a)** The effect of engaging each excitatory (resp., inhibitory) neurotransmitter receptor are modelled by varying the excitatory (resp., inhibitory) gain of each region, in proportion to its normalised receptor density measured from *in vivo* PET [2]. The regional distribution of each receptor is shown on the cortical surface. **(b)** Virtual pharmacology recapitulates known empirical results, including desynchronisation and flattening of the brain’s functional hierarchy induced by 5*HT*_2*A*_ agonists. Three distinct groups of neurotransmitter receptors can be discerned from their functional consequences. Heatmap shows how modulating regional excitatory or inhibitory gain according to empirical receptor density reshapes the functional organisation of the brain, in terms of effect size (Hedge’s *g*) against the un-perturbed model.

Our computational assessment indicates that nearly every measure of brain functional organisation can be either increased or decreased through virtual pharmacology, depending on which neurotransmitter receptor is engaged—with three broad groups of receptors being evident, consisting of the excitatory receptors, and two groups of inhibitory receptors (Fig. 2b). However, functional measures differ in terms of their overall susceptibility to virtual pharmacology. Some exhibit comparatively weak responses throughout (e.g., metastability, intrinsic timescale), or comparatively strong responses throughout (e.g., anticorrelations, hierarchical integration). In contrast, other aspects of brain functional organisation appear relatively unaffected by all but a select few receptors, such as temporal irreversibility being selectively responsive to GABA_A_, 5-HT_1A_, 5-HT_1B_, CB_1_ and *MU* receptors (Fig. 2b). Overall, the fact that each of the functional properties considered here can be manipulated bi-directionally, lends credence to the possibility of using *in silico* pharmacology to devise treatments for re-balancing brain function.

Crucially, many of our *in silico* predictions are successfully validated against the empirical consequences of pharmacological interventions *in vivo*. Reductions of synchrony and hierarchical integration, and contraction of the functional hierarchy, have been reported for several anaesthetics including GABA-ergic agents [94–96]. These results are consistent with our model’s predictions from engaging *GABA*_*A*_ receptors. We note that anaesthesia also tends to reduce anticorrelations [74], whereas our model predicts an increase. However, the same prediction of increased anticorrelations under heterogeneous GABA receptor engagement was also made by the model of [92], in line with our own results. Our model further predicts that global synchrony of fMRI signals should be substantially reduced by agonism of the 5*HT*_2*A*_ receptor—and indeed, a recent large-scale study of the serotonin 2A receptor agonist psilocybin reported that psilocybin desynchronises the human brain [97]. 5*HT*_2*A*_ receptor agonsist DMT, LSD, and psilocybin also ‘flatten’ the functional hierarchy of the human brain [98, 99]: this is again consistent with our model’s results.

#### Role of intrinsic excitability: ‘Virtual patients’ from gradients of cortical thickness abnormality

Although pharmacology can temporarily modulate regional biophysics, another another clinically relevant application of our model is to assess the functional consequences of cytoarchitectonic heterogeneity. There is growing evidence that many neurodevelopmental, neurodegenerative, and psychiatric disorders are not restricted to single regions, but rather affect the entire cortex, often with complex combinations of atrophy and enlargement [65–67, 100]. To model realistic perturbations of local biophysical properties, we turn to the ENIGMA consortium’s large-scale database of changes in cortical thickness associated with 11 neurological, neurodevelopmental, and neuropsychiatric diagnostic categories [65–67, 100] (Fig. 3a). Our approach is inspired by recent work that employed atrophy maps from Alzheimer’s disease and frontotemporal dementia to modulate local parameters in a whole-brain model [101]. For each diagnostic category, we generate ‘virtual patients’ by modulating the regional level of intrinsic excitability in the model according to the regional pattern of increases or decreases in cortical thickness associated with that condition (Fig. 3). Namely, when atrophy of cortical thickness is observed, we model it as reduced excitation; and when an increase in thickness is observed, we increase excitation in the corresponding region. While we acknowledge that this is inevitably a simplification, it is based on the rationale that most neurons are excitatory, and therefore increases or decreases in grey matter would also primarily involve excitatory neurons. Indeed, a similar approach has been adopted before to model the effects of cortical thickness abnormalities *in silico* [102].

**Figure 3.**
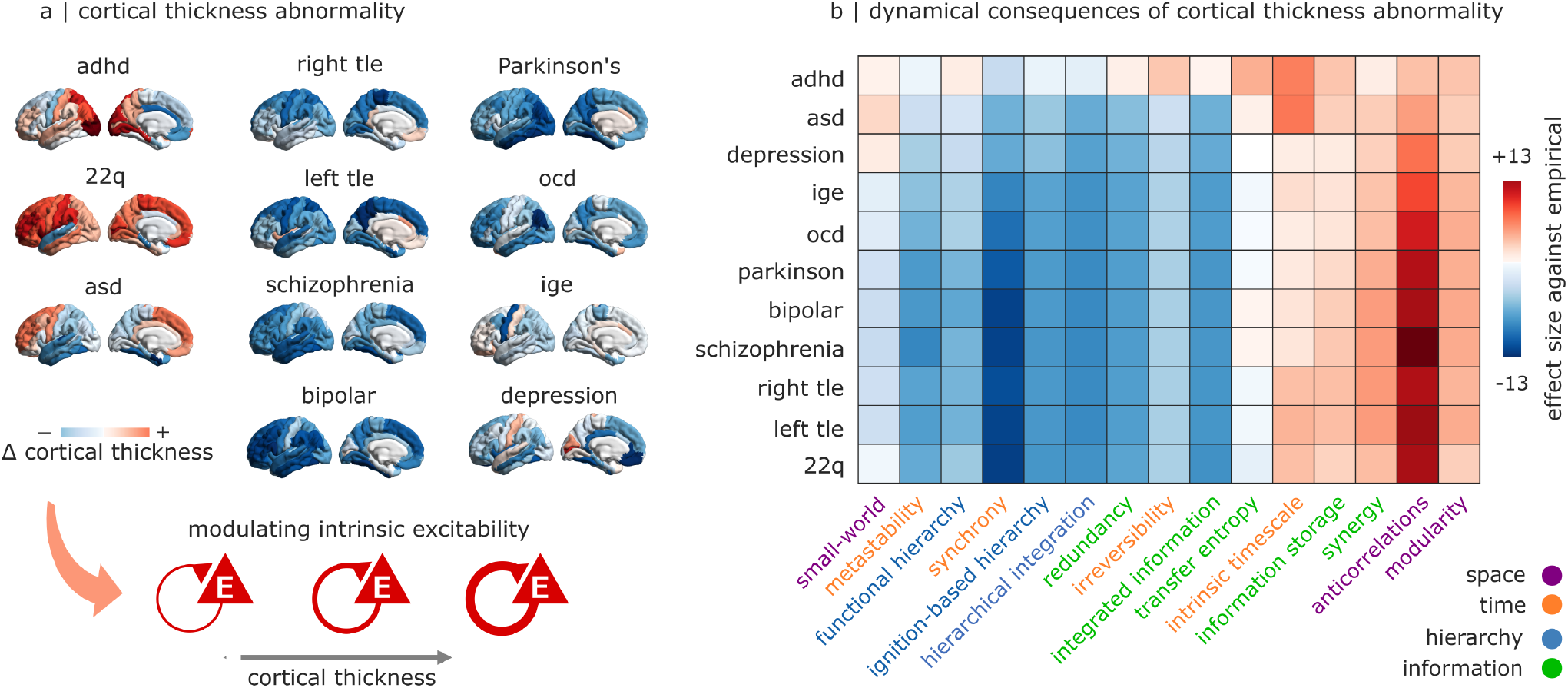
Functional consequences of changes in intrinsic excitability from cortical thickness abnormality. | **(a)** The effects of cortical thickness abnormality are modelled by varying the intrinsic excitation level of each region, in proportion to the extent of its cortical thickness alteration from healthy controls (Cohen’s *d*, as provided by the ENIGMA Toolbox [65]): atrophied regions have less excitation, and regions of increased thickness have greater excitation. The empirical changes in cortical thickness associated with each diagnostic category are shown on the cortical surface. (b) Synchrony and the prevalence of anticorrelations are especially susceptible to changes in intrinsic excitation. Heatmap shows how modulating regional excitation according to each empirical pattern of cortical thickness abnormality reshapes the functional organisation of the brain, in terms of effect size (Hedge’s *g*) against the un-perturbed model. adhd = attention deficit/hyperactivity disorder; asd = autistic spectrum disorder; ocd = obsessive- compulsive disorder; ige = idiopathic generalised epilepsy; right tle = right temporal lobe epilepsy; left tle = left temporal lobe epilepsy.

We find that different aspects of brain functional organisation are differentially affected by cortex-wide changes in regional intrinsic excitation. Some measures (e.g., modularity, prevalence of anticorrelations) tend to increase throughout, whereas others (e.g., synchrony, hierarchical integration) are reduced by most perturbations (Fig. 3b). However, even when the direction of change is relatively consistent, the magnitude can vary widely depending on the specific diagnostic category that is used for the perturbation. Overall, both the direction and magnitude of change are dependent on the specific combination of functional measure and regional pattern (Fig. 3b).

Some of the model’s computational predictions have already found empirical validation. Numerous reports indicate that schizophrenia and bipolar disorder are characterised by reductions of fMRI global synchrony and functional connectivity [103–107]—consistent with our model’s indication of reduced synchrony and redundancy (which is closely linked with FC). Our model also predicts that the cortical thickness changes associated with epilepsy should induce reduced fMRI synchrony and increased modularity. Surprisingly, this is in fact consistent with empirical evidence: although epilepsy is famously associated with EEG hyper-synchrony during seizures, fMRI studies instead indicate decreased FC as a common finding in patients at rest [108, 109]. More broadly, the similar functional consequences resulting from diverse patterns of cortical thickness abnormality is consistent with the growing recognition that most diagnostic categories have intricate co-morbidities [110– 112].

### Functional role of inter-regional connectivity

#### Global manipulations

Having considered one of the two main ingredients of whole-brain models—regional biophysics —we now turn to the other: inter-regional connections between regions. Given the prominent role of the global effective coupling (*G*) in determining model goodness-of-fit in the literature (for example, by finding the value of *G* that maximises some property such as synchrony or metastability [79, 83, 85]), we take the opportunity provided by our model to systematically evaluate the impact of global coupling on the broader set of functional properties of the brain. We simulate BOLD signals using models with *G* values ranging from 0.1 to 3.0, in increments of 0.1, which scales the strength of inter-regional connections in the model’s structural connectome to determine the effective connectivity between regions. At each value of *G*, we compute our entire battery of functional properties, to examine how they change as a function of inter-regional effective coupling; greater value of effective coupling indicates stronger signal transmission between regions.

We find that functional properties exhibit a diverse range of behaviours in response to changes in the global effective coupling. Whereas some measures monotonically increase or decrease with greater coupling (e.g., anticorrelations, functional hierarchy) others are relatively insensitive to changes in the effective coupling between regions, except at the very highest values (e.g., information storage, intrinsic timescale). Finally, several functional measures (including hierarchical integration and all information-theoretic ones) exhibit non-linear relationships with the effective coupling between regions, characterised by prominent peaks or valleys around *G* = 2.2: higher than the optimal working point of *G* = 1.6 where the KS divergence between empirical and simulated FCD is minimised (Fig. S2).

On one hand, the observation that many functional properties including metastability and integrated information peak around the same level of effective coupling, is consistent with previous modelling literature [113]. On the other hand, by considering a much broader repertoire of functional measures we can finally see that the bigger picture is more complex. The global level of effective coupling has drastically variable effects across different aspects of functional brain organisation. Indeed, many measures appear to exhibit their most extreme value away from the point where our model best fits the empirical data. Crucially, this observation is relevant beyond the specific dataset or criterion that we used for fitting, or the specific point of best-fit identified here: since different measures peak at different coupling levels, it follows that no single value of global effective coupling can simultaneously maximise all functional properties, regardless of how ‘best fit’ is defined. Rather, complex trade-offs exist between different functional properties.

The indication that the brain does not simultaneously maximise all its functional properties is not only an important neurobiological insight. It also bears consequences for computational modelling. It is common practice in the literature to fit models by maximising a single functional feature (such as synchrony or metastability [79, 83, 85]). However, our results highlight the need to look at the bigger picture, and take into account the brain’s broader repertoire of functional properties.

As an alternative approach to interrogate the functional role of global network organisation, we can also keep the global effective coupling fixed at the optimal working point, and instead rewire how regions are connected. This kind of global rewiring into a desired topology is not currently possible in living organisms, where we are limited to assessing naturally-occurring variation across individuals or across species. In contrast, by selectively disrupting or preserving any chosen features of the network, computational models provide a unique avenue to understand how specific aspects of global network organisation influence brain dynamics - highlighting the power and flexibility of this approach.

We rewire the connectome according to seven network rewiring schemes [114]: binary network (preserving topology but setting all weights to the same value), lattice network with preserved weight distribution, modular network with preserved weight distribution, and four types of increasingly constrained random networks: fully random (but preserving the distribution of edge weights) [115], weight-and degree-preserving [116], strength-preserving (i.e., preserving both a region’s number of connections and their combined weight) [117], and geometry-preserving (i.e., preserving degree and also connection length, to respect the spatial embedding of the empirical connectome) [118] (Fig. S3a).

Notably, we find that biophysical models based on the empirical human connectome exhibit the highest values of integrated information and redundancy, as well as most measures of hierarchical organisation (Fig. S3b). We also find that increasingly drastic interventions on the brain’s network structure incur a corresponding deterioration of the brain’s functional properties (Fig. S3c). It is noteworthy that the largest repercussions arise from binarisation, which preserves topology but erases the heterogeneity of weight distribution. The result is a prominent reduction in measures of synchrony and hierarchy, with increased prevalence of anticorrelations (Fig. S3b). However, the presence of heterogeneous edge weights alone is not enough to preserve the functional properties of the empirical human connectome: although this property is preserved by all other rewiring schemes than binarisation, profound functional differences still exist among them (Fig. S3c). Rather, it is also important to take into account how edge weights are distributed across nodes: functional differences with the empirical human connectome are minimised for the strength-preserving rewiring, which maintains each region’s weighted degree (sum of its edge weights; Fig. S3c). Overall, our *in silico* rewiring of the human connectome reveals the importance of edge weights, and how they are aggregated across regions.

#### Local structural lesions

Having addressed the dynamical consequences of local biophysics and global network organisation, we next consider a middle-ground between the two, by probing the role of local structural connectivity on a region-by-region basis. Whereas the global manipulations considered above are only possible *in silico*, perturbations of a region’s connectivity can be viewed as simulating the effects of a focal lesion. Concretely, we implement local perturbations as follows: for each region of the Desikan-Killiany atlas (combined across both hemispheres), we implement a “virtual lesion” by setting all its connections to 1*/*10 of their original strength. We then use the biophysical model to generate new regional BOLD signals based on the lesioned connectome, for*N* = 30 repetitions. From these simulated BOLD signals, we compute our entire battery of functional measures, and obtain an effect size (Hedge’s *g*, whose interpretation is analogous to Cohen’s *d*) from comparing them against the same measures obtained from the original (un-perturbed) connectome. Therefore, our results reflect the *brain-wide* effects of *local* lesions. To visualise the results, we plot them on the cortical surface such that each region’s colour reflects the statistical effect size (magnitude and direction) of the change in *brain-wide* measure, resulting from virtually lesioning that region 4a).

Our results indicate that most measures of brain function tend to be either consistently increased or consistently decreased by local structural lesions, with lesion location influencing the magnitude of change, but not its direction. For example, most lesions induce increases in network modularity, and reductions in synchrony—as might be expected since the underlying structural connectome is becoming less connected. However, we also observe exceptions where both notable increases and decreases can be seen (e.g., intrinsic timescale, synergy, active information storage; Fig. 4a).

**Figure 4.**
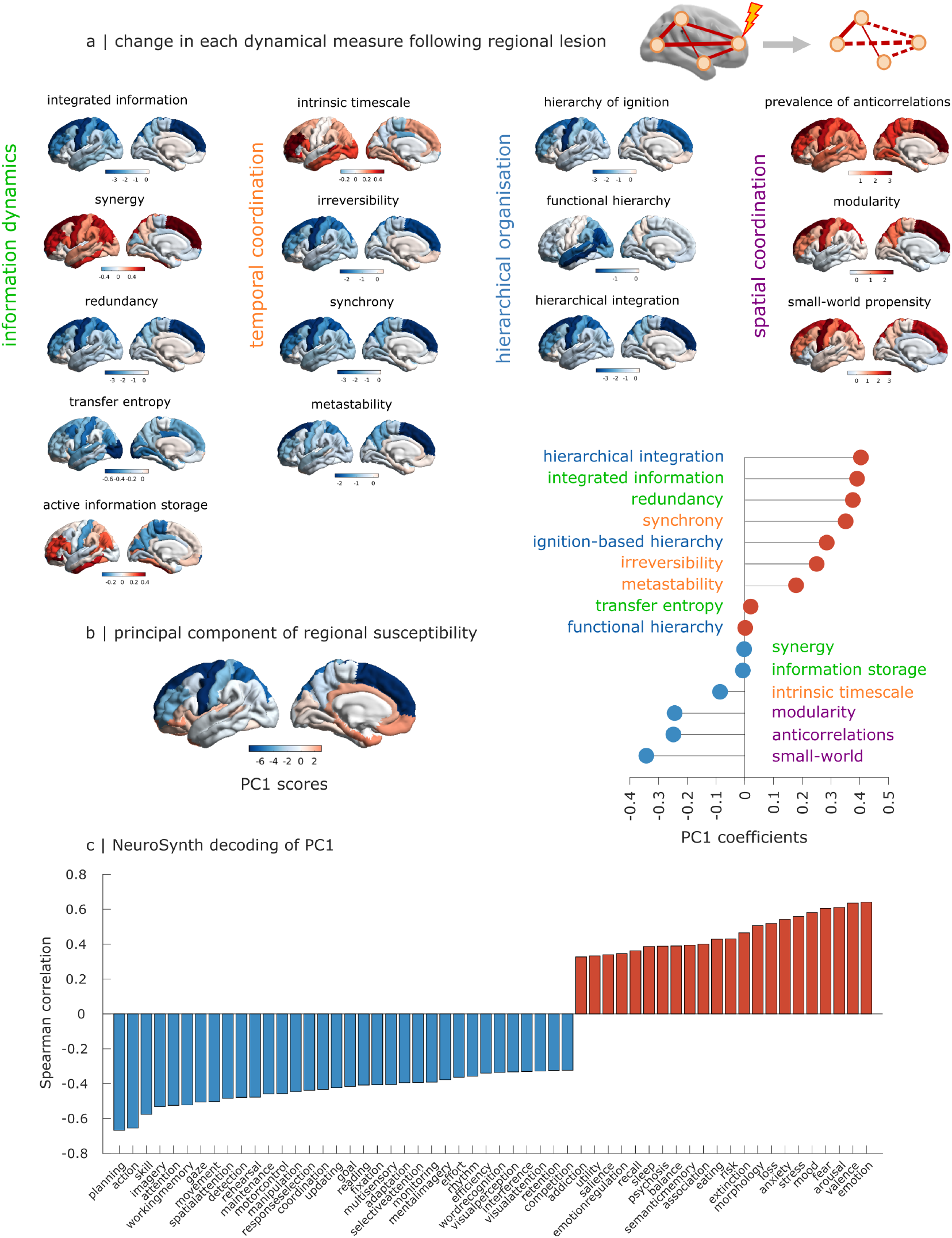
Functional consequences of local structural lesions. | (**a**) Most measures of brain function (e.g., synchrony) tend to be either consistently increased or consistently decreased by local structural lesions. Each brain map shows, for each cortical region, the effect size (Hedge’s *g*) from comparing whole-brain values obtained from *N* = 30 simulations with the empirical (un-perturbed) connectome, and *N* = 30 simulations from the locally lesioned connectome. Location on the brain map corresponds to the lesioned region (note that the same region is lesioned simultaneously in both hemispheres). Positive values (red) indicate lesioned *>* empirical, and negative values (blue) indicate lesioned *<* empirical. (**b**) The principal component of dynamical susceptibility to local perturbations delineates a core-periphery axis of cortical organisation. Brain scores are shown on the cortical surface on the left, and the contribution of each functional property to the principal component is shown on the right. (**c**) NeuroSynth decoding reveals that the principal axis of susceptibility to local lesions differentiates regions pertaining to externally-oriented versus internally-oriented modes of cognition. Bars indicate spatial correlation; statistical significance is assessed against spatial autocorrelation-preserving null models [119]; only significant terms that survive FDR correction across 123 total NeuroSynth terms are shown

To summarise the role of each cortical region in re-shaping global functional properties of the brain, we apply principal component analysis (PCA) on the regions-by-measures matrix. Principal components provide a low-dimensional representation that best explains the association between regions and functional measures. Regions with similar consequences on our battery of functional measures will exhibit similar PC scores. We find that a single principal component accounts for most of the variance (92%) in functional susceptibility to local structural lesions. This principal component of local susceptibility (‘susceptibility PC1’ for short) delineates a core-periphery axis of cortical organisation, being anchored in ventromedial cortices at one extreme, and lateral-dorsal cortices at the other extreme (Fig. 4b). The ventromedial end is primarily associated with measures of hierarchical organisation and temporal signal co-ordination, whereas measures of spatial coordination are associated with the dorsolateral end (Fig. 4b).

To characterise the cognitive relevance of this spatial pattern, we perform a term-based meta-analysis using 123 meta-analytic brain maps from the neuroimaging literature, available in the online database NeuroSynth [62]. A large number of NeuroSynth cortical patterns are significantly spatially correlated with the principal axis of functional susceptibility to local structural lesions, even after correcting for multiple comparisons and the effect of spatial autocorrelation [114, 119] (Fig. 4c). In particular, many of the terms significantly associated with the dorsolateral end pertain to goal-directed behaviour (‘action’, ‘skill’, ‘planning’, ‘expertise’, ‘goal’, among others). In contrast, terms associated with the ventromedial end pertain to emotion (‘arousal’, ‘fear’, ‘anxiety’, ‘valence’, ‘emotion’) and mental health (‘stress’, ‘psychosis’, ‘addiction’). Thus, we may characterise the principal axis of susceptibility to local lesions as differentiating regions pertaining to externally-oriented versus internally-oriented modes of cognition.

Since the models differ only in terms of which region’s connections are lesioned, any differences must be attributed to network properties. This ability to isolate the causal roles of different features is a key advantage of computational modelling. Therefore, we next seek to determine which aspects of the network organisation of the empirical (un-lesioned) structural connectome account for the regional pattern of functional susceptibility to lesions (summarised by the susceptibility PC1). Specifically, we consider several key aspects of regional network organisation: the weighted degree (sum of a node’s connection weights); local efficiency (a measure of how well interconnected a node’s neighbours are); participation coefficient (diversity of modules that a node connects to); and average and modal controllability, which account for connectivity across paths of varying length to estimate a node’s ability to steer network dynamics towards easy-to-reach (average controllability) or hard-to-reach (modal controllability) activation states [120]. To disentangle the respective contributions of these various structural properties, we use dominance analysis [121], which compares all possible combinations of predictors to identify their relative contribution to the total variance explained.

Our results indicate that the principal component of susceptibility to local lesions is best predicted by how well-connected each region is (both directly, in terms of nodes’ weighted degree, and indirectly, as quantified by average and modal controllability metrics), but with little regard for whether its neighbours are themselves connected, or whether they belong to diverse modules (Fig. S4b). It stands to reason that the best predictor of a region’s impact on functional measures is its weighted degree: regions with higher weighted degree will be those whose connections suffer the most drastic change in connectivity, in absolute terms. Demonstrating the empirical validity of this prediction, our virtual lesion results suggest that most structural lesions will reduce functional and ignition-based measures of hierarchy in the brain - consistent with reports in patients with severe brain injuries [122].

We also contextualise the susceptibility PC1 with respect to other large-scale patterns of cortical anatomy and function. Anatomically we consider intracortical myelination (quantified from in vivo T1w:T2w MRI ratio; [9]); the principal component of gene expression from the Allen Human Brain Atlas database (AHBA PC1 [10]); the principal component of receptor density from in vivo PET (receptor PC1; [2]). Functionally we consider the principal component of meta-analytic activation from the NeuroSynth database (NeuroSynth PC1 [62, 123]); and the principal gradient of functional connectivity [55]. Remarkably, we find that the principal component of susceptibility to local lesions is best recapitulated by the principal component of NeuroSynth meta-analytic activation, which alone accounts for more variance (>50%) than all other predictions combined (Fig. S4c).

Finally, we consider how regions resemble each other in terms of their functional susceptibility to structural lesions, by correlating each pair of regions across measures. This produces a matrix of co-susceptibility to structural lesions (Fig. S4d). Across regions, we find that regions’ co-susceptibility to structural lesions in our model, is significantly associated with regions’ co-susceptibility to abnormalities in cortical thickness across 11 neurological, neurodevelopmental, and neuropsychiatric diagnostic categories from the ENIGMA consortium, reflecting cortical morphometry data from over 17 000 patients [65, 67] (Fig. S4d). In other words, regions that induce similar functional changes when their connectivity is lesioned *in silico*, tend to be similarly susceptible to empirical cortical abnormality—suggesting that our model-based functional phenotyping may be capturing some fundamental underlying aspects of brain organisation.

### Reshaping univariate dynamical features of brain activity

Up to this point, we considered functional consequences in terms of cortex-wide functional organisation. Next, we evaluate the effects of each perturbation across a battery of 21 dynamical features of individual BOLD time-series [124]. These features were chosen to be representative of the broader literature on dynamical systems, encompassing *>* 6 000 time-series properties such as linear and nonlinear autocorrelation, periodicity, forecasting, and statistical distribution of the data-points [60, 61]—which we also compute and make available to the reader through our interactive website: https://systematic-causal-mapping.up.railway.app/ (Fig. 5a; see Fig. S7 for investigation of the relationship between local dynamics and global functional connectivity).

**Figure 5.**
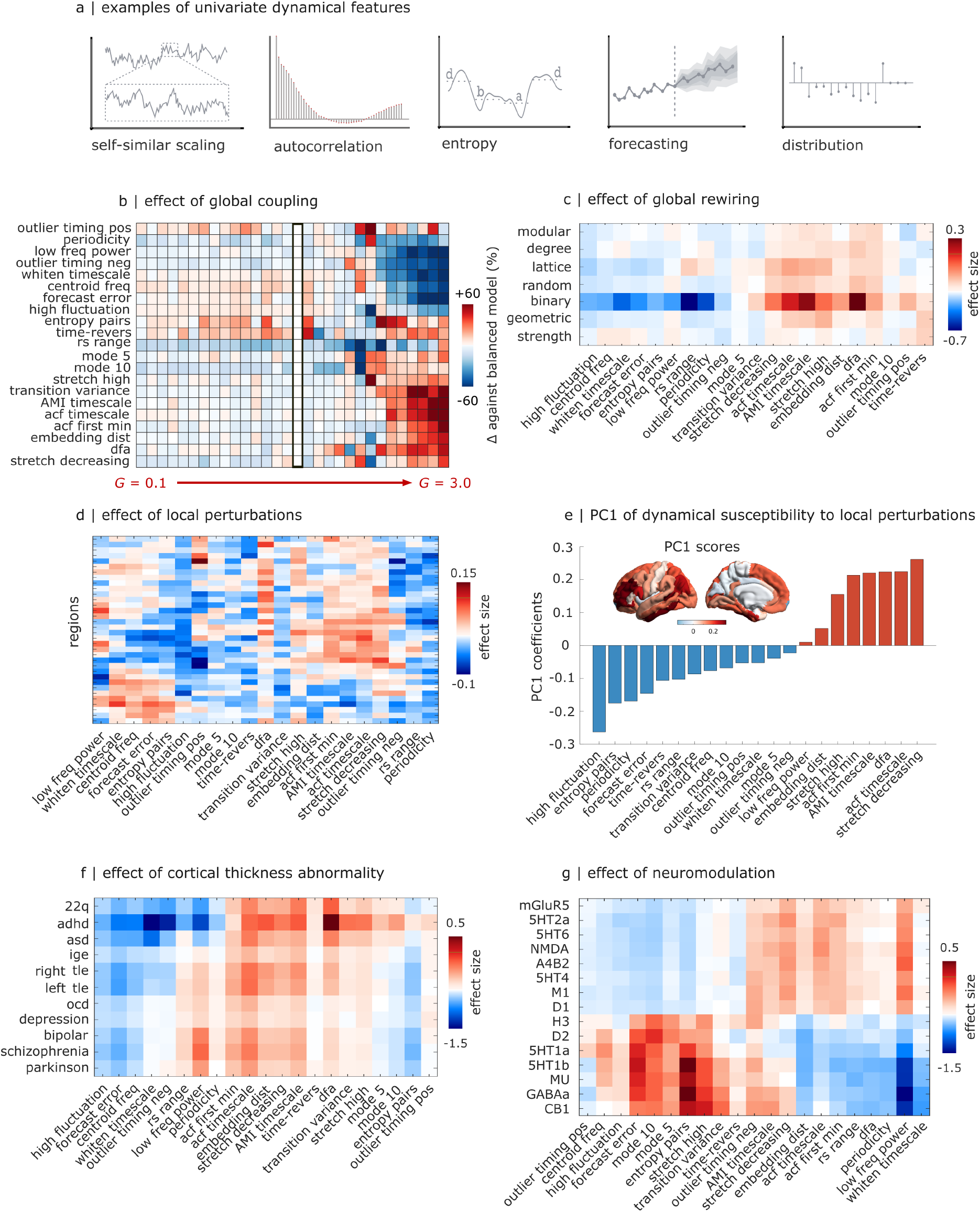
Effects of model perturbations on dynamical features of neural time-series. | **(a)** Examples of dynamical features of univariate time-series include temporal autocorrelation, entropy, and statistics of the data distribution. **(b)** Dynamical features become most extreme as global effective coupling increases. Percent change in each univariate dynamical feature is shown as a function of the global effective coupling parameter *G*, compared against the value observed for the best-fitting model (*G*=1.6). **(c)** Dynamical features are especially sensitive to the distribution of weights, being most perturbed by binarisation rather than any kind of weight-preserving rewiring. Heatmap shows the effect of global network rewiring across a battery of 21 univariate dynamical features (each averaged across all brain regions), in terms of effect size (Hedge’s *g*). **(d)** Heatmap shows the effect (Hedge’s *g*) across a battery of 21 univariate dynamical features, upon lesioning the structural connectivity of each cortical region. **(e)** A localised-versus-distributed principal component summarises the effects of local structural lesions on univariate dynamics, accounting for 35% of the variance. **(f)** Reshaping intrinsic regional excitation according to cortical thickness abnormality can either increase or decrease each feature, but not both. Heatmap shows the effect (Hedge’s *g*) on our 21 dynamical features, upon heterogeneously increasing or decreasing intrinsic excitation of each region, according to empirical patterns of cortical thickness abnormality. **(g)** Nearly every univariate dynamical feature can be both increased and decreased by virtual pharmacology, depending on which empirical receptor maps are used to modulate excitatory or inhibitory gain. Heatmap shows the effect (Hedge’s *g*) across our battery of 21 dynamical features [124], upon simulating engagement of each neurotransmitter receptor. adhd =

We find that dynamical features vary in their sensitivity to changes of the global effective coupling (Fig. 5b). As with global functional organisation, the value of the effective coupling parameter *G* where most univariate dynamical features exhibit maxima or minima is higher than the optimal value of *G* = 1.6 where the fit between simulated and empirical FCD is maximised (Fig. 5b). However, many global functional features tend to peak around *G* = 2.2, whereas most univariate dynamical features exhibit the most pronounced changes beyond this point, at the very highest values of global coupling. Thus, we find that univariate and global aspects of brain function are differentially sensitive to the level of global effective coupling between regions (Fig. 5b).

Pertaining to global network organisation, we find that for most univariate features, the most extreme values (highest or lowest) are produced by biophysical models based on the empirical human connectome, such that different rewirings will only decrease (resp., increase) the univariate feature (Fig. 5c). This is consistent with what we observed for global features, as is the fact that dynamical features are especially sensitive to the distribution of weights, being most perturbed by binarisation rather than any kind of rewiring that preserves the weight distribution (Fig. 5c).

In contrast, we find that many univariate dynamical features can exhibit both increases and decreases as a result of local structural lesions, depending on lesion location (Fig. 5d). This pattern represents a departure both from what is observed with global rewiring, and from what is observed for global features, for which more lesions tend to induce either increases or decreases, but not both. As a result, the principal component of univariate dynamical susceptibility to local lesions is different from the principal component of global functional susceptibility (Fig. 5e). Instead of a broad ventromedial-versus-dorsolateral division, we observe a localised-versus-distributed spatial pattern. Namely, one extreme is localised to the precuneus and the neighbouring visual and somatosensory medial cortices, driven by features such as periodicity and forecasting error. The other extreme is instead more distributed, exhibiting three separate peaks in lateral prefrontal, angular gyrus, and inferior temporal cortices. This end of the principal component is primarily driven by linear and nonlinear autocorrelation and intrinsic timescale (Fig. 5e).

When we consider the effects of virtual disease, implemented as regionally heterogeneous perturbations of local biophysics based on empirical cortical thickness abnormality, the results on univariate time-series features once again resemble those observed for global functional organisation. Namely, reshaping intrinsic regional excitation according to cortical thickness abnormality can either increase or decrease each feature, but not both, instead primarily differing in the magnitude of induced change (Fig. 5f). In contrast, nearly every univariate dynamical feature can be both increased and decreased by virtual pharmacology, depending on which empirical receptor maps are used to modulate excitatory or inhibitory gain (Fig. 5g).

Here we focused on the results of univariate features after aggregating them across the entire brain. However, there are undoubtedly rich patterns of structure-function relationships to be found, by looking at the spatial distribution of individual features across the cortex. Therefore, we release the results of our simulations as a resource for the scientific community: users can freely explore all results through our interactive website at https://systematic-causal-mapping.up.railway.app/.

### Emergence of functional circuits as an alternative window onto brain function

Finally, up to this point we considered brain function in terms of how brain regions act and interact (i.e., the sense in which the field typically refers to ‘structure-function relationships’ [13, 24, 26]. However, a complementary way to model brain function was adopted by Luppi et al. [102], who used network control theory to induce patterns of regional brain activity corresponding to different cognitive operations, derived from meta-analytic aggregation. Indeed, it is well established in cognitive and systems neuroscience that different brain regions form functional circuits that support different cognitive operations. Even at rest, regions exhibit preferential fMRI co-activation with other regions belonging to the same cognitive circuit [125–128].

Inspired by Luppi et al. [102], here we use the automated meta-analytic engine NeuroSynth to extract brain patterns reflecting the association between each cortical region, and a set of fundamental cognitive operations: attention, cognitive control, emotion, vision (fixation), language, memory, and movement (shown in Fig. 6a). We then use our model to ask how each perturbation affects the ability of regions belonging to the same cognitive circuit to co-activate together. Practically, we do so by quantifying at each point in time, the magnitude of the spatial correlation between spontaneous activity and each meta-analytic pattern [76].

**Figure 6.**
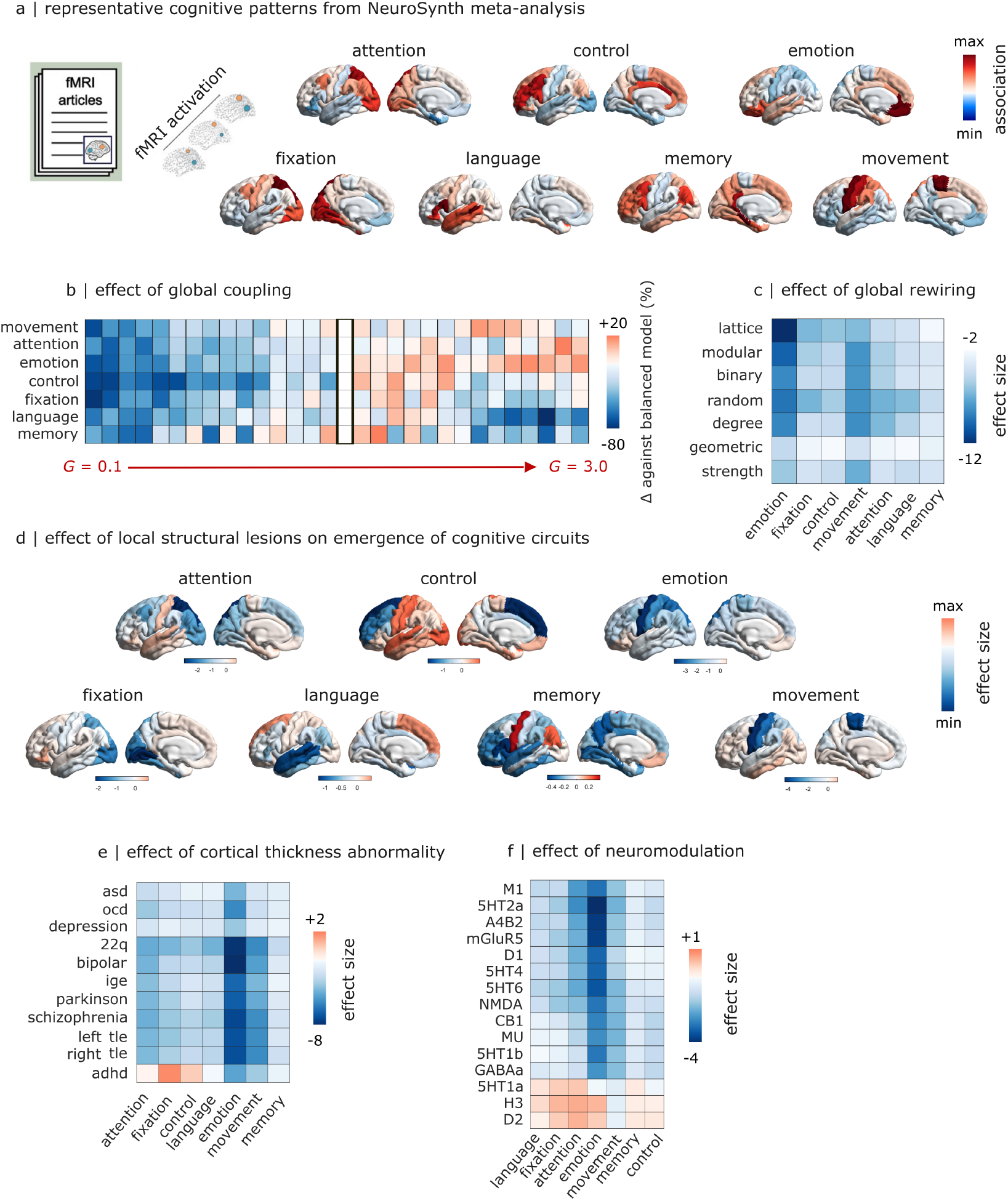
Effects of model perturbations on the emergence of cognitive functional circuits. | **(a)** NeuroSynth meta-analytic patterns for representative cognitive operations (values are z-scored; note that NeuroSynth does not distinguish activations from de-activations). **(b)** Weaker coupling impairs regions’ ability to self-organise into cognitively meaningful circuits. Percent change in the match between simulated brain activity and meta-analytic brain patterns, as a function of the global effective coupling parameter *G*, compared against the value observed for the best-fitting model (*G*=1.6). **(c)** The human structural connectome is the most appropriate network architecture for supporting co-activations for every single cognitive operation. Heatmap shows the effect of global network rewiring on the match between simulated brain activity and meta-analytic brain patterns, in terms of effect size (Hedge’s *g*). **(d)** Lesioning regions that are strongly involved with a given cognitive operation produces a strong negative impact on the brain’s ability to express the corresponding cognitive pattern. Each brain map shows, for each cortical region, the effect of lesioning that region on the match between simulated brain activity and meta-analytic brain patterns, in terms of increased or decreased correlation compared against the baseline model. **(e)** Emotion-related co-activations are consistently the most affected by disease-related changes in cortical abnormality. Heatmap shows the effect (Hedge’s *g*) on the match between simulated brain activity and meta-analytic brain patterns, upon heterogeneously increasing or decreasing intrinsic excitation of each region, according to empirical patterns of cortical thickness abnormality. **(f)** Most cognitive brain patterns can be modulated bi-directionally by appropriate engagement of different neurotransmiter systems. Heatmap shows the effect (Hedge’s *g*) on the match between simulated brain activity and meta-analytic brain patterns, upon simulating engagement of each neurotransmitter receptor. adhd = attention deficit/hyperactivity disorder; asd = autistic spectrum disorder; ocd = obsessive-compulsive disorder; ige = idiopathic generalised epilepsy; right tle = right temporal lobe epilepsy; left tle = left temporal lobe epilepsy.

We find that regional co-activations become least brain-like for values of effective coupling close to zero, as indicated by reduced prevalence of each meta-analytic map in the simulated BOLD signals (Fig. 6b). This is an important sanity check: if the effective coupling is too low, regions are effectively isolated and unable to influence each other. They cannot self-organise into cognitively meaningful patterns. We also see that at higher values of effective coupling some cognitive patterns would be expressed more strongly than in the balanced model—but this improvement would be incurred at the expense of others (Fig. 6b). Another reassuring observation about the validity of our model is that lesioning regions that are strongly involved with a given cognitive operation produces a strong negative impact on the brain’s ability to express the corresponding cognitive pattern (Fig. 6d). This result matches the expectation that regions’ ability to co-activate together depends on the anatomical connectivity between them.

We also find that the human structural connectome is the most appropriate network architecture for supporting co-activations for every single cognitive operation (Fig. 6c). Even relatively conservative rewiring schemes that preserve the weighted degree and geometric embedding lead to notable reductions in the expression of cognitive patterns in spontaneous brain activity (Fig. 6c). Most cognitive patterns are also impaired following perturbations intended to simulate different disorders—with the notable exception of ADHD (Fig. 6e). In particular, we find that emotion-related co-activations are consistently the most affected, not just for depression but across all diagnostic categories (Fig. 6e). The emotion pattern is also particularly affected by our *in vitro* pharmacology. More broadly, we see that engaging most receptors will suppress the expression of every cognitive pattern. However, three receptors exert the opposite effect, boosting the expression of most cognitive patterns (except for movement): histamine *H*_3_, dopamine *D*_2_ and serotonin 1A (Fig. 6f). As a result, the expression of 6 out of 7 cognitive brain patterns can be modulated bi-directionally by appropriate engagement of different neurotransmiter systems—fully in line with our results pertaining to global functional organisation and local dynamics (Fig. 6f).

Intriguingly, we also find that some cognitive patterns are paradoxically boosted upon lesioning specific regions. This phenomenon tends to occur for regions that are negatively associated with the patterns in question. Such a paradoxical boost may be due to the lesioned region belonging to a competing functional circuit: consistent with the existence of anti-coordinated patterns of activity in the human brain across both task and rest [129, 130].

### Convergent functional consequences of local and global macrostructural interventions

Since the functional consequences of perturbations are evaluated in terms of the same set of functional measures, we can correlate these measures against each other (with perturbations as data-points), to ask whether there are systematic associations between different aspects of brain function, as revealed by similar changes in response to perturbations. For each of the four perturbation types (local lesions, global network rewiring, neuromodulation, and cortical thickness abnormality) we find a similar division of functional properties into two broadly opposite clusters (SI Fig. S5a-d). The first, larger cluster includes most measures of hierarchy and temporal signal coordination (except intrinsic timescale) as well as redundancy and integrated information. The other, smaller cluster broadly pertains to segregation/differentiation (including modularity, anticorrelations, but also synergy, AIS and intrinsic timescale) (Fig. S5). The same two clusters are found in response to each of the four perturbations, as indicated by statistically significant correlations between each pair of correlation matrices S6). Altogether, by comparing how measures of brain function respond similarly or differently to a variety of causal manipulations, we find that they converge onto two broadly opposite categories, reflecting integration and segregation/differentiation.

## DISCUSSION

Understanding the mechanistic origins of brain function in the connectivity and physiology of the brain is a central endeavour for cognitive, systems, and computational neuroscience. Capitalising on the unparalleled experimental accessibility of computational models, here we systematically evaluated *in silico* the functional consequences of causally manipulating the brain’s network wiring and regional cyto- and chemo-architecture. Our computational structure-to-function mapping characterised the effects of each intervention against a comprehensive battery of functional measures encompassing information dynamics, spatial and temporal signal coordination, and hierarchical organisation—as well as spontaneous co-activation of distinct cognitive circuits, and thousands of univariate time-series features from the broader literature on dynamical systems[60, 61]. To catalyse additional discoveries, we make all results openly available for the neuroscience community through our interactive website (https://systematic-causal-mapping.up.railway.app/)—encompassing over 6 000 local and global functional read-outs, for each of 97 distinct causal interventions.

### Revealing the functional role of connectome architecture

Our modelling approach provides insights about the functional roles of distinct architectural features in the human structural connectome, by enabling systematic manipulations of the brain’s wiring diagram that would be simply unfeasible *in vivo*. We found that every form of global network rewiring impaired the brain’s ability to self-organise into cognitively relevant patterns, indicating that the empirical human connectome may be the most suitable wiring diagram for supporting cognition. Across both local and global manipulations, we also provide convergent evidence that the brain’s anatomical wiring diagram is finely tuned to favour the integration of information and hierarchical organisation, in support of diverse cognitive functions. The empirical human connectome was the most conducive network architecture for these fundamental properties, with virtually all local lesions inducing reductions in hierarchy and integration, and increasingly disruptive rewiring of global network topology inducing increasingly large deviations. These results are consistent with those of Fukushima and Sporns [52], who found that connectome topology and geometry both contribute to realistic integration-segregation dynamics in a Kuramoto model, but without fully accounting for them—in line with our own results.

More specifically, our global and local rewiring provide convergent evidence for prominent roles of both heterogeneous structural weights and their regional distribution. Across both univariate time-series features and global functional organisation, the most extreme differences were observed for the binary network, highlighting the crucial role of a heterogeneous weight distribution. Once heterogeneous weights are present, their distributed across regions (i.e. nodes’ weighted degree) becomes key, as indicated by the similarity of the strength-preserving rewiring to the empirical connectome in terms of functional outcomes (outperforming the various perturbations that preserve the number of connections, but without preserving their combined weight). This result is also in line with our observation that following localised structural lesions, weighted degree is the dominant predictor of the resulting influence on global functional organisation.

Many previous modelling studies have investigated the effects of lesions, rewiring, and regional heterogeneity on brain function [53, 91, 131–134]. In particular, the importance of weighted degree for predicting the functional effects of local interventions is highly consistent with reports from a diverse range of models and perturbations. These include local lesions in Kuramoto and neural mass models [135–137] as well as local stimulation in Kuramoto [138] or Wilson-Cowan models [139]. Notably, Vasa and colleagues [131] evaluated synchrony and metastability after local lesions in a Kuramoto model of coupled oscillators. On balance, both global metastability and global synchrony were decreased by lesions— as in our own model. However, lesioning some midline cortical regions could produce small increases in metastability: precisely what we also observed. This consistency of observations is reassuring especially since they were obtained using different models, hinting at broader generalisability of the present results. In addition to this computational convergence across modelling strategies and functional read-outs, the consistent observation that brain function is especially vulnerable to perturbations of high-centrality nodes is relevant for our understanding of disease [136]. Specifically, connectome hubs (the most highly connected nodes) are disproportionally implicated across a variety of brain disorders [140–142]. Among possible reasons, a high number of connections implies greater metabolic demand to sustain firing, making high-strength nodes more vulnerable to hypoxic injury [140, 143]. Altogether, when considering the distribution and placement of structural connections, our results indicate that the human connectome is the most suitable wiring scheme to promote information integration, hierarchical organisation, and the emergence of cognitively-relevant coalitions.

### Trade-offs in the functional organisation of the human brain

In addition to network wiring, interactions between regions also depend on the global level of coupling across the entire network, and on the local biophysical properties of each individual region. Crucially, our systematic assessment reveals that most measures including measures of hierarchy, integrated information and cognitive co-activations are not maximised at the value of global effective coupling where we observe the best fit between simulated and empirical brain activity. Note that although we quantified model fit using the FCD criterion (which is not one of the measures that we assessed, thereby avoiding issues of circular analysis), the observation that measures are not all maximised together is independent of which fitting criterion is chosen. This is because, regardless of which value of *G* is identified as the best fit for the real human brain, there is *no* value of *G* for which all functional features are maximised. Likewise, we found that virtually any chosen measure of brain function can be individually increased or decreased by means of *in silico* pharmacology—but not all together. For any functional measure that is increased by pharmacology or changing the effective coupling, there is at least one other whose expression is reduced.

This observation does not contradict the optimality of connectome wiring discussed above, since changes in network wiring were performed while the global effective coupling and regional excitation and inhibition were kept fixed. However, the question arises: Why does the human brain not maximise all functional properties together? A possible reason why the model’s optimal fitting point to the empirical data (*G* = 1.6) is not where most functional measures peak (*G* = 2.2), may be because other aspects of functional organisation are instead minimised around *G* = 2.2, giving rise to trade-offs between competing properties that cannot be satisfied together.

Indeed, we observed two main clusters of functional properties that behave in opposite ways in response to most causal manipulations. These clusters may be broadly described as pertaining to integration and segregation/differentiation, respectively. Many of the properties in both clusters are widely held to be beneficial for computation and cognition, such as information integration and hierarchical organisation, but also small-world architecture and information storage [54, 56, 72, 75, 144–146]. However, our model shows that such properties tend to behave in opposite fashion across most manipulations, such that when some are optimal others display their least favourable value. Furthermore, this antagonistic behaviour becomes increasingly prominent at higher levels of effective coupling, where the most extreme values are observed in each direction: the closer the model comes to optimising some properties, the further it will be from optimising others.

The best-fitting point of the model may represent a solution to this complex trade-off between competing desiderata. Indeed, this interpretation is in line with the model of Deco and colleagues [146], who found that the best fitting to empirical fMRI signals occurs at an intermediate value of the global coupling, where neither measures of integration nor segregation are maximised, but rather the two are balanced against each other. This balancing act may afford the brain the flexibility to shift between different configurations depending on environmental demands—for example thanks to endogenous engagement of different neurotransmitter systems, as suggested by our own simulations with receptor maps.

### Core-periphery organisation of brain function

Notably, the principal component of regions’ functional susceptibility to local structural lesions divides the cortex into two clear extremes. On one hand, a set of regions situated at the brain’s core (medially and ventrally), associated with internally-oriented cognitive processes (emotion, arousal, valence, memory). On the other hand, a set of regions situated at the anatomical periphery (dorsal and lateral) that pertain to more externally-oriented modes of cognition, such as attention and working memory, vision, motion, and action. Broadly, one may think of the core regions as determining what goals to pursue, and the peripheral regions as determining how to pursue them. Indeed, a similar core-periphery organisation was also found by Gollo et al [133], who related these regions with interoception and exteroception, respectively. In turn, this core-periphery distinction maps onto the two broad categories of global functional measures. Lesions to the core, value-related regions map onto measures of integration. Lesions to the peripheral regions related to perception and action map onto measures of segregation/differentiation.

We also found that the principal component of local functional susceptibility is best predicted by the principal component of NeuroSynth co-activation, far outstripping predictors derived from anatomy or even resting-state FC. In other words, regions that are involved in the same cognitive processes *in vivo*, as indicated by meta-analytic co-activation, also exhibit similar responses to perturbations in our *in silico* model. Similarly, regions’ co-susceptibility to structural lesions recapitulates their co-susceptibility to cortical thickness abnormalities associated with a variety of diagnostic categories from the ENIGMA consortium—further supported by correlations with NeuroSynth maps pertaining to mental health terms (stress, fear, anxiety, psychosis, addiction). Therefore, our computational structure-to-function mapping recapitulates both regions’ vulnerability to brain disease, and their joint involvement across different cognitive operations—even though neither of these properties was explicitly introduced into our model of local lesions. By revealing regions’ joint involvement in cognitive circuits and functional brain properties, our model is inspired by ‘lesion network mapping’ [35, 38]. NeuroSynth identifies functional circuits by finding statistical associations between cognitive constructs and cortical locations across neuroimaging studies [62, 125, 128]. In contrast, lesion network mapping identifies circuits of regions that produce common neurological symptoms when lesioned— even when the lesion locations themselves do not overlap. Our present approach provides a way to map from lesions to their predicted functional consequences.

### Brain function from dynamics to cognition

The main meaning of ‘brain function’ that we adopted in the present study reflects how brain regions act, interact, and relate to each other along multiple dimensions. To also approach ‘brain function’ from a different angle, we identified functional circuits that co-activate together to support fundamental cognitive operations— from attention to emotion and movement—as indicated by meta-analytic co-activation across thousands of neuroimaging studies [62]. We found that the brain’s ability to self-organise into functional cognitive circuits is highly sensitive to the wiring of the human structural connectome. We also found that lesioning pivotal members of a functional circuit will destabilise the entire circuit, diminishing its prevalence in spontaneous brain activity. This pattern of results from our *in silico* model is clearly consistent with empirical observations *in vivo* from neuropsychology, with its long tradition of deciphering the functional roles of different cortical regions by examining the cognitive deficits of patients with focal lesions [38, 147–155].

Overall, the results of our ‘virtual neuropsychology’ indicate that the emergence of cognitive circuits in brain activity is strongly dependent on the anatomical connectivity between regions. We also found that the expression of cognitive circuits in brain activity is compromised for nearly every *in silico* model of diagnostic categories. In particular, the meta-analytic pattern reflecting emotional processing is the most affected across disorders: possibly reflecting a shared dimension of symptomatology among many diagnostic categories. It is therefore noteworthy that the opposite result can be obtained by engaging *D*_2_, *H*_3_, and 5*HT*_1*A*_ receptors, which promote the emergence of specific cognitive patterns in spontaneous brain activity. The dopamine *D*_2_ receptor is the main target of many antipsychotic drugs [156–159]. Likewise, the serotonin 1A receptor is targeted by some atypical antipsychotics, and several medications with antidepressant and anxiolytic effects [159–163]. Based on these computational predictions, future work may investigate whether psychiatric disorders reduce the prevalence of specific cognitive patterns in patients’ spontaneous brain activity—and whether the same patterns are restored by medication acting on *D*_2_ or 5*HT*_1*A*_ receptors.

Future work will be able to expand on the present results along several directions. Models can vary greatly in their level of neurobiological detail and associated trade-off between interpretability and complexity, ranging from phenomenological Ising and Kuramoto models, to detailed Hodgkin-Huxley models. Indeed, ‘all models are wrong; the practical question is how wrong do they have to be to not be useful” (George Box, 1919-2013). Potential future extensions that do not require distinguishing excitatory and inhibitory dynamics may adopt the Hopf model to better capture oscillations. Our goal of a systematic many-to-many mapping from perturbations to function(s) imposed further computational burdens due to combinatorial explosion, requiring the use of a relatively coarse-grained model with only 68 cortical regions [77]. Use of a cortical model further ensured that the same model can integrate all our datasets, some of which are only available for the cortex. Future work could extend our results with subcortical regions; or by lesioning multiple regions at once rather than one at a time [164–167]; or implementing alternative network wiring schemes inspired by development or evolution [79, 168]; or simulating different doses of ‘virtual pharmacology’. Directionality of structural connections could also be included in future developments [169– 171]. Our model is based on a group-consensus connectome; an important development in the field will be the use of personalised models based on individual subjects’ own neuroimaging data, particularly for patients. On the functional side, alternative functional read-outs may include using the model as an artificial neural network to perform actual tasks [172–175]. Although we have shown our model’s convergence both with empirical observations and with alternative models, every model is inevitably a simplification of the underlying biology: the price for the experimental accessibility and mechanistic insight afforded by computational modelling.

### Outlook

Overall, we identified two antagonistic modes of brain function pertaining to functional integration and segregation, which are supported by ventromedial versus dorsolateral regions, and coincide with internally-oriented and externally-oriented modes of cognition. Because of this antagonistic organisation, no single intervention can increase all functional properties at once, inducing complex trade-offs. Indeed, our modelling results suggest that the wiring of the human connectome may be especially suitable to support the kinds of dynamics that promote hierarchical ignition and integration of information, while balancing other functional desiderata. Supporting this notion, most disease-related alterations of cortical thickness and alternative wiring schemes induce functional deviations from this working point. However, our *in silico* pharmacology suggests that further bi-directional tuning of each individual property may be possible by acting on the interplay of regional excitation and inhibition through drugs targeting specific neurotransmitters.

A fundamental use of computational models is to provide a mechanistic explanation for empirically observed phenomena. Since our results encompass a wide range of both structural interventions and functional read-outs, they can be used for computational ‘reverse inference’, to estimate which causal manipulations would be more or less consistent with an observed functional alteration. For example, inferring whether an individual’s level of effective coupling or regional excitation or inhibition may be higher or lower than average, based on how their profile of functional measures deviates from a normative sample. Indeed, we found that regions’ functional vulnerability to lesions *in silico* recapitulates their *in vivo* vulnerability to cortical thickness abnormalities.

More broadly, our systematic structure-to-function mapping provides a powerful engine for computational causal discovery. Our model provides unambiguous predictions about the functional consequences that we should expect from a broad range of drugs acting on different neurotransmitter systems. These extensive predictions can be tested *in vivo* through targeted pharmacological manipulations, potentially guiding the selection of appropriate pharmacological treatments. Predictions about local lesions can also be assessed in non-human animals, for example using chemo- or optogenetic activation and deactivation of specific regions, in combination with functional MRI recording [176–178]. Looking forward, our model could also find use as an *in silico* screening tool for pre-operative planning (e.g. for patients with brain tumor), to predict the functional impact of resecting tissue from different regions.

## METHODS

### Human Connectome Project data

We used resting-state functional MRI data from the 100 unrelated subjects (54 females and 46 males, mean age = 29.1 ± 3.7 years) of the HCP 900 subjects data release [179]. All HCP scanning protocols were approved by the local Institutional Review Board at Washington University in St. Louis. The diffusion-weighted imaging (DWI) acquisition protocol is covered in detail elsewhere [180]. Data were acquired using the following parameters. Structural MRI: 3D MPRAGE T1-weighted, TR = 2, 400 ms, TE = 2.14 ms, TI = 1, 000 ms, flip angle = 8^°^, FOV = 224 × 224, voxel size = 0.7 mm isotropic. Two sessions of 15-min resting-state fMRI: gradient-echo EPI, TR = 720 ms, TE = 33.1 ms, flip angle = 52^°^, FOV = 208 ×180, voxel size = 2 mm isotropic. Here, we used functional data from only the first scanning session, in LR direction.

We also used diffusion MRI (dMRI) data from the same 100 unrelated participants. The diffusion MRI scan was conducted on a Siemens 3T Skyra scanner using a 2D spin-echo single-shot multiband EPI sequence with a multi-band factor of 3 and monopolar gradient pulse. The spatial resolution was 1.25 mm isotropic. TR=5500 ms, TE=89.50ms. The b-values were 1000, 2000, and 3000 s/mm^2^. The total number of diffusion sampling directions was 90, 90, and 90 for each of the shells in addition to 6 b0 images. We used the version of the data made available in DSI Studio-compatible format at http://brain.labsolver.org/diffusion-mri-templates/hcp-842-hcp-1021 [181].

#### Functional MRI preprocessing and denoising

HCP-minimally preprocessed data [180] were used for all acquisitions. The minimal preprocessing pipeline includes bias field correction, functional realignment, motion correction, and spatial normalisation to Montreal Neurological Institute (MNI-152) standard space with 2mm isotropic resampling resolution [180].

Denoising was performed using the CONN toolbox [**?**]. The anatomical CompCor (aCompCor) method removes physiological fluctuations by extracting principal components from regions unlikely to be modulated by neural activity; these components are then included as nuisance regressors [182]. Following this approach, five principal components were extracted from white matter and cerebrospinal fluid signals (using individual tissue masks obtained from the T1-weighted structural MRI images) [183]; and regressed out from the functional data together with six individual-specific realignment parameters (three translations and three rotations) as well as their first-order temporal derivatives; followed by scrubbing of outliers identified by ART, using Ordinary Least Squares regression [183]. The de-noised BOLD signal timeseries were linearly detrended and band-pass filtered to eliminate both low-frequency drift effects and high-frequency noise, thus retaining frequencies between 0.008 and 0.09 Hz. Finally, denoised time-series were parcellated into 68 cortical regions from the Desikan-Killiany anatomical atlas [184].

#### Human structural connectome from Human Connectome Project

We adopted previously reported procedures to reconstruct the human connectome from DWI data. The minimally-preprocessed DWI HCP data [180] were corrected for eddy current and susceptibility artifact. DWI data were then reconstructed using q-space diffeomorphic reconstruction (QSDR [185]), as implemented in DSI Studio (www.dsi-studio.labsolver.org). QSDR calculates the orientational distribution of the density of diffusing water in a standard space, to conserve the diffusible spins and preserve the continuity of fiber geometry for fiber tracking. QSDR first reconstructs diffusion-weighted images in native space and computes the quantitative anisotropy (QA) in each voxel. These QA values are used to warp the brain to a template QA volume in Montreal Neurological Institute (MNI) space using a nonlinear registration algorithm implemented in the statistical parametric mapping (SPM) software. A diffusion sampling length ratio of 2.5 was used, and the output resolution was 1 mm. A modified FACT algorithm [186] was then used to perform deterministic fiber tracking on the reconstructed data, with the following parameters [187]: angular cutoff of 55°, step size of 1.0 mm, minimum length of 10 mm, maximum length of 400 mm, spin density function smoothing of 0.0, and a QA threshold determined by DWI signal in the cerebrospinal fluid. Each of the streamlines generated was automatically screened for its termination location. A white matter mask was created by applying DSI Studio’s default anisotropy threshold (0.6 Otsu’s threshold) to the spin distribution function’s anisotropy values. The mask was used to eliminate streamlines with premature termination in the white matter region. Deterministic fiber tracking was performed until 1, 000, 000 streamlines were reconstructed for each individual.

For each individual, their structural connectome was reconstructed by drawing an edge between each pair of regions *i* and *j* from the Desikan-Killiany cortical atlas [184] if there were white matter tracts connecting the corresponding brain regions end-to-end; edge weights were quantified as the number of streamlines connecting each pair of regions, normalised by ROI distance and size.

A group-consensus matrix *A* across participants was then obtained using the distance-dependent procedure of Betzel and colleagues, to mitigate concerns about inconsistencies in reconstruction of individual participants’ structural connectomes [188]. This approach seeks to preserve both the edge density and the prevalence and length distribution of inter- and intra-hemispheric edge length distribution of individual participants’ connectomes, and it is designed to produce a representative connectome [188, 189]. This procedure produces a binary consensus network indicating which edges to preserve. The final edge density was 27%. The weight of each non-zero edge is then computed as the mean of the corresponding non-zero edges across participants.

### Whole-brain computational model

Macroscale whole-brain computational models represent regional activity in terms of two key ingredients: (i) a biophysical model of each region’s local dynamics; and (ii) inter-regional anatomical connectivity. Thus, such *in silico* models provide a well-suited tool to investigate how the structural connectivity of the brain shapes the corresponding macroscale neural dynamics [18, 43, 44, 63]. In particular, the Dynamic Mean Field (DMF) model employed here simulates each region (defined via a species-specific brain parcellation scheme) as a macroscopic neural field comprising mutually coupled excitatory and inhibitory populations, providing a neuro-biologically plausible account of regional neuronal firing rate. Specifically, the model simulates local biophysical dynamics of excitatory (NMDA) and inhibitory (GABA) neuronal populations, interacting over long-range neuroanatomical connections. Here, we used a consensus human connectome reconstructed from *in vivo* diffusion MRI tractography.

The following differential equations therefore govern the model’s behaviour:

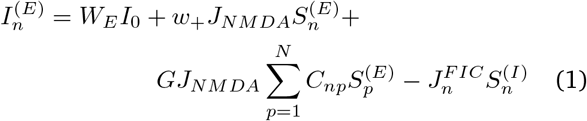

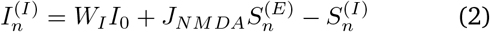

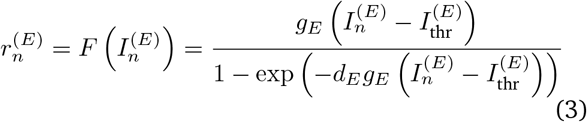

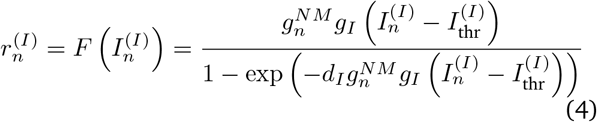

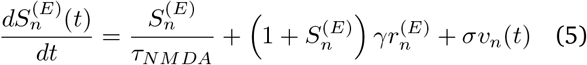

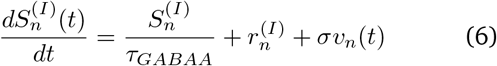

Following previous work [18, 63, 64, 91], all parameters were set as in Wong and Wang [] [79]. for each excitatory (E) and inhibitory (I) neural mass, the quantities 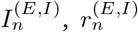, and 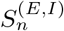 represent its total input current (nA), firing rate (Hz), and synaptic gating variable, respectively. The function *F* (·) is the transfer function (or F–I curve), representing the non-linear relationship between the input current and the output firing rate of a neural population. Finally, 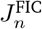 is the local feed-back inhibitory control of region *n*, which is optimized to keep its average firing rate at approximately 3 Hz, and *ν*_*n*_ is uncorrelated Gaussian noise injected to region *n*. The model’s fixed parameters are reported in 1. Due to its multi-platform compatibility, low memory usage, and high speed, we used the recently developed and publicly available FastDMF library [64]. The code used to run all the simulations in this study was written in optimised C++ using the high-performance library *Eigen*. The C++ core of the code, together with Python and Octave/Matlab interfaces is publicly available as FastDMF and maintained at http://www.gitlab.com/concog/fastdmf [64].

#### Model fitting and global effective coupling

Following [18, 64], all but one of the DMF model parameters are set as per [190], leaving only one free parameter, known as ‘global coupling’ and denoted by *G*, which controls the overall effectiveness of signal transmission between brain regions (conductivity of the white matter fibers is assumed to be constant across the brain). We first tune *G* to match the high-quality Human Connectome Project functional MRI dataset [179]. We set the model to have the same TR as the HCP fMRI data (0.72s), with data filtered in the same frequency band (0.008-0.09 Hz). Then we use a Bayesian optimiser to identify the *G* value that maximises the fit between model and empirical HCP fMRI data [64]. Goodness of fit was quantified as the similarity between the data’s and the model’s functional connectivity dynamics (FCD), computed as follows. First, we obtained Pearson correlation matrices between regional fMRI signal time-series, computed within a sliding window of 30 TRs with increments of 3 TRs [18, 91]. Subsequently, the resulting matrices of functional connectivity at times *t*_*i*_ and *t*_*j*_ were themselves correlated, for each pair of timepoints *t*_*i*_ and *t*_*j*_, thereby obtaining an FCD matrix of time-versus-time correlations. Thus, each entry in the FCD matrix represents the similarity between functional connectivity patterns at different points in time. The best-fitting value of the *G* parameter is identified as the one that minimises the Kolmogorov-Smirnov distance between the histograms of empirical (group-wise) and simulated FCD values (obtained from the upper triangular FCD matrix) [18, 91]. Use of the FCD is well established as fitting target for the DMF model [18, 64, 91], and we note that this measure is not one of the measures that we investigate in the present study, thereby avoiding the issue of circularity.

### Local lesions

Local perturbations were implemented at the level of each region. For each region in turn (considering both left and right hemispheres together), a locally perturbed connectome was obtained by setting all of that region’s connections to 0.1× its original value, thereby preserving the topology of the connectome and its binary degree (and ensuring that it does not become disconnected), while diminishing the region’s ability to exchange signals with the rest of the brain. For each perturbation, 30 instances of the model were run, with random initial conditions.

### Global rewiring schemes for the connectome

Global perturbations were implemented as rewirings of the connectome. The total number of edges in the network was always preserved, and so was the distribution of edge weights (except for the binary connectome described below). Additional constraints and specifics of each type of rewiring are provided below. For each global perturbation, 30 instances of the model were run, with random initial conditions.

#### Binarisation

The first perturbation that we consider is binarisation, whereby all edges’ weights are set to unity, without otherwise altering the network.

#### Random rewiring

The network’s edges were randomly rewired. The resulting network has the same edge density and distribution of edge weights, but does not preserve the degree sequence.

#### Modular network

To rewire the connectome into a random hierarchical modular network, we began by generating a binary random network with a specified number of densely connected modules (here, 4) linked together by evenly distributed remaining random connections, having the same number of nodes and the same edge density as the original structural connectome. Because this method creates a directed network, the matrix was symmetrized by selecting the upper triangle of the resultant matrix and using these connections to create an undirected, symmetric binary network. Weights were then assigned at random to the non-zero edges by drawing them without replacement from the distribution of empirical edge weights, while preserving network symmetry.

#### Rewiring into a lattice

The network was rewired to have a lattice topology, while preserving the degree sequence and the distribution of edge weights [117].

#### Degree-preserving random rewiring

The well-known Maslov-Sneppen degree-preserving rewired network swaps edges so as to randomise the topology while preserving the exact binary degree of each node (degree sequence), and the overall distribution of edge weights [116].

#### Geometry-preserving rewiring

In addition to preserving exactly the same degree sequence and exactly the same edge weight distribution as the original network, the cost-preserving model also approximately preserves the original network’s edge length distribution (based on Euclidean distance between regions), and the weight-length relationship [118].

#### Degree- and strength-preserving rewiring

The last null model employed simulated annealing to preserve the original network’s strength sequence (sum of edge weights incident to each node). Simulated annealing is a stochastic search algorithm that approximates the global minimum of a given function [191] using the Metropolis technique [192], controlled by the temperature parameter *T*. A high temperature regime allows the exploration of costly system configurations, whereas fine-tuned adjustments with smaller effects on the system cost are provided at lower temperatures. Initially, the simulated annealing algorithm is set at a high temperature, preventing the process from getting stuck in local minima. Throughout the procedure, the system is slowly cooled down while descending along the optimization landscape, yielding increasingly limited uphill rearrangements. Here, we minimize the cost function *E* defined as the sum of squares between the strength sequence vectors of the original and the randomized networks. To optimize this function, weights were randomly permuted among edges. Reconfigurations were only accepted if they lowered the cost of the system or met the probabilistic Metropolis acceptance criterion: *r < exp*(− (*E*^*′*^− *E*)*/T*), where *r* ∼*U* (0, 1). The annealing schedule consisted of 100 stages of 10000 permutations with an initial temperature of 1000, halved at each stage.

### Virtual patients with cortical thickness abnormality maps from the ENIGMA database

Spatial maps of case-versus-control cortical thickness were obtained by including all the neurological, neurodevelopmental, and psychiatric diagnostic categories available from the ENIGMA (Enhancing Neuroimaging Genetics through Meta-Analysis) consortium [66, 67] and the Enigma toolbox (https://github.com/MICA-MNI/ENIGMA) [65] and recent related publications (https://github.com/netneurolab/hansen_crossdisorder_vulnerability) [100], except for obesity and schizotypy. This resulted in a total of 11 maps, pertaining to 22q11.2 deletion syndrome [193], attention-deficit/hyperactivity disorder [194], autism spectrum disorder [195], idiopathic generalized epilepsy [196], right temporal lobe epilepsy [196], left temporal lobe epilepsy [196], depression [197], obsessive-compulsive disorder [198], schizophrenia [199], bipolar disorder [200], and Parkinson’s disease [201]. The ENIGMA consortium is a data-sharing initiative that relies on standardized image acquisition and processing pipelines, such that cortical thickness maps are comparable [67]. Altogether, over 17 000 patients were scanned across the eleven diagnostic categories, against almost 22 000 controls. The values for each map are effect sizes (Cohen’s *d*) of cortical thickness in patient populations versus healthy controls. Imaging and processing protocols can be found at http://enigma.ini.usc.edu/protocols/. The largest increase in cortical thickness across all diagnostic categories is 0.87 (expressed in terms of Cohen’s *d*), whereas the largest decrease is 0.59.

For each diagnostic category, we modulate the regional level of intrinsic excitability in the model according to the regional pattern of increases or decreases in cortical thickness associated with that condition, following the rationale of [102]. Namely, when atrophy of cortical thickness is observed, we model it as reduced intrinsic excitation; and when an increase in thickness is observed, we increase the intrinsic excitation in the corresponding region. Concretely, for each of the 11 ENIGMA maps we modulate the excitatory scaling factor *W*_*E*_ of each region, which in the baseline DMF is uniformly set to *W*_*E*_ = 1 for all regions, by adding to it the corresponding region’s value of cortical thickness change (a positive number for thickness increase, and a negative number for thinning). For each map, this results in a DMF model with regionally heterogeneous intrinsic excitation.

### Virtual pharmacology with receptor maps from Positron Emission Tomography

Receptor densities were estimated using PET tracer studies for a total of 15 receptors, across 8 neurotransmitter systems, recently made available by Hansen and colleagues at https://github.com/netneurolab/hansen_receptors [2]. These include dopamine (D_1_ [202], D_2_ [203–206], serotonin (5-HT_1A_ [207], 5-HT_1_B [207– 212], 5-HT_2A_ [213], 5-HT_4_ [213], 5-HT_6_ [214, 215], acetylcholine (*α*_4_*β*_2_ [216, 217], M_1_ [218], glutamate (mGluR_5_ [219, 220] NMDA [221, 222], GABA (GABA_A_ [223])), histamine (H_3_ [224]), cannabinoid (CB_1_ [225– 228]) and opioid (MOR [229]). Volumetric PET images were registered to the MNI-ICBM 152 nonlinear 2009 (version c, asymmetric) template, averaged across participants within each study, then parcellated and receptors with more than one mean image of the same tracer (5-HT_1_B, D_2_) were combined using a weighted average [2]. Finally, each map was normalised between 0 and 1. Following the approach of [18], the effect of engaging an excitatory receptor is modelled by increasing the excitatory gain parameter of each region, according to that region’s receptor density (normalised between 0 and 1). Likewise, the effect of engaging an inhibitory receptor is modelled by increasing the inhibitory gain parameter of each region, according to that region’s receptor density (normalised between 0 and 1)—as per [91]. Importantly, these previous studies had the goal of investigating whether tuning regional excitatory or inhibitory gain according to specific receptor maps would increase the goodness-of-fit to specific empirical data (obtained under the effects of LSD and anaesthesia, respectively) over and above the fit to placebo/baseline data [18, 91]. Therefore, they explicitly optimised an additional global gain-scaling parameter, whose value was allowed to change for each map until the fit to the drug data was maximised for that map. Here, our goal is different. We are not trying to use the receptor maps to maximise the fit to some specific external dataset; rather, we are interested in evaluating the maps’ differential effects on brain function. By using the receptor maps as inputs to the same model, with everything else kept fixed, we can unambiguously conclude that any resulting functional difference is attributable to the map’s identity. Therefore, we do not include a separate global gain scaling parameter to be optimised for each map. Rather, the original excitatory (resp., inhibitory) gain parameter of each region, which is uniformly set to 1 in the baseline model, is then increased by that region’s normalised receptor density value (which ranges between 0 and 1). Future work may expand on this approach by evaluating different scaling values for each receptor map, thereby simulating different doses of ‘virtual pharmacology’.

### Information dynamics

The framework of integrated information decomposition (ΦID) decomposes information flow into interpretable, disjoint parts, providing a taxonomy of “modes” of information dynamics in multivariate dynamical systems. In this section we provide a brief description of (ΦID).

Integrated Information Decomposition (ΦID [68]) is a generalisation of Partial Information Decomposition (PID; [230]), a technique to decompose the information that multiple *source* variable have about a single *target* variable. ΦID generalises PID by extending the formalism to systems with multiple sources *and* multiple targets. This makes ΦID particularly well-suited for the study of multivariate stochastic processes, where the state of each process at time *t* can be seen as the sources of information and the states at *t*^′^ *> t* as targets. In such scenarios, ΦID describes how information is carried by the variables in the system over time.

For the case of two sources of information *X, Y* and one target *Z*, PID divides their information into four “information atoms:” redundancy (Red), synergy (Syn), and unique of the first and second sources (`Un`^X^ and `Un`^Y^). The ΦID of two timeseries (…, *X*_*t*_, *X*_*t*+1_, …) and (…, *Y*_*t*_, *Y*_*t*+1_, …) between timepoints *t* and *t*^′^ considers two sources of information *X*_*t*_, *Y*_*t*_ and two targets *X*_*t*_^*′*^, *Y*_*t*_^*′*^, and their information is divided into 16 atoms that comprise all possible ways in which the 4 atoms of traditional PID can evolve in time. For example, ΦID’s atom is the information transferred from *X*_*t*_ to *Y*_*t*_^*′*^ — i.e. information that was uniquely present in *X* at time *t*, and is present only in *Y* at time *t*^′^. Similarly, is a case of information *erasure*: it was present in both *X* and *Y* (i.e., redundant) at time *t*, and is present only *Y* (and hence not anymore in *X*) at time *t*^′^. As another example, is information that is persistently redundant: it was present redundantly in both *X* and *Y* at time *t*, and stays redundant at time *t*^′^. Similarly, was carried synergistically by *X* and *Y* at time *t*, and is persistently synergistic at time *t*^′^. More details about ΦID can be found in Ref. [68].

Given a time series dataset from a bivariate stochastic process, it is possible to compute all atoms in its ΦID decomposition given a suitable form of the joint probability density function *p*(*X*_*t*_, *Y*_*t*_, *X*_*t*_^*′*^, *Y*_*t*_^*′*^). In this work, we take the simple yet empirically useful assumption that this probability density is a multivariate Gaussian distribution (which is an accurate description of functional brain activity data [231]).

Under these assumptions, the mutual information between two random variables *U* and *V* can be computed from their joint covariance matrix as

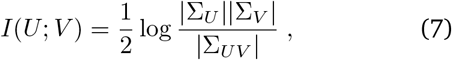

where Σ_*U*_ (resp. Σ_*V*_) is the covariance matrix of *U* (resp. *V*), and Σ_*UV*_ is the joint covariance matrix of *U* and *V*. With this expression, we can compute the mutual information between any pair of (sets of) variables in the data. As an example, the time-delayed mutual information of the joint stochastic process is given by

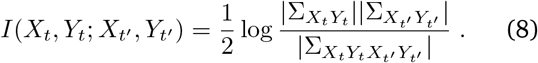

Equipped with this expression for mutual information, we can now tackle the problem of computing the atoms of PID and ΦID. Under reasonable assumptions [232], the PID of Gaussian variables reduces to the so-called Minimum Mutual Information (MMI) approach, under which redundancy is calculated as

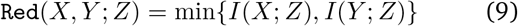

for any target variable *Z*. Similarly, in the case of ΦID, this redundancy measure can be extended to yield the double-redundancy atom,

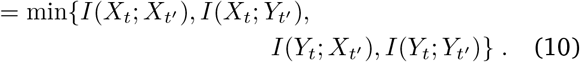

Finally, after calculating with Eq. (10), PID redundancy with Eq. (9), and mutual information with Eq. (7) for all relevant system subsets, the other 15 information atoms can be calculated as the solution of a simple linear system of equations, as described in Ref. [68]. Overlapping segments of the functional time series with one time step (TR) delay were used to define the past and future states. We calculated ΦID for every pair of the original functional time series using time-delayed mutual information (mutual information between the past and future states) under the Gaussian assumption for continuous variables, and the MMI definition of redundancy. An open-source implementation can be found at https://github.com/ Imperial-MIND-lab/integrated-info-decomp.

All information-theoretic measures that we include here can be obtained from Information Decomposition. Specifically, we consider the persistent redundancy () and the persistent synergy (), as well as the additional information-dynamic phenomena of Integrated Information, Transfer Entropy, and Active Information Storage, described below.

#### Integrated Information

Integrated Information quantifies the co-existence of integration and differentiation in a system. An initial formulation for dynamical systems was introduced by Tononi and Balduzzi [233] and adapted for use in empirical data by [234], corresponding to:

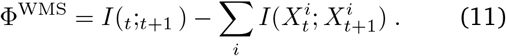

This formulation is intended to reflect the extent to which the information contained within the whole system (*I*(_*t*_;_*t*+1_)) exceeds the information contained within the sum of the parts 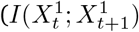 and 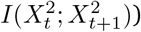. In other words, it indicates whether ‘the whole is greater than the sum’.

However, this initial formulation has well-known shortcomings, including the fact that it can take on negative values. Through the framework of Integrated Information Decomposition we can decompose the constituent elements of Integrated Information and show that it is composed of different information atoms: it contains all the synergistic information in the system, the unique information transferred from X to Y and vice versa, and, importantly, a negative redundancy contribution [68]. This explains why Φ can be negative, which will occur in redundancy-dominated systems. This subtraction of the double-redundancy occurs because redundancy is (by definition) present in each of the parts of the system, Therefore, when computing the ’sum of the parts’ to subtract from the whole in order to obtain Integrated Information as “whole minus sum of the parts”, the redundant information is double-counted and subtracted twice. Thus, the original formulation of Balduzzi and Tononi is actually not the difference between whole and sum of the parts, but the balance between Integrated Information and redundancy. The proper whole-minus-sum measure of Integrated Information can be obtained by ‘adding back’ the redundancy:

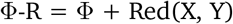

[68]. This is the measure of Integrated Information that we use here. Note that several variants of Φ have been proposed over the years, including the original formulation of Balduzzi and Tononi [233], other formulations based on causal perturbation [235] (see [236, 237] for comprehensive reviews).

#### Active information storage

Active Information Storage (AIS) [238, 239], quantifies how much information from a variable’s past remains available in its future (meaning that it has been stored in the variable’s past); formally, it is defined as the mutual information between the present of one variable, 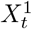, and its own future, 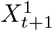 (equivalently, the time-delayed mutual information of an individual part of the system). AIS can be obtained from information decomposition in terms of redundancy and unique information, but now taking into account *both* past and future:

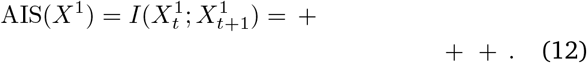

#### Information transfer

Information-theoretic measures of transfer were introduced by Schreiber [240], Massey et al. [241]. In particular, Schreiber’s Transfer Entropy is one of the most widely used measures of information transfer: considering X as source and Y as target, *TE*(*X, Y*) quantifies the information about Y’s future that is provided by X’s past, over and above what is already provided by Y’s own past. Hence, TE corresponds to:

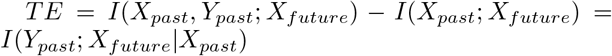

By explicitly excluding any information in the future of Y that was already present in Y’s own past, TE ensures that information storage in Y is not mistaken as having been transferred [238, 242]. Note that this is the same rationale underpinning the Wiener-Granger causality measure of statistical causal discovery [243, 244]. Indeed, TE and Granger Causality are identical in linear systems [245]. TE can also be obtained in terms of information decomposition, and this is what we did here, as follows:

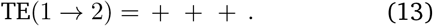

### Temporal signal coordination

#### Synchrony and Metastability

Metastability was quantified using a widely-used proxy signature, the standard deviation of the Kuramoto Order Parameter across time (referred to as *std*(*KOP*) for brevity) [58, 59, 85].

In turn, the Kuramoto Order Parameter is defined by the following equation:

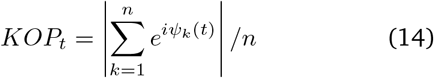

where *ψ*_*k*_(*t*) is the instantaneous phase of each bandpass-filtered BOLD signal at node *k* (note that the symbol *ϕ* is often used in equations for the KOP instead of *ψ*, but here we avoid using *ϕ* to avoid confusion with the measure of Integrated Information [246]).

Following Shanahan [59]: we computed the instantaneous phase *ψ*_*k*_(*t*) of each bandpass-filtered signal *k* using the Hilbert transform. The Hilbert transform yields the associated analytical signals. The analytic signal represents a narrowband signal, *s*(*t*), in the time domain as a rotating vector with an instantaneous phase, *ψ*(*t*), and an instantaneous amplitude, *A*(*t*). Thus, *s*(*t*) = *A*(*t*) cos(*ψ*(*t*)). The phase and the amplitude are given by the argument and the modulus, respectively, of the complex signal *z*(*t*), given by *z*(*t*) = *s*(*t*) + *iH*[*s*(*t*)], where *i* is the imaginary unit and *H*[*s*(*t*)] is the Hilbert transform of *s*(*t*).

At each point in time, the Kuramoto order parameter measures the global level of synchronization across these oscillating signals. Under complete independence, the phases are uniformly distributed and thus *KOP* is nearly zero, whereas *KOP* = 1 if all phases are equal (full synchronization).

The variability (standard deviation) of the *KOP* over time is commonly used as a signature of metastability, first proposed by Shanahan [59]. If *std*(*KOP*) is high, it indicates that the system alternates between high and low synchronisation, thereby combining tendencies for integration (high synchrony) and segregation (low synchrony). For each individual (and simulation), we obtain the *std*(*KOP*) signature of metastability, as well as the maximum observed value of global synchrony.

#### Intrinsic timescale

We estimated the intrinsic timescale of each brain region by computing the autocorrelation function of each brain region. Following [70, 247], we used least-square fitting to fit a nonlinear exponential decay function to the empirically estimated autocorrelation function. The exponential decay function was fitted as a function of the time lag *k*Δ between time bins, where *k* = | *i* −*j*|, and obeyed the equation:

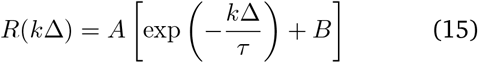

where *A* corresponds to a scaling factor, *B* reflects the offset for contribution of timescales longer than the observation window, and they obey parameter bounds:

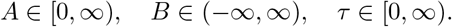

The intrinsic timescale (i.e., rate of decay) is then given by *τ*.

#### Temporal irreversibility

We estimate pairwise interactions between brain regions by computing time-shifted correlations between both the forward and reversed fMRI BOLD time series of any two regions. This method effectively quantifies the asymmetry in interactions between region pairs, thereby indicating how one region influences another. This approach is inspired by thermodynamics, where the breaking of detailed balance is associated with non-reversibility, often referred to as the ‘arrow of time’ [57]. Irreversibility is captured as the difference between the time-shifted correlations of forward and reverse time series.

To illustrate, consider the detection of irreversibility between two time series, *x*(*t*) and *y*(*t*). The temporal dependency between *x*(*t*) and *y*(*t*) is measured using time-shifted correlations. For forward evolution, the time-shifted correlation is given by:

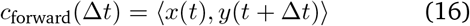

Similarly, we create a reversed version of *x*(*t*) (or *y*(*t*)), denoted *x*^*r*^(*t*) (or *y*^*r*^(*t*)), by inverting the time sequence. The time-shifted correlation for the reversed evolution is then:

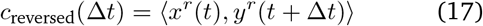

The pairwise level of irreversibility, representing the degree of temporal asymmetry or the arrow of time, is quantified as the absolute difference between the forward and reversed time-shifted correlations at a given shift Δ*t* = *T* (here, for consistency across species we set *T* = 1 TR):

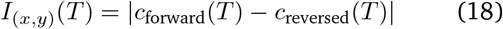

To compute the whole-brain level of irreversibility, we define forward and reversal matrices of time-shifted correlations for the forward version *x*_*n*_ and the reversed backward version 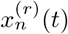 of a multidimensional time series, where the subscript *n* represents different brain regions. These matrices capture the functional causal dependencies between the variables in the forward and artificially generated reversed time series, respectively. The forward and reversed matrices are expressed as:

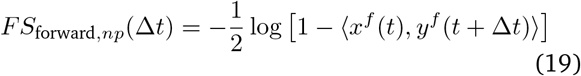

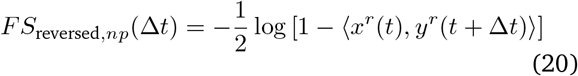

These matrices, representing the functional temporal dependencies, are based on the mutual information derived from the respective time-shifted correlations.

*FS*_diff,*np*_ is a matrix containing the squared differences of the elements between the forward and reversed matrices:

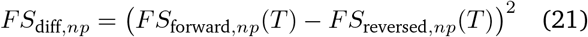

where each element reflects the irreversibility level for that region pair. The mean of this matrix provides a value of whole-brain irreversibility [57].

## Spatial signal coordination

### Prevalence of anticorrelations

We measured the prevalence of anticorrelations as the proportion of negative edges, out of the total number of edges in the functional connectivity matrix.

#### Modularity

We constructed a network from the functional connectivity, with nodes being regions and edges being the functional connectivity between them (excluding negative ones). We then applied Newman’s algorithm, which quantifies modularity as the degree to which the network can be subdivided into nonoverlapping groups of nodes in a way that maximizes the number of within-group edges, and minimizes the number of between-group edges [248].

#### Small-World Propensity

A small-world network combines the presence of tightly interconnected clusters (characterising lattice networks, and theorised to support specialised processing) with a short characteristic path length (a key feature of random network, facilitating integration between different clusters) [144, 249]. Thus, small-worldness represents a mark of optimal balance between global and local processing. We adopted the measure of small-world propensity recently developed by Muldoon et al. [72], which provides a theoretically principled way to quantify and compare the extent that different networks exhibit small-world structure while accounting for network density. The small-world propensity, *SMP*, is designed to quantify the extent that a network displays small-world organisation by taking into account the deviation of the network’s empirically observed clustering coefficient, *C*_obs_, and characteristic path length, *L*_obs_, from equivalent lattice (*C*_latt_, *L*_latt_) and random (*C*_rand_, *L*_rand_) networks:

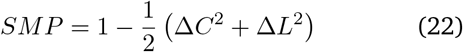

where

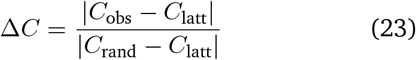

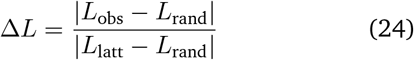

Thus, Δ*C* and Δ*L* quantify the fractional deviation of the empirically observed clustering coefficient and characteristic path length from the corresponding null models according to the definition of a small-world network: namely, a lattice network for the clustering coefficient, and a random network for the characteristic path length [72].

Following Muldoon et al. [72], we further bound both measures of fractional deviation Δ*C* or Δ*L* between 0 and 1 (to account for the possibility of empirical networks exceeding the corresponding null models), by setting negative values of Δ*C* or Δ*L* to 0, and values that exceed unity to be exactly 1. In turn, this ensures that the resulting values of small-world propensity will also be bounded between 0 and 1.

Small-world propensity is then interpreted as follows: both a large Δ*C* or Δ*L* would indicate a large deviation of the network’s properties from the corresponding properties that define small-world organisation. Thus, large Δ*C* or Δ*L* would lead to the measure of small-world propensity becoming closer to zero. Conversely, if a network exhibits both the high clustering coefficient of a lattice and the low path length of a random network (thereby satisfying both requirements of the small-world network definition), then it will have low Δ*C* and low Δ*L*, and the small-world propensity as a whole will be closer to 1. Hence, higher small-world propensity intuitively indicates better adherence to the requirements of a small-world network Muldoon et al. [72].

### Hierarchical organisation

#### Spatiotemporal Hierarchy from Intrinsic-driven Ignition

‘Intrinsic-driven ignition’ quantifies the extent to which spontaneously occurring (‘intrinsic’) local events elicit whole-brain activation (‘ignition’) [75, 122]. To compute intrinsic-driven ignition, the timeseries are transformed into z-scores, and subsequently thresholded to obtain a binary sequence based on the combined mean and standard deviation of the regional transformed signal, such that *σ*(*t*) = 1 if *z*(*t*) *>* 1 and is crossing the threshold from below, indicating that a local event has been triggered; otherwise, *σ*(*t*) = 0. Note that the threshold of 1 standard deviation for triggering an event is chosen for consistency with previous work, but it has been demonstrated that the results of this procedure are robust to the specific threshold chosen [250]. Subsequently, for each brain region, when that region triggers a local event (*σ*(*t*) = 1), the resulting global ignition is computed within a time-window of 4 TRs, in line with previous work. An *NxN* binary matrix *M* is then constructed, indicating whether in the period of time under consideration two regions *i* and *j* both triggered an event (*M*_*ij*_ = 1). Intrinsic-driven ignition (IDI) is given by the size of the largest connected component of this binary matrix *M*, quantifying the breadth of the global ignition generated by the driver region at time *t*. To obtain a measure of spatio-temporal hierarchy of local-global integration, each region’s IDI values are averaged over time, and the variability (standard deviation) across regions is then computed. Consequently, higher standard deviation reflects more heterogeneity across brain regions with respect to their capability to induce ignition, which suggests in turn a more elaborate hierarchical organisation between them [75, 122].

#### Hierarchical Integration

A distinct but complementary perspective on hierarchical brain function is in terms of nested organisation. Such a perspective can be obtained from studying the brain’s eigenmodes, and how the latter support the balance between integration and segregation across scales [56, 95]. Being a symmetric matrix, the functional connectivity can be decomposed as *FC* = *U* Λ*U* ^*T*^, where *U* is an orthogonal matrix whose columns are eigenvectors (eigenmodes) of FC, and Λ is a diagonal matrix whose entries are the eigenvalues of FC. Each eigenmode of functional connectivity identifies a distinct pattern of regions that are jointly activated (same sign) or alternate (opposite sign). Therefore, hierarchical modules can be identified based on the concordance or discordance of signs between regions across eigenmodes, progressively partitioning the FC into a larger number of modules and submodules, up to the level where each module coincides with a single region, indicative of completely segregated activity. The first eigenmode has the same sign throughout the entire cortex, reflecting global integration. At the next level, two partitions can be detected based on their different signs in the second eigenmode, and each in turn is subdivided at the following level of the hierarchy (i.e., from the third eigenmode) on the basis of regional signs. Thus, segregated modules at one level of the hierarchy can become integrated by being part of the same superordinate module (note that this hierarchical modularity based on eigenmodes is not equivalent to the clustering or modularity maximisation methods [56, 251]. During this nested partitioning process, we obtain the module number *M*_*i*_, (*i* = 1…*N*) and the modular size *m*_*j*_, (*i* = 1..*M*_*i*_) at each level.

Each level *i* of the hierarchy is characterised by two quantities: the number of modules *M*_*i*_ into which the cortex is divided, and the covariance explained by the corresponding eigenmode (given by its squared eigenvalue 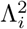). However, the number of modules alone may not properly describe the picture of nested segregation and integration because the size of modules may be heterogeneous. The correction factor was calculated as *p*_*i*_ = Σ _*j*_ |*m*_*j*_ − *N/M*_*i*_| */N* which reflects the deviation from the optimized modular size in the ith level. Thus, the correction effect is stronger for a larger deviation of modular size from homogeneity [56].

Since the first eigenmode encompasses the entire cortex into one global module, the corresponding eigenvalue 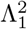 quantifies the overall contribution of global integration, 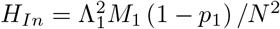 [56].

#### Functional processing hierarchy

A third way to conceptualise ‘hierarchical organisation’ in the brain is in terms of the distance between opposite ends of the information processing stream. Recently, this hierarchy has been quantified in terms of gradients of functional connectivity [55, 95, 96, 98, 99, 252, 253]. Here, we delineate gradients using the common nonlinear dimensionality reduction technique known as diffusion map embedding implemented in the BrainSpace toolbox https://github.com/MICA-MNI/BrainSpace [252], with default parameters for kernel, similarity metric, and sparsity (see below).

Following previous work, each functional connectivity matrix was thresholded row-wise to achieve 90% sparsity, retaining only the strongest connections in each row [95, 98]. The cosine similarity matrix was calculated on the thresholded z-matrix to generate a similarity matrix reflecting the similarity in whole-brain connectivity patterns between vertices. While the FC matrix reflects how similar each pair of regions are in terms of their temporal cofluctuations, this similarity matrix reflects how similar two regions are in terms of their patterns of FC. The relative influence of density of sampling points on the manifold is controlled by an additional parameter in the range of 0 to 1, which for diffusion map embedding is set to 0.5 to provide a balance between local and global contributions to the embedding space estimation [254]. The high-dimensional similarity matrix is treated as a graph, with ‘connections’ (entries of the similarity matrix) reflecting the similarity between the regional patterns of FC. The technique estimates a low-dimensional set of embedding components (gradients); in this low-dimensional space, proximity reflects similarity of the patterns of FC: regions with similar FC patterns (which are strongly connected in the network) are placed close to each other, and regions with low similarity are placed far apart. In this way, each gradient represents one dimension of covariance in the inter-regional similarity between FC patterns, with a small number of gradients capturing most of the dimensions of inter-regional similarity [55, 98, 252].

In the embedding space, each gradient can be understood to be “anchored” at regions that have the strongest values for that gradient, suggesting that this particular embedding dimension captures their similarity profiles of FC well. In contrast, regions that are close to the origin (i.e. have a low absolute value for a particular gradient) mean that they are only minimally similar to the ‘anchor points’ of that gradient, which overall does not strongly capture their FC similarity profile well overall [55, 98, 252, 253].

Therefore, the more different the extremes of a gradient differ along the axis of the gradient, the more the differentiation between regions is being captured by that gradient. To quantify this formally, we calculated the difference between the maximum and minimum values of each scan along the first gradient [95, 96, 98] (which mathematically captures most of the variation in FC profiles within each scan) and compared these differences across conditions.

### Dynamical features

To extract the dynamical phenotype of each brain region, we performed time-series feature extraction using the *catch22* toolbox Lubba et al. [124]. This set of 22 univariate dynamical features was identified as capturing a diverse range of interpretable time-series properties (such as autocorrelation, statistical properties of the distribution) that well summarise the broader literature on dynamical systems, exhibiting strong classification performance across a broad collection of time-series problems Lubba et al. [124]. Note that one of these 22 features, *CO_HistogramAMI_even_2_5*, was excluded by pre-filtering in empirical data and therefore is not included among the features that we extract, leaving 21 (2). Each dynamical feature was extracted for each region, and we then obtained a single whole-brain value by averaging across all regions.

#### Massive temporal feature extraction using hctsa

We also performed massive time-series feature extraction using the *highly comparative time-series analysis* toolbox, hctsa [60, 61], of which *catch22* is a representative subset. For each region, the hctsa toolbox extracted >7 300 univariate dynamical features, derived from diverse fields including neuroscience, physics, ecology and economics [60, 61]. Features range from basic statistics of the distribution of time-points, linear correlations among time-points, and stationarity, to measures of entropy, time-delay embeddings, and signal complexity, among others. Each individual feature is the implementation of a computation (termed ‘master operation’) on the input time-series, using specific parameters. For example, sample entropy (SampEn) computes the probability that similar sequences of observations in a time-series will remain similar as their size increases. Its computation therefore requires a threshold *r* for deciding when two sequences will be considered similar; and an embedding dimension *m* that determines the size of the sequences. Multiple individual features are obtained from this master operation as different combinations of *r* and *m*.

We performed an initial pre-filtering, and did not compute any features that had returned *NaN* across all regions, or that had displayed no variance, in an independent dataset of human functional MRI (Human Connectome Project; [179]). The values of different features can vary across several orders of magnitude. Unless otherwise specified, here we do not normalise features across regions, since this could obscure differences across conditions. Instead, we use effect sizes computed on the original features (see below). Despite our initial pre-filtering, not all of the remaining time-series features could be extracted successfully. We therefore performed an additional post-filtering. Features that failed to be extracted were excluded.

#### Dynamical profile similarity

To quantify the similarity between the temporal profiles of each pair of brain regions, each dynamical feature is z-scored across regions. The vectors of z-scored regional features are then correlated for each pair of brain regions, producing a matrix of ‘dynamical profile similarity’ (also termed ‘temporal profile similarity’ [22]) that represents the strength of the similarity of the local dynamical fingerprints of brain areas. The coupling between DPS and functional connectivity is in turn obtained by correlating the vectorised DPS and FC matrices. FC is computed as the zero-lag correlation between pairs of regional time-series.

### NeuroSynth meta-analytic maps

Continuous measures of the association between voxels and cognitive categories were obtained from NeuroSynth, an automated term-based meta-analytic tool that synthesizes results from more than 14 000 published fMRI studies by searching for high-frequency key words (such as “pain” and “attention” terms) that are systematically mentioned in the papers alongside fMRI voxel co-ordinates (https://github.com/neurosynth/neurosynth), using the volumetric association test maps [62]. This measure of association strength is the tendency that a given term is reported in the functional neuroimaging study if there is activation observed at a given voxel. The probabilistic measure reported by NeuroSynth can be interpreted as a quantitative representation of how regional activity is related to psychological processes, as indicated by their co-occurrence in the published literature. Specifically, we focus on seven brain maps reflecting fundamental cognitive operations: attention, cognitive control, emotion, vision (‘fixation’), language, memory, and movement. For each simulated individual, their regional BOLD signals at each point in time were spatially correlated against each of the seven NeuroSynth-derived cognitive patterns, and the magnitudes of these spatial correlations were averaged over time to obtain a single value per cognitive pattern, per individual. For each meta-analytic pattern, the resulting value quantifies its prevalence in the spontaneous brain dynamics.

### Statistical analyses

The effect sizes were estimated using Hedge’s measure of the standardized mean difference, *g*, which is interpreted in the same way as Cohen’s *d*, but more appropriate for small sample sizes [255]. The Measures of Effect Size Toolbox for MATLAB https://github.com/ hhentschke/measures-of-effect-size-toolbox was used [256].

#### Spatial null models

The statistical significance of spatial correlation between brain maps was assessed non-parametrically via comparison against a null distribution of null maps with preserved spatial autocorrelation [114, 257]. For each map, parcel coordinates were projected onto the spherical surface and then randomly rotated and original parcels were reassigned the value of the closest rotated parcel (10 000 repetitions)

#### Meta-analysis with NeuroSynth

To contextualise the PC1 of local susceptibility, we performed a meta-analysis using 123 term-based meta-analytic brain maps from the NeuroSynth database (see above) [62]. Although more than a thousand terms are catalogued in the NeuroSynth engine, we refine our analysis by focusing on cognitive function and therefore we limit the terms of interest to cognitive and behavioural terms. To avoid introducing a selection bias, we opted for selecting terms in a data-driven fashion instead of selecting terms manually. Therefore, terms were selected from the Cognitive Atlas, a public ontology of cognitive science [258], which includes a comprehensive list of neurocognitive terms. This approach totaled to *t* = 123 terms, ranging from umbrella terms (“attention”, “emotion”) to specific cognitive processes (“visual attention”, “episodic memory”), behaviours (“eating”, “sleep”), and emotional states (“fear”, “anxiety”) (note that the 123 term-based meta-analytic maps from NeuroSynth do not explicitly exclude patient studies). The Cognitive Atlas subdivision has previously been used in conjunction with NeuroSynth [257, 259, 260], so we opted for the same approach to make our results comparable to previous reports. The full list of terms included in the present analysis is shown in Table **??**.

To perform this analysis, the PC1 map was spatially correlated against each NeuroSynth map, and statistical significance was assessed against our autocorrelation-preserving null model. The false positive rate against multiple comparisons was further controlled using the false discovery rate (FDR) correction [261].

#### Connectomic predictors of local susceptibility PC1

We characterised the PC1 of local susceptibility in terms of its relationship with several well-known graph-theoretic properties of the structural connectome. From the consensus connectome we computed the weighted degree (sum of a node’s connection weights); local efficiency (a measure of how well interconnected a node’s neighbours are); participation coefficient (diversity of modules that a node connects to, here defined based on the modular assignment of each region to the well-known intrinsic connectivity networks [262]); and average and modal controllability, which account for connectivity across paths of varying length to estimate a node’s ability to steer network dynamics towards easy-to-reach (average controllability) or hard-to-reach (modal controllability) [120, 263]. Average controllability of a network equals the average input energy needed at a set of control nodes, averaged over all possible target states. Average input energy is proportional to

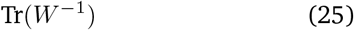

where *W* is the controllability Gramian. However, since the trace of the inverse Gramian is often uncomputable due to ill-conditioning, we follow previous work [120] in using

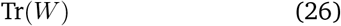

instead, which encodes the energy of the network impulse response, since the traces of the Gramian and its inverse are inversely proportional.

Modal controllability describes how easily the system can be induced to transition to a state that is distant on the energy landscape of its possible states. Technically, it corresponds to the ability of a node to control each of the dynamic modes of the network, and it can be computed from the matrix *V* of the eigenvectors of the adjacency matrix of the structural connectivity *A*. Following previous work, we define a scaled measure of the controllability from brain region *i* of all the *N* modes of the system as:

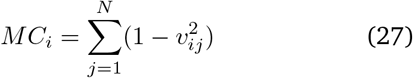

From this definition, a region will have high modal controllability if it is able to control all the dynamic modes of the system, which implies that it is well-suited to drive the system towards difficult-to-reach configurations in the energy landscape [120, 263].

#### Anatomical predictors of susceptibility PC1

We used *neuromaps* (https://netneurolab.github.io/neuromaps/) [264] to fetch the map of intracortical myelination obtained from T1w/T2w MRI ratio [10], the principal component of variation in gene expression from the Allen Human Brain Atlas transcriptomic database (‘AHBA PC1’) [10], the principal component of variation in task activation from the NeuroSynth database (‘NeuroSynth PC1’) [123], the principal component of variation in receptor density (‘receptor PC1’) [2], and the principal gradient of FC [55].

#### Dominance analysis

For both the anatomical and connectomic predictors, we used dominance analysis to consider all predictors together and evaluate their respective contributions. Dominance analysis seeks to determine the relative contribution (‘dominance’) of each independent variable to the overall fit of the multiple linear regression model when all predictors are considered (https://github.com/ dominance-analysis/dominance-analysis) [121]. This is done by fitting the same regression model on every combination of predictors (2^*p*^ ™1 submodels for a model with *p* predictors). Total dominance is defined as the average of the relative increase in explained variance (*R*^2^) when adding a single predictor of interest to a sub-model, across all 2^*p*^ ™ 1 submodels. The sum of the dominance of all input variables is equal to the total adjusted *R*^2^ of the complete model, making the percentage of relative importance an intuitive method that partitions the total effect size across predictors. Therefore, unlike other methods of assessing predictor importance, such as methods based on regression coefficients or univariate correlations, dominance analysis accounts for predictor–predictor interactions and is interpretable. We establish the statistical significance of the dominance analysis model using a non-parametric permutation test (one-sided), by comparing the empirical variance explained against a null distribution of *R*^2^ obtained from repeating the multiple regression with spatial autocorrelation-preserving null maps [114, 257].

## Data availability

We have made the results of all simulations openly available through an interactive website (https://systematic-causal-mapping.up.railway.app/). Human Connectome Project Young Adult resting-state, task-based, and diffusion MRI data are available from https://www.humanconnectome.org/study/hcp-young-adult. Diffusion MRI data for the Human Connectome Project in DSI Studio-compatible format are available at http://brain.labsolver.org/diffusion-mri-templates/hcp-842-hcp-1021. The ENIGMA cortical thickness data are provided as part of the ENIGMA Toolbox (v1.1.3), available at https://github.com/MICA-MNI/ENIGMA. PET receptor maps are available at https://github.com/netneurolab/hansen_receptors. NeuroSynth meta-analytic maps are freely available from the NeuroSynth database at https://github.com/neurosynth/neurosynth.

## Code Availability

Python and Octave/Matlab code for the dynamic mean-field model used in the present study is publicly available as FastDMF and maintained at http://www.gitlab.com/concog/fastdmf [64]. The Highly Comparative Time-Series Analysis (hctsa) toolbox is freely available at https://github.com/benfulcher/hctsa. The BrainSpace toolbox for gradient decomposition is available at https://brainspace.readthedocs.io/en/latest/. The Brain Connectivity Toolbox used for graph-theoretical properties and to generate rewired networks is freely available at https://sites.google.com/site/bctnet/. MATLAB code used to generate geometry-preserving null networks is freely available at https://www.brainnetworkslab.com/coderesources. The code for spin-based permutation testing of cortical correlations is freely available at https://github.com/frantisekvasa/rotate_parcellation. Third-party Python software (version 3.8 was used) for dominance analysis is freely available at https://github.com/dominance-analysis/dominance-analysis. The Enigma toolbox (v1.1.3) for fetching disorder-related maps is freely available at https://github.com/MICA-MNI/ ENIGMA. The Neuromaps toolbox for fetching brain maps (version 0.0.1) is freely available at https://netneurolab.github.io/neuromaps/.MATLAB/Octave and Python code to compute measures of Integrated Information Decomposition of timeseries with the Gaussian MMI solver is freely available at https://github.com/Imperial-MIND-lab/integrated-info-decomp.

## Author contributions

AIL, BM, MLK conceived the work. AIL performed the analysis. FM, LES, GS contributed to the analysis. JV, YSP, GD, MLK, FR, PAMM contributed to interpretation. HA developed the online website. AIL made figures. AIL wrote the first draft with feedback from all coauthors.

## Acknowledgments

A.I.L. acknowledges support from St John’s College, Cambridge; and a Wellcome Early Career Award (grant number 226924/Z/23/Z). G.S. was supported by a post-doctoral fellowship from the Canadian Institutes of Health Research (CIHR). The work of F.R. has been supported by the ARIA Safeguarded AI program and by PIBBSS affiliateship program. J.V. is supported by EU H2020 FET Proactive project Neurotwin (101017716). Y.S.P. is supported by European Union’s Horizon 2020 research and innovation programme under the Marie Sklodowska-Curie grant 896354, and ‘ERDF A way of making Europe’, ERDF, EU, Project NEurological MEchanismS of Injury, and Sleep-like cellular dynamics (NEMESIS; ref. 101071900) funded by the EU ERC Synergy Horizon Europe. F.M. was funded by a UNIQUE Neuro-AI Excellence Scholarship. M.L.K. is supported by the Center for Music in the Brain, funded by the Danish National Research Foundation (DNRF117), and Centre for Eudaimonia and Human Flourishing at Linacre College funded by the Pettit and Carlsberg Foundations. G.D. is supported by grant no. PID2022-136216NB-I00 funded by MICIU/AEI/10.13039/501100011033 and by ‘ERDF A way of making Europe’, ERDF, EU, Project NEurological MEchanismS of Injury, and Sleep-like cellular dynamics (NEMESIS; ref. 101071900) funded by the EU ERC Synergy Horizon Europe, and AGAUR research support grant (ref. 2021 SGR 00917) funded by the Department of Research and Universities of the Generalitat of Catalunya. B.M. acknowledges support from the Natural Sciences and Engineering Research Council of Canada (NSERC), Canadian Institutes of Health Research (CIHR), Brain Canada Foundation Future Leaders Fund, the Canada Research Chairs Program, the Michael J. Fox Foundation, and the Healthy Brains for Healthy Lives initiative. For the purpose of open access, the authors have applied a Creative Commons Attribution (CC BY) licence to any Author Accepted Manuscript version arising from this submission. Any opinions, findings, and conclusions or recommendations expressed in this material are those of the authors and do not reflect the views of the funders.

---

**TABLE 1.**
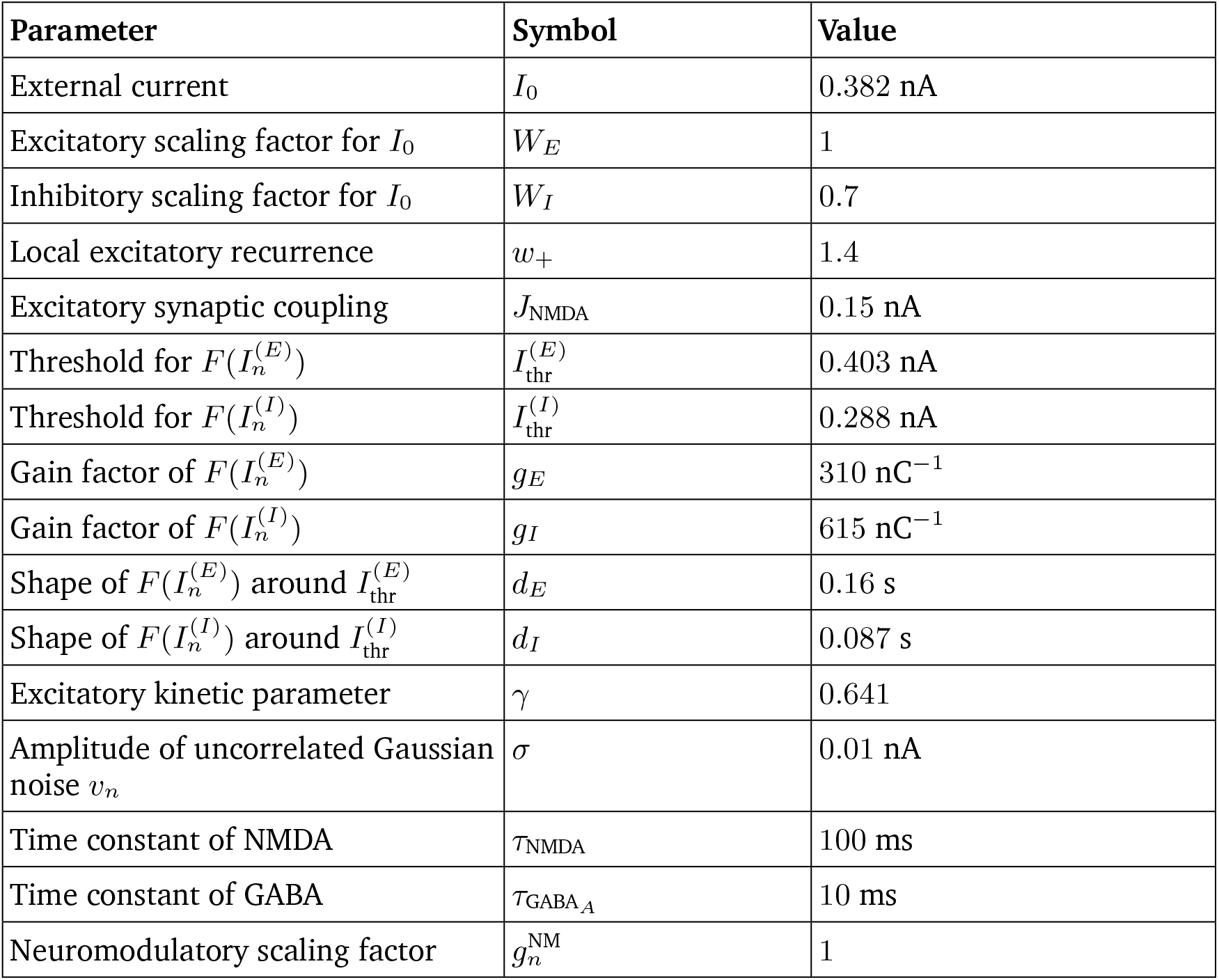
Fixed parameters of the dynamic mean field model and their values, as per [18].

**TABLE 2.** Representative subset of dynamical features from *catch22*, along with broad category assignment. Note that *ami2* (full feature ID: *CO*_*HistogramAMI*_*even*_2_5) was not included in the present work, as it failed to pass the pre-filtering stage.

| Feature name | Category | Description |
| --- | --- | --- |
| mode_5 | Distribution | 5-bin histogram mode |
| mode_10 | Distribution | 10-bin histogram mode |
| outlier_timing_pos | Outlier | Timing of positive extreme event |
| outlier_timing_neg | Outlier | Timing of negative extreme event |
| acf_timescale | Autocorrelation | First $1/e$ crossing of the ACF |
| acf_first_min | Autocorrelation | First minimum of the ACF |
| low_freq_power | Spectrum | Power in the lowest 20% frequencies |
| centroid_freq | Spectrum | Centroid frequency |
| forecast_error | Forecasting | Error of 3-point rolling mean forecast |
| whiten_timescale | Forecasting | Change in autocorrelation timescale after incremental differencing |
| High fluctuation | Other | Proportion of high incremental changes in the series |
| stretch_high | Symbolic | Longest stretch of above-mean values |
| stretch_decreasing | Symbolic | Longest stretch of decreasing values |
| entropy_pairs | Symbolic | Entropy of successive two-symbol motifs in the symbolized series |
| ami2 | Autocorrelation | Histogram-based automutual information (lag 2, 5 bins) |
| time_revers | Correlation | Time reversibility |
| ami_timescale | Autocorrelation | First minimum of the AMI function |
| transition_variance | Symbolic | Transition matrix column variance |
| periodicity | Other (periodicity) | Wang's periodicity metric |
| embedding_dist | Correlation | Goodness of exponential fit to embedding distance distribution |
| rs_range | Self-affine scaling | Rescaled range fluctuation analysis (low-scale scaling) |
| dfa | Self-affine scaling | Detrended fluctuation analysis (low-scale scaling) |

## References

[1] Alexandros Goulas, Jean-Pierre Changeux, Konrad Wagstyl, Katrin Amunts, Nicola Palomero-Gallagher, and Claus C Hilgetag. The natural axis of transmitter receptor distribution in the human cerebral cortex. Proceedings of the National Academy of Sciences, 118(3): e2020574118, 2021.

[2] Justine Y Hansen, Golia Shafiei, Ross D Markello, Sylvia Cox, Kelly Smart, Etienne Aumont, Stijn Servaes, Stephanie Scala, Gabriel Wainstein, Gleb Bezgin, Thomas Funck, W Schmitz, Marc-andre Bedard, R Nathan Spreng, Jean-paul Soucy, and Synthia Guimond. Mapping neurotransmitter systems to the structural and functional organization of the human neocortex. Nature Neuroscience, (accepted):1–26, 2022.

[3] Karl Zilles and Nicola Palomero-Gallagher. Multiple transmitter receptors in regions and layers of the human cerebral cortex. Frontiers in neuroanatomy, 11:78, 2017.

[4] Michael J Hawrylycz, Ed S Lein, Angela L Guillozet-Bongaarts, Elaine H Shen, Lydia Ng, Jeremy A Miller, Louie N Van De Lagemaat, Kimberly A Smith, Amanda Ebbert, Zackery L Riley, et al. An anatomically comprehensive atlas of the adult human brain transcriptome. Nature, 489(7416):391, 2012.

[5] Sean Froudist-Walsh, Ting Xu, Meiqi Niu, Lucija Rapan, Ling Zhao, Daniel S Margulies, Karl Zilles, Xiao-Jing Wang, and Nicola Palomero-Gallagher. Gradients of neurotransmitter receptor expression in the macaque cortex. Nature neuroscience, 26(7):1281–1294, 2023.

[6] Ed S Lein, Michael J Hawrylycz, Nancy Ao, Mikael Ayres, Amy Bensinger, Amy Bernard, Andrew F Boe, Mark S Boguski, Kevin S Brockway, Emi J Byrnes, et al. Genome-wide atlas of gene expression in the adult mouse brain. Nature, 445(7124):168–176, 2007.

[7] Katrin Amunts, Claude Lepage, Louis Borgeat, Hartmut Mohlberg, Timo Dickscheid, Marc-Étienne Rousseau, Sebastian Bludau, Pierre-Louis Bazin, Lindsay B Lewis, Ana-Maria Oros-Peusquens, et al. Bigbrain: an ultrahigh-resolution 3d human brain model. science, 340 (6139):1472–1475, 2013.

[8] Jakob Seidlitz, Ajay Nadig, Siyuan Liu, Richard AI Bethlehem, Petra E Vértes, Sarah E Morgan, František Váša, Rafael Romero-Garcia, François M Lalonde, Liv S Clasen, et al. Transcriptomic and cellular decoding of regional brain vulnerability to neurodevelopmental disorders. Nature Communications,.(.):., 2020.

[9] Matthew F Glasser and David C Van Essen. Mapping human cortical areas in vivo based on myelin content as revealed by t1-and t2-weighted mri. Journal of neuroscience, 31(32):11597–11616, 2011.

[10] Joshua B. Burt, Murat Demirtąs, William J. Eckner, Natasha M. Navejar, Jie Lisa Ji, William J. Martin, Alberto Bernacchia, Alan Anticevic, and John D. Murray. Hierarchy of transcriptomic specialization across human cortex captured by structural neuroimaging topography. Nature Neuroscience, 21(9):1251–1259, 9 2018. ISSN 15461726. doi:10.1038/s41593-018-0195-0. PMID: 30082915 publisher: Nature Publishing Group.

[11] Julia M Huntenburg, Pierre-Louis Bazin, Alexandros Goulas, Christine L Tardif, Arno Villringer, and Daniel S Margulies. A systematic relationship between functional connectivity and intracortical myelin in the human cerebral cortex. Cerebral Cortex, 27(2):981–997, 2017.

[12] Ben D Fulcher, John D Murray, Valerio Zerbi, and Xiao-Jing Wang. Multimodal gradients across mouse cortex. Proc Natl Acad Sci USA, 116(10):4689–4695, 2019.

[13] Laura E. Suárez, Ross D. Markello, Richard F. Betzel, and Bratislav Misic. Linking structure and function in macroscale brain networks. Trends in Cognitive Sciences, 24(4):302–315, 4 2020. ISSN 1879307X. doi:10.1016/j.tics.2020.01.008. PMID: 32160567 publisher: Elsevier Ltd.

[14] Xiao-Jing Wang. Macroscopic gradients of synaptic excitation and inhibition in the neocortex. Nat Rev Neurosci, pages 1–10, 2020.

[15] Songting Li and Xiao-Jing Wang. Hierarchical timescales in the neocortex: Mathematical mechanism and biological insights. Proceedings of the National Academy of Sciences, 119(6):e2110274119, 2022.

[16] Vincent Bazinet, Justine Y Hansen, and Bratislav Misic. Towards a biologically annotated brain connectome. Nature reviews neuroscience, 24(12):747–760, 2023.

[17] Andrea I. Luppi, Pedro A. M. Mediano, Fernando E. Rosas, Negin Holland, Tim D. Fryer, John T. O’Brien, James B. Rowe, David K. Menon, Daniel Bor, and Emmanuel A. Stamatakis. A synergistic core for human brain evolution and cognition. Nature Neuroscience, 25(6):771–782, 6 2022. ISSN 1546-1726. doi: 10.1038/s41593-022-01070-0. number: 6 publisher: Nature Publishing Group.

[18] Gustavo Deco, Josephine Cruzat, Joana Cabral, Gitte M Knudsen, Robin L Carhart-Harris, Peter C Whybrow, Nikos K Logothetis, and Morten L Kringelbach. Wholebrain multimodal neuroimaging model using serotonin receptor maps explains non-linear functional effects of lsd. Current biology, 28(19):3065–3074, 2018.

[19] Joshua B Burt, Katrin H Preller, Murat Demirtas, Jie Lisa Ji, John H Krystal, Franz X Vollenweider, Alan Anticevic, and John D Murray. Transcriptomics-informed largescale cortical model captures topography of pharmacological neuroimaging effects of lsd. Elife, 10:e69320, 2021.

[20] Murat Demirtąs, Joshua B Burt, Markus Helmer, Jie Lisa Ji, Brendan D Adkinson, Matthew F Glasser, David C Van Essen, Stamatios N Sotiropoulos, Alan Anticevic, and John D Murray. Hierarchical heterogeneity across human cortex shapes large-scale neural dynamics. Neuron, 101(6):1181–1194, 2019.

[21] Richard Gao, Ruud L van den Brink, Thomas Pfeffer, and Bradley Voytek. Neuronal timescales are functionally dynamic and shaped by cortical microarchitecture. Elife, 9: e61277, 2020.

[22] Golia Shafiei, Ross D Markello, Reinder Vos de Wael, Boris C Bernhardt, Ben D Fulcher, and Bratislav Misic. Topographic gradients of intrinsic dynamics across neocortex. elife, 9:e62116, 2020.

[23] Gustavo Deco, Morten L Kringelbach, Aurina Arnatkeviciute, Stuart Oldham, Kristina Sabaroedin, Nigel C Rogasch, Kevin M Aquino, and Alex Fornito. Dynamical consequences of regional heterogeneity in the brain’s transcriptional landscape. Science Advances, 7 (29):eabf4752, 2021.

[24] Panagiotis Fotiadis, Linden Parkes, Kathryn A Davis, Theodore D Satterthwaite, Russell T Shinohara, and Dani S Bassett. Structure–function coupling in macroscale human brain networks. Nature Reviews Neuroscience, 25(10):688–704, 2024.

[25] Ed Bullmore and Olaf Sporns. Complex brain networks: graph theoretical analysis of structural and functional systems. Nature reviews neuroscience, 10(3):186–198, 2009.

[26] Caio Seguin, Olaf Sporns, and Andrew Zalesky. Brain network communication: concepts, models and applications. Nature Reviews Neuroscience, pages 1–18, jul 12 2023. ISSN 1471-0048. doi:10.1038/s41583-023-00718-5. URL https://www.nature.com/articles/s41583-023-00718-5. publisher: Nature Publishing Group.

[27] Andrea Avena-Koenigsberger, Bratislav Misic, and Olaf Sporns. Communication dynamics in complex brain networks. Nature Reviews Neuroscience, 19(1):17–33, 2017. ISSN 1471-003X. doi:10.1038/nrn.2017.149. PMID: 29238085 publisher: Nature Publishing Group Citation Key: Avena-Koenigsberger2017.

[28] Kieran CR Fox, Lin Shi, Sori Baek, Omri Raccah, Brett L Foster, Srijani Saha, Daniel S Margulies, Aaron Kucyi, and Josef Parvizi. Intrinsic network architecture predicts the effects elicited by intracranial electrical stimulation of the human brain. Nature human behaviour, 4(10): 1039–1052, 2020.

[29] James M Shine, Michael Breakspear, Peter T Bell, Kaylena A Ehgoetz Martens, Richard Shine, Oluwasanmi Koyejo, Olaf Sporns, and Russell A Poldrack. Human cognition involves the dynamic integration of neural activity and neuromodulatory systems. Nat Neurosci, 22 (2):289–296, 2019.

[30] James M Shine, Patrick G Bissett, Peter T Bell, Oluwasanmi Koyejo, Joshua H Balsters, Krzysztof J Gorgolewski, Craig A Moodie, and Russell A Poldrack. The dynamics of functional brain networks: integrated network states during cognitive task performance. Neuron, 92(2):544–554, 2016.

[31] Joana Cabral, Morten L. Kringelbach, and Gustavo Deco. Functional connectivity dynamically evolves on multiple time-scales over a static structural connectome: Models and mechanisms. NeuroImage, 160(March):84–96, 2017. ISSN 10959572. doi:10.1016/j.neuroimage.2017.03.045. PMID: 28343985 publisher: Elsevier Citation Key: Cabral2017a ISBN: 1095-9572 (Electronic) 1053-8119 (Linking).

[32] C. J. Honey, O. Sporns, L. Cammoun, X. Gigandet, J. P. Thiran, R. Meuli, and P. Hagmann. Predicting human resting-state functional connectivity from structural connectivity. Proceedings of the National Academy of Sciences, 106(6):2035–2040, 2 2009. doi:10.1073/pnas.0811168106. publisher: Proceedings of the National Academy of Sciences.

[33] Pablo Barttfeld, Lynn Uhrig, Jacobo D Sitt, Mariano Sigman, Béchir Jarraya, and Stanislas Dehaene. Signature of consciousness in the dynamics of resting-state brain activity. Proceedings of the National Academy of Sciences, 112(3):887–892, 2015.

[34] Athena Demertzi, Enzo Tagliazucchi, Stanislas Dehaene, Gustavo Deco, Pablo Barttfeld, Federico Raimondo, Charlotte Martial, Davinia Fernández-Espejo, Benjamin Rohaut, HU Voss, et al. Human consciousness is supported by dynamic complex patterns of brain signal coordination. Science advances, 5(2):eaat7603, 2019.

[35] Michael D Fox. Mapping symptoms to brain networks with the human connectome. New England Journal of Medicine, 379(23):2237–2245, 2018.

[36] Aaron D Boes, Sashank Prasad, Hesheng Liu, Qi Liu, Alvaro Pascual-Leone, Verne S Caviness Jr, and Michael D Fox. Network localization of neurological symptoms from focal brain lesions. Brain, 138(10):3061–3075, 2015.

[37] Michael D. Fox and et al. Resting-state networks link invasive and noninvasive brain stimulation across diverse psychiatric and neurological diseases. Proceedings of the National Academy of Sciences of the United States of America, 111:E4367–E4375, 2014. doi: 10.1073/pnas.1405003111.

[38] Juho Joutsa, Michael D Fox, et al. Lesion network mapping for symptom localization: recent developments and future directions. Current opinion in neurology, 35(4): 453–459, 2022.

[39] Emmanuel Carrera and Giulio Tononi. Diaschisis: past, present, future. Brain, 137(9):2408–2422, 2014.

[40] Aaron D Boes and Michel Thiebaut de Schotten. Brain disconnections refine the relationship between brain structure and function. Brain structure and function, 227 (9):2893–2895, 2022.

[41] Shan H. Siddiqi, Konrad P. Kording, Josef Parvizi, and Michael D. Fox. Causal mapping of human brain function. Nature Reviews Neuroscience, 23(6):361–375, 2022. doi:10.1038/s41583-022-00583-8.

[42] Gaute T Einevoll, Alain Destexhe, Markus Diesmann, Sonja Grün, Viktor Jirsa, Marc de Kamps, Michele Migliore, Torbjørn V Ness, Hans E Plesser, and Felix Schürmann. The scientific case for brain simulations. Neuron, 102(4):735–744, 2019.

[43] James M. Shine, Eli J. Müller, Brandon Munn, Joana Cabral, Rosalyn J. Moran, and Michael Breakspear. Computational models link cellular mechanisms of neuromodulation to large-scale neural dynamics. Nature Neuroscience, pages 1–12, 5 2021. ISSN 1097-6256. doi: 10.1038/s41593-021-00824-6. publisher: Nature Publishing Group.

[44] Morten L. Kringelbach and Gustavo Deco. Brain states and transitions: Insights from computational neuroscience. Cell Reports, 32(10):108128, 9 2020. ISSN 22111247. doi:10.1016/j.celrep.2020.108128. PMID: 32905760 publisher: Elsevier B.V.

[45] Michael Breakspear. Dynamic models of large-scale brain activity. Nature Neuroscience, 20(3):340–352, 2017. ISSN 1097-6256. doi:10.1038/nn.4497. URL http://www.nature.com/doifinder/10.1038/nn.4497. PMID: 28230845 Citation Key: Breakspear2017 ISBN: 1546-1726 (Electronic) 1097-6256 (Linking).

[46] Julia K. Brynildsen, Kanaka Rajan, Michael X. Henderson, and Dani S. Bassett. Network models to enhance the translational impact of cross-species studies. Nature Reviews. Neuroscience, jul 31 2023. ISSN 1471-0048. doi:10.1038/s41583-023-00720-x. PMID: 37524935.

[47] Egidio D’Angelo and Viktor Jirsa. The quest for multiscale brain modeling. Trends in Neurosciences, 45(10):777–790, oct 1 2022. ISSN 0166-2236, 1878-108X. doi:10.1016/j.tins.2022.06.007. URL https://www.cell.com/trends/neurosciences/abstract/S0166-2236(22)00125-4. publisher: Elsevier PMID: 35906100.

[48] Gustavo Deco and Morten L Kringelbach. Great expectations: Using whole-brain computational connectomics for understanding neuropsychiatric disorders. Neuron, 84(5):892–905, 2014. ISSN 10974199. doi:10.1016/j.neuron.2014.08.034. PMID: 25475184.

[49] Petra Ritter, Michael Schirner, Anthony R McIntosh, and Viktor K Jirsa. The virtual brain integrates computational modeling and multimodal neuroimaging. Brain connectivity, 3(2):121–145, 2013.

[50] Andrea I Luppi, Joana Cabral, Rodrigo Cofre, Alain Destexhe, Gustavo Deco, and Morten L Kringelbach. Dynamical models to evaluate structure–function relationships in network neuroscience. Nature Reviews Neuroscience, 23(12):767–768, 2022.

[51] Maria Pope, Makoto Fukushima, Richard F Betzel, and Olaf Sporns. Modular origins of high-amplitude cofluctuations in fine-scale functional connectivity dynamics. Proceedings of the National Academy of Sciences, 118(46): e2109380118, 2021.

[52] Makoto Fukushima and Olaf Sporns. Structural determinants of dynamic fluctuations between segregation and integration on the human connectome. Communications biology, 3(1):606, 2020.

[53] Leonardo L Gollo, James A Roberts, Vanessa L Cropley, Maria A Di Biase, Christos Pantelis, Andrew Zalesky, and Michael Breakspear. Fragility and volatility of structural hubs in the human connectome. Nature neuroscience, 21 (8):1107–1116, 2018.

[54] Andrea I. Luppi, Fernando E. Rosas, Pedro A. M. Mediano, David K. Menon, and Emmanuel A. Stamatakis. Information decomposition and the informational architecture of the brain. Trends in Cognitive Sciences, 28(4):352–368, jan 9 2024. ISSN 1364-6613, 1879-307X. doi:10.1016/j.tics.2023.11.005. URL https://www.cell.com/trends/cognitive-sciences/abstract/S1364-6613(23)00284-X. publisher: Elsevier PMID: 38199949.

[55] Daniel S Margulies, Satrajit S Ghosh, Alexandros Goulas, Marcel Falkiewicz, Julia M Huntenburg, Georg Langs, Gleb Bezgin, Simon B Eickhoff, F Xavier Castellanos, Michael Petrides, Elizabeth Jefferies, and Jonathan Smallwood. Situating the default-mode network along a principal gradient of macroscale cortical organization. Proceedings of the National Academy of Sciences of the United States of America, 113(44):12574–12579, 2016. ISSN 1091-6490. doi:10.1073/pnas.1608282113. PMID: 27791099 arxiv: 1408.1149 Citation Key: Margulies2016 ISBN: 0027-8424.

[56] Rong Wang, Mianxin Liu, Xinhong Cheng, Ying Wu, Andrea Hildebrandt, and Changsong Zhou. Segregation, integration, and balance of large-scale resting brain networks configure different cognitive abilities. Proceedings of the National Academy of Sciences, 118(23): e2022288118, 2021.

[57] Gustavo Deco, Yonatan Sanz Perl, Hernan Bocaccio, Enzo Tagliazucchi, and Morten L. Kringelbach. The INSIDEOUT framework provides precise signatures of the balance of intrinsic and extrinsic dynamics in brain states. Communications Biology, 5(1):1–13, jun 10 2022. ISSN 2399-3642. doi:10.1038/s42003-022-03505-7. URL https://www.nature.com/articles/s42003-022-03505-7. number: 1 publisher: Nature Publishing Group.

[58] Fran Hancock, Fernando E Rosas, Andrea I Luppi, Mengsen Zhang, Pedro AM Mediano, Joana Cabral, Gustavo Deco, Morten L Kringelbach, Michael Breakspear, JA Kelso, et al. Metastability demystified—the foundational past, the pragmatic present and the promising future. Nature Reviews Neuroscience, pages 1–19, 2024.

[59] Murray Shanahan. Metastable chimera states in community-structured oscillator networks. Chaos (Woodbury, N.Y.), 20(1):013108, 3 2010. ISSN 1089-7682. doi:10.1063/1.3305451. PMID: 20370263.

[60] Ben D Fulcher, Max A Little, and Nick S Jones. Highly comparative time-series analysis: the empirical structure of time series and their methods. Journal of the Royal Society Interface, 10(83):20130048, 2013.

[61] Ben D Fulcher and Nick S Jones. hctsa: A computational framework for automated time-series phenotyping using massive feature extraction. Cell systems, 5(5):527–531, 2017.

[62] Tal Yarkoni, Russell A Poldrack, Thomas E Nichols, David C Van Essen, and Tor D Wager. Large-scale automated synthesis of human functional neuroimaging data. Nature methods, 8(8):665–670, 2011.

[63] Gustavo Deco, Adrián Ponce-Alvarez, Patric Hagmann, Gian Luca Romani, Dante Mantini, and Maurizio Corbetta. How local excitation–inhibition ratio impacts the whole brain dynamics. Journal of Neuroscience, 34(23): 7886–7898, 2014.

[64] Rubén Herzog, Pedro AM Mediano, Fernando E Rosas, Andrea I Luppi, Yonatan Sanz-Perl, Enzo Tagliazucchi, Morten L Kringelbach, Rodrigo Cofré, and Gustavo Deco. Neural mass modeling for the masses: Democratizing access to whole-brain biophysical modeling with fastdmf. Network Neuroscience, 8(4):1590–1612, 2024.

[65] Sara Larivière, Casey Paquola, Bo-yong Park, Jessica Royer, Yezhou Wang, Oualid Benkarim, de Reinder Vos Wael, Sofie L. Valk, Sophia I. Thomopoulos, Matthias Kirschner, Lindsay B. Lewis, Alan C. Evans, Sanjay M. Sisodiya, Carrie R. McDonald, Paul M. Thompson, and Boris C. Bernhardt. The enigma toolbox: multiscale neural contextualization of multisite neuroimaging datasets. Nature Methods 2021 18:7, 18(7):698–700, 6 2021. ISSN 1548-7105. doi:10.1038/s41592-021-01186-4. publisher: Nature Publishing Group.

[66] Paul M Thompson, Jason L Stein, Sarah E Medland, Derrek P Hibar, Alejandro Arias Vasquez, Miguel E Renteria, Roberto Toro, Neda Jahanshad, Gunter Schumann, Barbara Franke, et al. The enigma consortium: largescale collaborative analyses of neuroimaging and genetic data. Brain imaging and behavior, 8:153–182, 2014.

[67] Paul M Thompson, Neda Jahanshad, Christopher RK Ching, Lauren E Salminen, Sophia I Thomopoulos, Joanna Bright, Bernhard T Baune, Sara Bertolín, Janita Bralten, Willem B Bruin, et al. Enigma and global neuroscience: A decade of large-scale studies of the brain in health and disease across more than 40 countries. Translational psychiatry, 10(1):100, 2020.

[68] Pedro A M Mediano, Fernando E Rosas, Andrea I Luppi, Robin L Carhart-Harris, Daniel Bor, Anil K Seth, and Adam B Barrett. Towards an extended taxonomy of information dynamics via Integrated Information Decomposition. arXiv, 2021. doi: 10.48550/arXiv.2109.13186. URL http://arxiv.org/abs/2109.13186. arxiv: 2109.13186 ISBN: 2109.13186v1.

[69] Emmanuelle Tognoli and JA Scott Kelso. The metastable brain. Neuron, 81(1):35–48, 2014.

[70] Takuya Ito, Luke J Hearne, and Michael W Cole. A cortical hierarchy of localized and distributed processes revealed via dissociation of task activations, connectivity changes, and intrinsic timescales. NeuroImage, 221: 117141, 2020.

[71] Annemarie Wolff, Nareg Berberian, Mehrshad Golesorkhi, Javier Gomez-Pilar, Federico Zilio, and Georg Northoff. Intrinsic neural timescales: temporal integration and segregation. Trends in cognitive sciences, 26(2): 159–173, 2022.

[72] Sarah Feldt Muldoon, Eric W Bridgeford, and Danielle S Bassett. Small-world propensity and weighted brain networks. Scientific reports, 6(1):22057, 2016.

[73] Olaf Sporns and Richard F Betzel. Modular brain networks. Annual review of psychology, 67(1):613–640, 2016.

[74] Athena Demertzi, Aaron Kucyi, Adrián Ponce-Alvarez, Georgios A. Keliris, Susan Whitfield-Gabrieli, and Gustavo Deco. Functional network antagonism and consciousness. Network Neuroscience, 6(4):998–1009, oct 1 2022. ISSN 2472-1751. doi:10.1162/netn_a_00244. URL https://doi.org/10.1162/netn_a_00244. [Online; accessed 2023-03-23].

[75] Gustavo Deco and Morten L Kringelbach. Hierarchy of information processing in the brain: A novel ‘intrinsic ignition’ framework. Neuron, 94:961–968, 2017. doi:10.1016/j.neuron.2017.03.028. Citation Key: Deco2017.

[76] Andrea I. Luppi, Daniel Golkowski, Andreas Ranft, Rudiger Ilg, Denis Jordan, Danilo Bzdok, Adrian M. I. Owen, Lorina Naci, Emmanuel A. Stamatakis, Enrico A. Amico, and Bratislav Misic. General anaesthesia reduces the uniqueness of brain connectivity across individuals and across species. bioRxiv, nov 13 2023. doi:10.1101/2023.11.08.566332. URL https://www.biorxiv.org/content/10.1101/2023.11.08.566332v1. page: 2023.11.08.566332 section: New Results.

[77] Rahul S Desikan, Florent Ségonne, Bruce Fischl, Brian T Quinn, Bradford C Dickerson, Deborah Blacker, Randy L Buckner, Anders M Dale, R Paul Maguire, Bradley T Hyman, et al. An automated labeling system for subdividing the human cerebral cortex on mri scans into gyral based regions of interest. NeuroImage, 31(3):968–980, 2006.

[78] David C Van Essen, Stephen M Smith, Deanna M Barch, Timothy EJ Behrens, Essa Yacoub, Kamil Ugurbil, WuMinn HCP Consortium, et al. The wu-minn human connectome project: an overview. Neuroimage, 80:62–79, 2013.

[79] Marilyn Gatica, Fernando E. Rosas, Pedro AM Mediano, Ibai Diez, Stephan P. Swinnen, Patricio Orio, Rodrigo Cofré, and Jesus M. Cortes. High-order functional redundancy in ageing explained via alterations in the connectome in a whole-brain model. PLoS computational biology, 18(9):e1010431, 2022.

[80] Carlos Coronel-Oliveros, Carsten Gießing, Vicente Medel, Rodrigo Cofré, and Patricio Orio. Whole-brain modeling explains the context-dependent effects of cholinergic neuromodulation. Neuroimage, 265:119782, 2023.

[81] I Mindlin, R Herzog, L Belloli, D Manasova, M Monge-Asensio, J Vohryzek, A Escrichs, Naji Alnagger, P Núñez, Olivia Gosseries, et al. Whole brain modelling for simulating pharmacological interventions on patients with disorders of consciousness. Communications Biology, 7 (1):1176, 2024.

[82] Jakub Vohryzek, Joana Cabral, Yonatan Sanz Perl, Murat Demirtas, Carles Falcon, Juan Domingo Gispert, Beatriz Bosch, Mircea Balasa, Morten Kringelbach, Raquel Sanchez-Valle, et al. Design of effective personalised perturbation strategies for enhancing cognitive intervention in alzheimer’s disease. bioRxiv, pages 2023–04, 2023.

[83] Victor M Saenger, Joshua Kahan, Tom Foltynie, Karl Friston, Tipu Z Aziz, Alexander L Green, Tim J van Hartevelt, Joana Cabral, Angus BA Stevner, Henrique M Fernandes, et al. Uncovering the underlying mechanisms and whole-brain dynamics of deep brain stimulation for parkinson’s disease. Scientific reports, 7(1):9882, 2017.

[84] Enrique CA Hansen, Demian Battaglia, Andreas Spiegler, Gustavo Deco, and Viktor K Jirsa. Functional connectivity dynamics: modeling the switching behavior of the resting state. Neuroimage, 105:525–535, 2015.

[85] Gustavo Deco, Morten L Kringelbach, Viktor K Jirsa, and Petra Ritter. The dynamics of resting fluctuations in the brain: metastability and its dynamical cortical core. Scientific reports, 7(1):3095, 2017.

[86] Ane López-González, Rajanikant Panda, Adrián Ponce-Alvarez, Gorka Zamora-López, Anira Escrichs, Charlotte Martial, Aurore Thibaut, Olivia Gosseries, Morten L Kringelbach, Jitka Annen, et al. Loss of consciousness reduces the stability of brain hubs and the heterogeneity of brain dynamics. Communications biology, 4(1):1037, 2021.

[87] Andrea I Luppi, Justine Y Hansen, Ram Adapa, Robin L Carhart-Harris, Leor Roseman, Christopher Timmermann, Daniel Golkowski, Andreas Ranft, Rüdiger Ilg, Denis Jordan, et al. In vivo mapping of pharmacologically induced functional reorganization onto the human brain’s neurotransmitter landscape. Science advances, 9 (24):eadf8332, 2023.

[88] Susan C McKarns. A review of neuroreceptors for clinical and experimental neuropharmacology in central nervous system disorders. Current Reviews in Clinical and Experimental Pharmacology Formerly Current Clinical Pharmacology, 18(3):192–241, 2023.

[89] Solomon H Snyder. Drug and neurotransmitter receptors: new perspectives with clinical relevance. JAMA, 261(21):3126–3129, 1989.

[90] Morten L Kringelbach, Josephine Cruzat, Joana Cabral, Gitte Moos Knudsen, Robin Carhart-Harris, Peter C Whybrow, Nikos K Logothetis, and Gustavo Deco. Dynamic coupling of whole-brain neuronal and neurotransmitter systems. Proceedings of the National Academy of Sciences, 117(17):9566–9576, 2020.

[91] Andrea I Luppi, Pedro AM Mediano, Fernando E Rosas, Judith Allanson, John D Pickard, Guy B Williams, Michael M Craig, Paola Finoia, Alexander RD Peattie, Peter Coppola, et al. Whole-brain modelling identifies distinct but convergent paths to unconsciousness in anaesthesia and disorders of consciousness. Communications biology, 5(1):384, 2022.

[92] Shreyas Harita, Davide Momi, Zheng Wang, Sorenza P. Bastiaens, and John D. Griffiths. The role of inhibition in resting-state fMRI negative correlations. bioRxiv, apr 2 2024. doi:10.1101/2024.03.01.583030. URL https://www.biorxiv.org/content/10.1101/2024.03.01.583030v2. page: 2024.03.01.583030 section: New Results.

[93] Bolaji P Eniwaye, Victoria Booth, Anthony G Hudetz, and Michal Zochowski. Modeling cortical synaptic effects of anesthesia and their cholinergic reversal. PLOS Computational Biology, 18(6):e1009743, 2022.

[94] Camilo Miguel Signorelli, Lynn Uhrig, Morten Kringelbach, Bechir Jarraya, and Gustavo Deco. Hierarchical disruption in the cortex of anesthetized monkeys as a new signature of consciousness loss. NeuroImage, 227: 117618, 2021.

[95] Andrea I. Luppi, Lynn Uhrig, Jordy Tasserie, Camilo M. Signorelli, Emmanuel A. Stamatakis, Alain Destexhe, Bechir Jarraya, and Rodrigo Cofre. Local orchestration of distributed functional patterns supporting loss and restoration of consciousness in the primate brain. Nature Communications, 15(1):2171, mar 11 2024. ISSN 2041-1723. doi:10.1038/s41467-024-46382-w. URL https://www.nature.com/articles/s41467-024-46382-w. publisher: Nature Publishing Group.

[96] Zirui Huang, George A Mashour, and Anthony G Hudetz. Functional geometry of the cortex encodes dimensions of consciousness. Nature communications, 14(1):72, 2023.

[97] Joshua S Siegel, Subha Subramanian, Demetrius Perry, Benjamin P Kay, Evan M Gordon, Timothy O Laumann, T Rick Reneau, Nicholas V Metcalf, Ravi V Chacko, Caterina Gratton, et al. Psilocybin desynchronizes the human brain. Nature, 632(8023):131–138, 2024.

[98] Manesh Girn, Leor Roseman, Boris Bernhardt, Jonathan Smallwood, Robin Carhart-Harris, and R Nathan Spreng. Serotonergic psychedelic drugs lsd and psilocybin reduce the hierarchical differentiation of unimodal and transmodal cortex. NeuroImage, 256:119220, 2022.

[99] Christopher Timmermann, Leor Roseman, Sharad Haridas, Fernando E Rosas, Lisa Luan, Hannes Kettner, Jonny Martell, David Erritzoe, Enzo Tagliazucchi, Carla Pallavicini, et al. Human brain effects of dmt assessed via eeg-fmri. Proceedings of the National Academy of Sciences, 120(13):e2218949120, 2023.

[100] Justine Y Hansen, Golia Shafiei, Jacob W Vogel, Kelly Smart, Carrie E Bearden, Martine Hoogman, Barbara Franke, Daan Van Rooij, Jan Buitelaar, Carrie R McDonald, et al. Local molecular and global connectomic contributions to cross-disorder cortical abnormalities. Nature communications, 13(1):4682, 2022.

[101] Yonatan Sanz Perl, Sol Fittipaldi, Cecilia Gonzalez Campo, Sebastián Moguilner, Josephine Cruzat, Matias E Fraile-Vazquez, Rubén Herzog, Morten L Kringelbach, Gustavo Deco, Pavel Prado, Agustin Ibanez, and Enzo Tagliazucchi. Model-based whole-brain perturbational landscape of neurodegenerative diseases. eLife, 12:e83970, mar 30 2023. ISSN 2050-084X. doi: 10.7554/eLife.83970. URL https://doi.org/10.7554/eLife.83970. publisher: eLife Sciences Publications, Ltd.

[102] Andrea I. Luppi, S. Parker Singleton, Justine Y. Hansen, Danilo Bzdok, Amy Kuceyeski, Richard F. Betzel, and Bratislav Misic. Contributions of network structure, chemoarchitecture and diagnostic categories to transitions between cognitive topographies. Nature Biomedical Engineering, 8:1142–1161, 2024. doi: 10.1101/2023.03.16.532981. PMID: 36993597 PMCID: PMC10055141.

[103] Tahereh S Zarghami, Gholam-Ali Hossein-Zadeh, and Fariba Bahrami. Deep temporal organization of fmri phase synchrony modes promotes large-scale disconnection in schizophrenia. Frontiers in Neuroscience, 14:214, 2020.

[104] Mary-Ellen Lynall, Danielle S Bassett, Robert Kerwin, Peter J McKenna, Manfred Kitzbichler, Ulrich Muller, and Ed Bullmore. Functional connectivity and brain networks in schizophrenia. Journal of Neuroscience, 30(28): 9477–9487, 2010.

[105] Timothy J Spellman and Joshua A Gordon. Synchrony in schizophrenia: a window into circuit-level pathophysiology. Current opinion in neurobiology, 30:17–23, 2015.

[106] Danielle S Bassett, Brent G Nelson, Bryon A Mueller, Jazmin Camchong, and Kelvin O Lim. Altered resting state complexity in schizophrenia. Neuroimage, 59(3): 2196–2207, 2012.

[107] Miklos Argyelan, Toshikazu Ikuta, Pamela DeRosse, Raphael J Braga, Katherine E Burdick, Majnu John, Peter B Kingsley, Anil K Malhotra, and Philip R Szeszko. Resting-state fmri connectivity impairment in schizophrenia and bipolar disorder. Schizophrenia bulletin, 40(1):100–110, 2014.

[108] Maria Centeno and David W Carmichael. Network connectivity in epilepsy: resting state fmri and eeg–fmri contributions. Frontiers in neurology, 5:93, 2014.

[109] MJ Vaessen, HMH Braakman, JS Heerink, JFA Jansen, MHJA Debeij-van Hall, PAM Hofman, AP Aldenkamp, and WH Backes. Abnormal modular organization of functional networks in cognitively impaired children with frontal lobe epilepsy. Cerebral cortex, 23(8):1997– 2006, 2013.

[110] Michael B First. Mutually exclusive versus co-occurring diagnostic categories: the challenge of diagnostic comorbidity. Psychopathology, 38(4):206–210, 2005.

[111] Eleni Bonti, Irini K Zerva, Christiana Koundourou, and Maria Sofologi. The high rates of comorbidity among neurodevelopmental disorders: reconsidering the clinical utility of distinct diagnostic categories. Journal of personalized medicine, 14(3):300, 2024.

[112] Chao Xie, Shitong Xiang, Chun Shen, Xuerui Peng, Jujiao Kang, Yuzhu Li, Wei Cheng, Shiqi He, Marina Bobou, M John Broulidakis, et al. A shared neural basis underlying psychiatric comorbidity. Nature medicine, 29(5): 1232–1242, 2023.

[113] Pedro AM Mediano, Juan Carlos Farah, and Murray Shanahan. Integrated information and metastability in systems of coupled oscillators. arXiv preprint arxiv:1606.08313, 2016.

[114] František Váša and Bratislav Mišić. Null models in network neuroscience. Nature Reviews Neuroscience, pages 1–12, 2022.

[115] P Erdős and A Rényi. On the evolution of random graphs. ii. Bull. Inst. Int. Stat, 38(4):343–347, 1961.

[116] Sergei Maslov and Kim Sneppen. Specificity and stability in topology of protein networks. Science, 296(5569): 910–913, 5 2002. doi:10.1126/science.1065103. publisher: American Association for the Advancement of Science.

[117] Mikail Rubinov and Olaf Sporns. Weight-conserving characterization of complex functional brain networks. NeuroImage, 56(4):2068–2079, 2011. ISSN 10538119. doi:10.1016/j.neuroimage.2011.03.069. Citation Key: Rubinov2011.

[118] Richard F Betzel and Danielle S Bassett. Specificity and robustness of long-distance connections in weighted, interareal connectomes. Proc. Nati. Acad. Sci. USA, 2018. doi:10.1073/pnas.1720186115. [Online; accessed 2019-07-08].

[119] Ross D Markello and Bratislav Misic. Comparing spatial null models for brain maps. NeuroImage, page 118052, 2021.

[120] Shi Gu, Fabio Pasqualetti, Matthew Cieslak, Qawi K Telesford, Alfred B Yu, Ari E Kahn, John D Medaglia, Jean M Vettel, Michael B Miller, Scott T Grafton, and Danielle S Bassett. Controllability of structural brain networks. Nature Communications, 6:8414, 2015. ISSN 20411723. doi:10.1038/ncomms9414. PMID: 26423222 Citation Key: Gu2015.

[121] Razia Azen and David V Budescu. The dominance analysis approach for comparing predictors in multiple regression. Psychological methods, 8(2):129, 2003.

[122] Andrea I. Luppi, Pedro A. M. Mediano, Fernando E. Rosas, Judith Allanson, John D. Pickard, Guy B. Williams, Michael M. Craig, Paola Finoia, Alexander R. D. Peattie, Peter Coppola, David K. Menon, Daniel Bor, and Emmanuel A. Stamatakis. Reduced emergent character of neural dynamics in patients with a disrupted connectome. NeuroImage, 269:119926, apr 1 2023. ISSN 1095-9572. doi: 10.1016/j.neuroimage.2023.119926. PMID: 36740030 PMCID: PMC9989666.

[123] Valerie J Sydnor, Bart Larsen, Danielle S Bassett, Aaron Alexander-Bloch, Damien A Fair, Conor Liston, Allyson P Mackey, Michael P Milham, Adam Pines, David R Roalf, Jakob Seidlitz, Ting Xu, Armin Raznahan, and Theodore D Satterthwaite. Neurodevelopment of the association cortices: Patterns, mechanisms, and implications for psychopathology. Neuron, 2021. ISSN 08966273. doi:10.1016/j.neuron.2021.06.016. URL https://doi.org/10.1016/j.neuron.2021.06.016. [Online; accessed 2021-07-26].

[124] Carl H Lubba, Sarab S Sethi, Philip Knaute, Simon R Schultz, Ben D Fulcher, and Nick S Jones. catch22: Canonical time-series characteristics: Selected through highly comparative time-series analysis. Data Mining and Knowledge Discovery, 33(6):1821–1852, 2019.

[125] Stephen M Smith, Peter T Fox, Karla L Miller, David C Glahn, Peter Mickle Fox, Clare E Mackay, Nicola Filippini, Kate E Watkins, Roberto Toro, Angela R Laird, and Christian F Beckmann. Correspondence of the brain’s functional architecture during activation and rest. Proceedings of the National Academy of Sciences, 106(31):13040–13045, 2009. ISSN 0027-8424. doi: 10.1073/pnas.0905267106. PMID: 19620724 arxiv: 0905267106 Citation Key: Smith2009b ISBN: 1091-6490 (Electronic)\r1091-6490 (Linking).

[126] Michael W Cole, Danielle S Bassett, Jonathan D Power, Todd S Braver, and Steven E Petersen. Intrinsic and task-evoked network architectures of the human brain. Neuron, 83(1):238–251, 2014. ISSN 10974199. doi: 10.1016/j.neuron.2014.05.014. PMID: 24991964 arxiv: NIHMS150003 Citation Key: Cole2014 ISBN: 1097-4199 (Electronic)\r0896-6273 (Linking).

[127] BT Yeo, Fenna M Krienen, Jorge Sepulcre, Mert R Sabuncu, Danial Lashkari, Marisa Hollinshead, Joshua L Roffman, Jordan W Smoller, Lilla Zöllei, Jonathan R Polimeni, et al. The organization of the human cerebral cortex estimated by intrinsic functional connectivity. J Neurophysiol, 106(3):1125–1165, 2011.

[128] Nicolas A. Crossley, Andrea Mechelli, Petra E. Vértes, Toby T. Winton-Brown, Ameera X. Patel, Cedric E. Ginestet, Philip McGuire, and Edward T. Bullmore. Cognitive relevance of the community structure of the human brain functional coactivation network. Proceedings of the National Academy of Sciences, 110(28):11583– 11588, 2013. doi:10.1073/pnas.1220826110.

[129] Alex Fornito, Ben J. Harrison, Andrew Zalesky, and Jon S. Simons. Competitive and cooperative dynamics of large-scale brain functional networks supporting recollection. Proceedings of the National Academy of Sciences, 109(31):12788–12793, jul 31 2012. doi: 10.1073/pnas.1204185109. URL https://www.pnas.org/doi/abs/10.1073/pnas.1204185109. publisher: Proceedings of the National Academy of Sciences.

[130] Luca Cocchi, Andrew Zalesky, Alex Fornito, and Jason B. Mattingley. Dynamic cooperation and competition between brain systems during cognitive control. Trends in Cognitive Sciences, 17(10):493–501, oct 1 2013. ISSN 1364-6613. doi:10.1016/j.tics.2013.08.006. URL https://www.sciencedirect.com/science/article/pii/S136466131300171X. [Online; accessed 2024-09-25].

[131] František Váša, Murray Shanahan, Peter J Hellyer, Gregory Scott, Joana Cabral, and Robert Leech. Effects of lesions on synchrony and metastability in cortical networks. Neuroimage, 118:456–467, 2015.

[132] Joana Cabral, Etienne Hugues, Morten L Kringelbach, and Gustavo Deco. Modeling the outcome of structural disconnection on resting-state functional connectivity. Neuroimage, 62(3):1342–1353, 2012.

[133] Leonardo L. Gollo, Andrew Zalesky, R. Matthew Hutchison, Martijn Van Den Heuvel, and Michael Breakspear. Dwelling quietly in the rich club: brain network determinants of slow cortical fluctuations. Philosophical Transactions of the Royal Society B: Biological Sciences, 370(1668), may 19 2015. ISSN 14712970. doi:10.1098/RSTB.2014.0165. URL https://royalsocietypublishing.org/doi/abs/10.1098/rstb.2014.0165. PMID: 25823864 publisher: The Royal Society.

[134] Gustavo Deco, Josephine Cruzat, Joana Cabral, Enzo Tagliazucchi, Helmut Laufs, Nikos K. Logothetis, and Morten L. Kringelbach. Awakening: Predicting external stimulation to force transitions between different brain states. Proceedings of the National Academy of Sciences, page 201905534, 2019. ISSN 0027-8424. doi:10.1073/pnas.1905534116. URL http://www.pnas.org/lookup/doi/10.1073/pnas.1905534116. ISBN: 1905534116.

[135] Jing Wei, Bin Wang, Yanli Yang, Yan Niu, Lan Yang, Yuxiang Guo, and Jie Xiang. Effects of virtual lesions on temporal dynamics in cortical networks based on personalized dynamic models. NeuroImage, 254:119087, 2022. doi:10.1016/j.neuroimage.2022.119087.

[136] Christopher J. Honey and Olaf Sporns. Dynamical consequences of lesions in cortical networks. Human Brain Mapping, 29(7):802–809, 2008. doi:10.1002/hbm.20579.

[137] Jeffrey Alstott, Michael Breakspear, Patric Hagmann, Leila Cammoun, and Olaf Sporns. Modeling the impact of lesions in the human brain. PLoS Computational Biology, 5(6):e1000408, 2009. doi:10.1371/journal.pcbi.1000408.

[138] Leonardo L. Gollo, James A. Roberts, and Luca Cocchi. Mapping how local perturbations influence systems-level brain dynamics. NeuroImage, 160:97–112, 2017. doi:10.1016/j.neuroimage.2017.01.057.

[139] Kanika Bansal, Javier O. Garcia, Steven H. Tompson, Timothy Verstynen, Jean M. Vettel, and Sarah F. Muldoon. Cognitive chimera states in human brain networks. Science Advances, 5(4):eaau8535, 2019. doi:10.1126/sciadv.aau8535.

[140] Nicolas A. Crossley, Andrea Mechelli, Jessica Scott, Francesco Carletti, Peter T. Fox, Philip McGuire, and Edward T. Bullmore. The hubs of the human connectome are generally implicated in the anatomy of brain disorders. Brain, 137(8):2382–2395, 2014. doi: 10.1093/brain/awu132.

[141] Randy L. Buckner, Jorge Sepulcre, Tanveer Talukdar, Fenna M. Krienen, Hesheng Liu, Trey Hedden, Jessica R. Andrews-Hanna, Reisa A. Sperling, and Keith A. Johnson. Cortical hubs revealed by intrinsic functional connectivity: mapping, assessment of stability, and relation to alzheimer’s disease. The Journal of Neuroscience, 29 (6):1860–1873, 2009. doi:10.1523/JNEUROSCI.506208.2009.

[142] Olaf Sporns. Towards network substrates of brain disorders. Brain, 137(8):2117–2118, 2014. doi: 10.1093/brain/awu148.

[143] Guusje Collin, Olaf Sporns, René C. W. Mandl, and Martijn P. van den Heuvel. Structural and functional aspects relating to cost and benefit of rich club organization in the human cerebral cortex. Cerebral Cortex, 24(9):2258– 2267, 2014. doi:10.1093/cercor/bht064.

[144] Danielle S Bassett and Edward T Bullmore. Small-world brain networks revisited. The Neuroscientist, 23(5):499– 516, 2017.

[145] Claus C Hilgetag and Alexandros Goulas. ‘hierarchy’ in the organization of brain networks. Philosophical Transactions of the Royal Society B, 375(1796), 2020. doi: 10.1098/rstb.2019.0319. [Online; accessed 2020-11-10].

[146] Gustavo Deco, Giulio Tononi, Melanie Boly, et al. Rethinking segregation and integration: contributions of whole-brain modelling. Nature Reviews Neuroscience, 16 (7):430–439, 2015. doi:10.1038/nrn3963.

[147] Tim Shallice. From Neuropsychology to Mental Structure. Cambridge University Press, Cambridge, UK, 1988.

[148] Alexander R. Luria. The Working Brain: An Introduction to Neuropsychology. Penguin Books, Harmondsworth, UK, 1973.

[149] Avinash R. Vaidya, Maia S. Pujara, Michael Petrides, Elisabeth A. Murray, and Lesley K. Fellows. Lesion studies in contemporary neuroscience. Trends in Cognitive Sciences, 23(8):653–671, 2019. doi:10.1016/j.tics.2019.05.009.

[150] Ralph Adolphs. Human lesion studies in the 21st century. Neuron, 90(6):1151–1153, 2016. doi: 10.1016/j.neuron.2016.05.014.

[151] Juho Joutsa, Nir Lipsman, Andreas Horn, G Rees Cosgrove, and Michael D Fox. The return of the lesion for localization and therapy. Brain, 146(8):3146–3155, 2023. doi:10.1093/brain/awad123.

[152] Paul Broca. Remarques sur le siège de la faculté du langage articulé, suivies d’une observation d’aphémie (perte de la parole). Bulletin et Memoires de la Societe anatomique de Paris, 6:330–357, 1861.

[153] Carl Wernicke. Der aphasische Symptomencomplex: eine psychologische Studie auf anatomischer Basis. M. Crohn und Weigert, 1874.

[154] Brenda Milner. Les troubles de la memoire accompagnant des lesions hippocampiques bilaterales. In Physiologie de l’hippocampe, pages 257–272. Centre National de la Recherche Scientifique, 1962.

[155] Margaret J Moore, Nele Demeyere, Chris Rorden, and Jason B Mattingley. Lesion mapping in neuropsychological research: A practical and conceptual guide. Cortex, 170:38–52, 2024.

[156] Bertha K. Madras, Zhihua Xie, Zhicheng Lin, Amy Jassen, Helen Panas, Laurie Lynch, Ryan Johnson, Eli Livni, Thomas J. Spencer, Ali A. Bonab, Gregory M. Miller, and Alan J. Fischman. Modafinil occupies dopamine and norepinephrine transporters in vivo and modulates the transporters and trace amine activity in vitro. Journal of Pharmacology and Experimental Therapeutics, 319(2):561–569, 11 2006. ISSN 0022-3565, 1521-0103. doi:10.1124/jpet.106.106583. publisher: American Society for Pharmacology and Experimental Therapeutics section: NEUROPHARMACOLOGY PMID: 16885432.

[157] B. K. Madras. History of the discovery of the antipsychotic dopamine d2 receptor: a basis for the dopamine hypothesis of schizophrenia. Journal of the History of the Neurosciences, 22(1):62–78, 2013. doi:10.1080/0964704X.2012.678199.

[158] S. Kapur and G. Remington. Dopamine d(2) receptors and their role in atypical antipsychotic action: still necessary and may even be sufficient. Biological Psychiatry, 50(11):873–883, 2001. doi:10.1016/s0006-3223(01)01251-3.

[159] W. M. Greenberg and L. Citrome. Pharmacokinetics and pharmacodynamics of lurasidone hydrochloride, a second-generation antipsychotic: A systematic review of the published literature. Clinical Pharmacokinetics, 56, 2017. doi:10.1007/s40262-016-0424-0.

[160] J. B. Cohn and K. Rickels. A pooled, double-blind comparison of the effects of buspirone, diazepam and placebo in women with chronic anxiety. Current Medical Research and Opinion, 11(5):304–320, 1989. doi:10.1185/03057348909106107.

[161] J. F. Cryan, A. M. Redmond, J. P. Kelly, and B. E. Leonard. The effects of the 5-ht1a agonist flesinoxan, in three paradigms for assessing antidepressant potential in the rat. European Neuropsychopharmacology, 7, 1997. doi:10.1016/S0924-977X(97)80013-3.

[162] G. A. Kennett, C. T. Dourish, and G. Curzon. Antidepressant-like action of 5-ht1a agonists and conventional antidepressants in an animal model of depression. European Journal of Pharmacology, 134(2):183– 188, 1987. doi:10.1016/0014-2999(87)90363-2.

[163] A. D. Stark, S. Jordan, K. A. Allers, R. L. Bertekap, R. Chen, Kannan T. Mistry, et al. Interaction of the novel antipsychotic aripiprazole with 5-ht1a and 5-ht 2a receptors: functional receptor-binding and in vivo electrophysiological studies. Psychopharmacology, 190(3):373– 378, 2007. doi:10.1007/s00213-006-0631-3.

[164] Kayson Fakhar, Fatemeh Hadaeghi, Caio Seguin, Shrey Dixit, Arnaud Messé, Gorka Zamora-López, Bratislav Misic, and Claus C Hilgetag. A general framework for characterizing optimal communication in brain networks. eLife, 13:RP101780, 2024. doi: 10.7554/eLife.101780.1.

[165] Kayson Fakhar, Shrey Dixit, Farid Sadeghi, Konrad P Körding, and Claus C Hilgetag. Downstream network transformations dissociate neural activity from causal functional contributions. Scientific Reports, 14:2103, 2024. doi:10.1038/s41598-024-52423-7.

[166] Caroline Malherbe, Bastian Cheng, Alina Königsberg, Tae-Hee Cho, Martin Ebinger, Matthias Endres, Jochen B Fiebach, Jens Fiehler, Ivana Galinovic, Josep Puig, Vincent Thijs, Robin Lemmens, Keith W Muir, Norbert Nighoghossian, Salvador Pedraza, Claus Z Simonsen, Anke Wouters, Christian Gerloff, Claus C Hilgetag, and Götz Thomalla. Game-theoretical mapping of fundamental brain functions based on lesion deficits in acute stroke. Brain Communications, 3(3):fcab204, 2021. doi: 10.1093/braincomms/fcab204.

[167] Melissa Zavaglia, Caroline Malherbe, Sebastian Schlaadt, Parashkev Nachev, and Claus C. Hilgetag. Ground-truth validation of uni-and multivariate lesion inference approaches. Brain Communications, 6: fcae251, 2024. doi:10.1093/braincomms/fcae251.

[168] James C Pang, James K Rilling, James A Roberts, Martijn P van den Heuvel, and Luca Cocchi. Evolutionary shaping of human brain dynamics. eLife, 11:e80627, oct 26 2022. ISSN 2050-084X. doi:10.7554/eLife.80627. URL https://doi.org/10.7554/eLife.80627. publisher: eLife Sciences Publications, Ltd.

[169] Jacob Tanner, Joshua Faskowitz, Andreia Sofia Teixeira, Caio Seguin, Ludovico Coletta, Alessandro Gozzi, Bratislav Mišić, and Richard F. Betzel. A multimodal, asymmetric, weighted, and signed description of anatomical connectivity. Nature Communications, 15(1):5865, jul 12 2024. ISSN 2041-1723. doi:10.1038/s41467-024-50248-6. URL https://www.nature.com/articles/s41467-024-50248-6. publisher: Nature Publishing Group.

[170] Andrea I. Luppi, Yonatan Sanz Perl, Jakub Vohryzek, Pedro A. M. Mediano, Fernando E. Rosas, Filip Milisav, Laura E. Suarez, Silvia Gini, Daniel Gutierrez-Barragan, Alessandro Gozzi, Bratislav Misic, Gustavo Deco, and Morten L. Kringelbach. Competitive interactions shape brain dynamics and computation across species. bioRxiv, 2024. doi:10.1101/2024.10.19.619194.

[171] Matthew D. Greaves, Leonardo Novelli, Sina L. Mansour, Andrew Zalesky, and Adeel Razi. Structurally informed models of directed brain connectivity. Nature Reviews Neuroscience, 26(1):23–41, 2025. doi:10.1038/s41583-024-00881-3.

[172] Michael Schirner, Gustavo Deco, and Petra Ritter. Learning how network structure shapes decision-making for bio-inspired computing. Nature Communications, 14(1):2963, may 23 2023. ISSN 2041-1723. doi:10.1038/s41467-023-38626-y. URL https://www.nature.com/articles/s41467-023-38626-y. number: 1 publisher: Nature Publishing Group.

[173] Alexandros Goulas, Fabrizio Damicelli, and Claus C. Hilgetag. Bio-instantiated recurrent neural networks: Integrating neurobiology-based network topology in artificial networks. Neural Networks, jul 24 2021. ISSN 0893-6080. doi:10.1016/J.NEUNET.2021.07.011. URL https://linkinghub.elsevier.com/retrieve/pii/S0893608021002744. publisher: Pergamon.

[174] Laura E. Suárez, Blake A. Richards, Guillaume Lajoie, and Bratislav Misic. Learning function from structure in neuromorphic networks. Nature Machine Intelligence 2021, pages 1–16, aug 9 2021. ISSN 2522-5839. doi:10.1038/s42256-021-00376-1. URL https://www.nature.com/articles/s42256-021-00376-1. publisher: Nature Publishing Group.

[175] Laura E. Suárez, Agoston Mihalik, Filip Milisav, Kenji Marshall, Mingze Li, Petra E. Vértes, Guillaume Lajoie, and Bratislav Misic. Connectome-based reservoir computing with the conn2res toolbox. Nature Communications, 15(1):656, jan 22 2024. ISSN 2041-1723. doi:10.1038/s41467-024-44900-4. URL https://www.nature.com/articles/s41467-024-44900-4. publisher: Nature Publishing Group.

[176] F. Rocchi, C. Canella, S. Noei, et al. Increased fmri connectivity upon chemogenetic inhibition of the mouse prefrontal cortex. Nature Communications, 13:1056, 2022. doi:10.1038/s41467-022-28591-3.

[177] Marija Markicevic, Oliver Sturman, Johannes Bohacek, Markus Rudin, Valerio Zerbi, Ben D. Fulcher, and Nicole Wenderoth. Neuromodulation of striatal d1 cells shapes bold fluctuations in anatomically connected thalamic and cortical regions. eLife, 12:e78620, 2023. doi:10.7554/eLife.78620.

[178] Andreas Spiegler, Javad Karimi Abadchi, Majid Mohajerani, and Viktor K. Jirsa. In silico exploration of mouse brain dynamics by focal stimulation reflects the organization of functional networks and sensory processing. Network Neuroscience, 4(3):807–851, 2020. doi:10.1162/netn_a_00152.

[179] David C Van Essen, Stephen M Smith, Deanna M Barch, Timothy E.J. Behrens, Essa Yacoub, and Kamil Ugurbil. The wu-minn human connectome project: An overview. NeuroImage, 80:62–79, 2013. ISSN 10538119. doi:10.1016/j.neuroimage.2013.05.041. PMID: 23684880.

[180] Matthew F Glasser, Stamatios N Sotiropoulos, Anthony Wilson, Timothy S Coalson, Bruce Fischl, Jesper L Andersson, Junqian Xu, Saad Jbabdi, Matthew Webster, Jonathan R Polimeni, David C Van Essen, and Mark Jenkinson. The minimal preprocessing pipelines for the human connectome project. Neuroimage, 80:105–124, 2013. doi:10.1016/j.neuroimage.2013.04.127.

[181] Fang-Cheng Yeh, Sandip Panesar, David Fernandes, Antonio Meola, Masanori Yoshino, Juan C Fernandez-Miranda, Jean M Vettel, and Timothy Verstynen. Population-averaged atlas of the macroscale human structural connectome and its network topology. NeuroImage, 178:57–68, 2018. ISSN 10959572. doi:10.1016/j.neuroimage.2018.05.027. PMID: 29758339.

[182] Behzadi Y, Restom K, Liau J, and Liu TT. A component based noise correction method (CompCor) for BOLD and perfusion based fMRI. NeuroImage, 37:90– 101, 2007. [Online; accessed 2019-06-26].

[183] Susan Whitfield-Gabrieli and Alfonso Nieto-Castanon. Conn: A Functional Connectivity Toolbox for Correlated and Anticorrelated Brain Networks. Brain Connectivity, 2(3):125–141, 2012. ISSN 21580022. doi: 10.1089/brain.2012.0073. Citation Key: Whitfield-Gabrieli.

[184] Rahul S Desikan, Florent Ségonne, Bruce Fischl, Brian T Quinn, Bradford C Dickerson, Deborah Blacker, Randy L Buckner, Anders M Dale, R Paul Maguire, Bradley T Hyman, et al. An automated labeling system for subdividing the human cerebral cortex on mri scans into gyral based regions of interest. Neuroimage, 31(3):968–980, 2006.

[185] Fang-Cheng Yeh, Van Jay Wedeen, and Wen-Yih Isaac Tseng. Estimation of fiber orientation and spin density distribution by diffusion deconvolution. NeuroImage, 55(3):1054–1062, 4 2011. ISSN 1053-8119. doi: 10.1016/J.NEUROIMAGE.2010.11.087. publisher: Academic Press.

[186] Fang-Cheng Yeh, T D Verstynen, Y Wang, J C Fernández-Miranda, and W-Yi Tseng. Deterministic diffusion fiber tracking improved by quantitative anisotropy. PLoS ONE, 8(11):80713, 2013. doi: 10.1371/journal.pone.0080713.

[187] Andrea I Luppi and Emmanuel A Stamatakis. Combining network topology and information theory to construct representative brain networks. Network Neuroscience, 5(1):96–124, 2021. ISSN 2472-1751. doi: 10.1162/netn_a_00170.

[188] Richard F Betzel, Alessandra Griffa, Patric Hagmann, and Bratislav Mišić. Distance-dependent consensus thresholds for generating group-representative structural brain networks. Network neuroscience, 3(2):475– 496, 2019.

[189] Bratislav Mišić, Richard F Betzel, Azadeh Nematzadeh, Joaquin Goni, Alessandra Griffa, Patric Hagmann, Alessandro Flammini, Yong-Yeol Ahn, and Olaf Sporns. Cooperative and competitive spreading dynamics on the human connectome. Neuron, 86(6):1518–1529, 2015.

[190] K. F. Wong and X. J. Wang. A recurrent network mechanism of time integration in perceptual decisions. Journal of Neuroscience, 26:1314–1328, 2006. doi: 10.1523/JNEUROSCI.3733-05.2006.

[191] S. Kirkpatrick, C. D. Jr. Gelatt, and M. P. Vecchi. Optimization by simulated annealing. Science, 220(4598): 671–680, 1983. doi:10.1126/science.220.4598.671.

[192] Nicholas Metropolis, Arianna W. Rosenbluth, Marshall N. Rosenbluth, Augusta H. Teller, and Edward Teller. Equation of state calculations by fast computing machines. Journal of Chemical Physics, 21:1087, 1953. doi:10.1063/1.1699114.

[193] Daqiang Sun, Christopher R. K. Ching, Amy Lin, Jennifer K. Forsyth, Leila Kushan, Ariana Vajdi, Maria Jalbrzikowski, Laura Hansen, Julio E. Villalon-Reina, Xiaoping Qu, Rachel K. Jonas, Therese van Amelsvoort, Geor Bakker, Wendy R. Kates, Kevin M. Antshel, Wanda Fremont, Linda E. Campbell, Kathryn L. McCabe, Eileen Daly, Maria Gudbrandsen, Clodagh M. Murphy, Declan Murphy, Michael Craig, Jacob Vorstman, Ania Fiksinski, Sanne Koops, Kosha Ruparel, David R. Roalf, Raquel E. Gur, J. Eric Schmitt, Tony J. Simon, Naomi J. Goodrich-Hunsaker, Courtney A. Durdle, Anne S. Bassett, Eva W. C. Chow, Nancy J. Butcher, Fidel Vila-Rodriguez, Joanne Doherty, Adam Cunningham, Marianne B. M. van den Bree, David E. J. Linden, Hayley Moss, Michael J. Owen, Kieran C. Murphy, Donna M. McDonald-McGinn, Beverly Emanuel, Theo G. M. van Erp, Jessica A. Turner, Paul M. Thompson, and Carrie E. Bearden. Large-scale mapping of cortical alterations in 22q11.2 deletion syndrome: Convergence with idiopathic psychosis and effects of deletion size. Molecular Psychiatry, 25(8):1822–1834, 8 2020. ISSN 1476-5578. doi:10.1038/s41380-018-0078-5. PMID: 29895892 PMCID: PMC6292748.

[194] Martine Hoogman, Ryan Muetzel, Joao P. Guimaraes, Elena Shumskaya, Maarten Mennes, Marcel P. Zwiers, Neda Jahanshad, Gustavo Sudre, Thomas Wolfers, Eric A. Earl, Juan Carlos Soliva Vila, Yolanda Vives-Gilabert, Sabin Khadka, Stephanie E. Novotny, Catharina A. Hartman, Dirk J. Heslenfeld, Lizanne J. S. Schweren, Sara Ambrosino, Bob Oranje, Patrick de Zeeuw, Tiffany M. Chaim-Avancini, Pedro G. P. Rosa, Marcus V. Zanetti, Charles B. Malpas, Gregor Kohls, Georg G. von Polier, Jochen Seitz, Joseph Biederman, Alysa E. Doyle, Anders M. Dale, Theo G. M. van Erp, Jeffery N. Epstein, Terry L. Jernigan, Ramona Baur-Streubel, Georg C. Ziegler, Kathrin C. Zierhut, Anouk Schrantee, Marie F. Høvik, Astri J. Lundervold, Clare Kelly, Hazel McCarthy, Norbert Skokauskas, Ruth L. O’Gorman Tuura, Anna Calvo, Sara Lera-Miguel, Rosa Nicolau, Kaylita C. Chantiluke, Anastasia Christakou, Alasdair Vance, Mara Cercignani, Matt C. Gabel, Philip Asherson, Sarah Baumeister, Daniel Brandeis, Sarah Hohmann, Ivanei E. Bramati, Fernanda Tovar-Moll, Andreas J. Fallgatter, Bernd Kardatzki, Lena Schwarz, Anatoly Anikin, Alexandr Baranov, Tinatin Gogberashvili, Dmitry Kapilushniy, Anastasia Solovieva, Hanan El Marroun, Tonya White, Georgii Karkashadze, Leyla Namazova-Baranova, Thomas Ethofer, Paulo Mattos, Tobias Banaschewski, David Coghill, Kerstin J. Plessen, Jonna Kuntsi, Mitul A. Mehta, Yannis Paloyelis, Neil A. Harrison, Mark A. Bellgrove, Tim J. Silk, Ana I. Cubillo, Katya Rubia, Luisa Lazaro, Silvia Brem, Susanne Walitza, Thomas Frodl, Mariam Zentis, Francisco X. Castellanos, Yuliya N. Yoncheva, Jan Haavik, Liesbeth Reneman, Annette Conzelmann, Klaus-Peter Lesch, Paul Pauli, Andreas Reif, Leanne Tamm, Kerstin Konrad, Eileen Oberwelland Weiss, Geraldo F. Busatto, Mario R. Louza, Sarah Durston, Pieter J. Hoekstra, Jaap Oosterlaan, Michael C. Stevens, J. Antoni Ramos-Quiroga, Oscar Vilarroya, Damien A. Fair, Joel T. Nigg, Paul M. Thompson, Jan K. Buitelaar, Stephen V. Faraone, Philip Shaw, Henning Tiemeier, Janita Bralten, and Barbara Franke. Brain imaging of the cortex in adhd: A coordinated analysis of large-scale clinical and population-based samples. The American Journal of Psychiatry, 176(7):531–542, 7 2019. ISSN 1535-7228. doi:10.1176/appi.ajp.2019.18091033. PMID: 31014101 PMCID: PMC6879185.

[195] Daan van Rooij, Evdokia Anagnostou, Celso Arango, Guillaume Auzias, Marlene Behrmann, Geraldo F. Busatto, Sara Calderoni, Eileen Daly, Christine Deruelle, Adriana Di Martino, Ilan Dinstein, Fabio Luis Souza Duran, Sarah Durston, Christine Ecker, Damien Fair, Jennifer Fedor, Jackie Fitzgerald, Christine M. Freitag, Louise Gallagher, Ilaria Gori, Shlomi Haar, Liesbeth Hoekstra, Neda Jahanshad, Maria Jalbrzikowski, Joost Janssen, Jason Lerch, Beatriz Luna, Mauricio Moller Martinho, Jane McGrath, Filippo Muratori, Clodagh M. Murphy, Declan G. M. Murphy, Kirsten O’Hearn, Bob Oranje, Mara Parellada, Alessandra Retico, Pedro Rosa, Katya Rubia, Devon Shook, Margot Taylor, Paul M. Thompson, Michela Tosetti, Gregory L. Wallace, Fengfeng Zhou, and Jan K. Buitelaar. Cortical and subcortical brain morphometry differences between patients with autism spectrum disorder and healthy individuals across the lifespan: Results from the enigma asd working group. The American Journal of Psychiatry, 175(4):359–369, 4 2018. ISSN 1535-7228. doi: 10.1176/appi.ajp.2017.17010100. PMID: 29145754 PMCID: PMC6546164.

[196] Christopher D. Whelan, Andre Altmann, Juan A. Botía, Neda Jahanshad, Derrek P. Hibar, Julie Absil, Saud Alhusaini, Marina K. M. Alvim, Pia Auvinen, Emanuele Bartolini, Felipe P. G. Bergo, Tauana Bernardes, Karen Blackmon, Barbara Braga, Maria Eugenia Caligiuri, Anna Calvo, Sarah J. Carr, Jian Chen, Shuai Chen, Andrea Cherubini, Philippe David, Martin Domin, Sonya Foley, Wendy França, Gerrit Haaker, Dmitry Isaev, Simon S. Keller, Raviteja Kotikalapudi, Magdalena A. Kowalczyk, Ruben Kuzniecky, Soenke Langner, Matteo Lenge, Kelly M. Leyden, Min Liu, Richard Q. Loi, Pascal Martin, Mario Mascalchi, Marcia E. Morita, Jose C. Pariente, Raul Rodríguez-Cruces, Christian Rummel, Taavi Saavalainen, Mira K. Semmelroch, Mariasavina Severino, Rhys H. Thomas, Manuela Tondelli, Domenico Tortora, Anna Elisabetta Vaudano, Lucy Vivash, Felix von Podewils, Jan Wagner, Bernd Weber, Yi Yao, Clarissa L. Yasuda, Guohao Zhang, Nuria Bargalló, Benjamin Bender, Neda Bernasconi, Andrea Bernasconi, Boris C. Bernhardt, Ingmar Blümcke, Chad Carlson, Gianpiero L. Cavalleri, Fernando Cendes, Luis Concha, Norman Delanty, Chantal Depondt, Orrin Devinsky, Colin P. Doherty, Niels K. Focke, Antonio Gambardella, Renzo Guerrini, Khalid Hamandi, Graeme D. Jackson, Reetta Kälviäinen, Peter Kochunov, Patrick Kwan, Angelo Labate, Carrie R. McDonald, Stefano Meletti, Terence J. O’Brien, Sebastien Ourselin, Mark P. Richardson, Pasquale Striano, Thomas Thesen, Roland Wiest, Junsong Zhang, Annamaria Vezzani, Mina Ryten, Paul M. Thompson, and Sanjay M. Sisodiya. Structural brain abnormalities in the common epilepsies assessed in a worldwide enigma study. Brain: A Journal of Neurology, 141(2):391–408, 2 2018. ISSN 1460-2156. doi:10.1093/brain/awx341. PMID: 29365066 PMCID: PMC5837616.

[197] L. Schmaal, D. P. Hibar, P. G. Sämann, G. B. Hall, B. T. Baune, N. Jahanshad, J. W. Cheung, T. G. M. van Erp, D. Bos, M. A. Ikram, M. W. Vernooij, W. J. Niessen, H. Tiemeier, A. Hofman, K. Wittfeld, H. J. Grabe, D. Janowitz, R. Bülow, M. Selonke, H. Völzke, D. Grotegerd, U. Dannlowski, V. Arolt, N. Opel, W. Heindel, H. Kugel, D. Hoehn, M. Czisch, B. Couvy-Duchesne, M. E. Rentería, L. T. Strike, M. J. Wright, N. T. Mills, G. I. de Zubicaray, K. L. McMahon, S. E. Medland, N. G. Martin, N. A. Gillespie, R. Goya-Maldonado, O. Gruber, B. Krämer, S. N. Hatton, J. Lagopoulos, I. B. Hickie, T. Frodl, A. Carballedo, E. M. Frey, L. S. van Velzen, B. W. J. H. Penninx, M.-J. van Tol, N. J. van der Wee, C. G. Davey, B. J. Harrison, B. Mwangi, B. Cao, J. C. Soares, I. M. Veer, H. Walter, D. Schoepf, B. Zurowski, C. Konrad, E. Schramm, C. Normann, K. Schnell, M. D. Sacchet, I. H. Gotlib, G. M. MacQueen, B. R. Godlewska, T. Nickson, A. M. McIntosh, M. Papmeyer, H. C. Whalley, J. Hall, J. E. Sussmann, M. Li, M. Walter, L. Aftanas, I. Brack, N. A. Bokhan, P. M. Thompson, and D. J. Veltman. Cortical abnormalities in adults and adolescents with major depression based on brain scans from 20 cohorts worldwide in the enigma major depressive disorder working group. Molecular Psychiatry, 22(6):900–909, 6 2017. ISSN 1476-5578. doi:10.1038/mp.2016.60. PMID: 27137745 PMCID: PMC5444023.

[198] Premika S. W. Boedhoe, Lianne Schmaal, Yoshinari Abe, Pino Alonso, Stephanie H. Ameis, Alan Anticevic, Paul D. Arnold, Marcelo C. Batistuzzo, Francesco Benedetti, Jan C. Beucke, Irene Bollettini, Anushree Bose, Silvia Brem, Anna Calvo, Rosa Calvo, Yuqi Cheng, Kang Ik K. Cho, Valentina Ciullo, Sara Dallaspezia, Damiaan Denys, Jamie D. Feusner, Kate D. Fitzgerald, Jean-Paul Fouche, Egill A. Fridgeirsson, Patricia Gruner, Gregory L. Hanna, Derrek P. Hibar, Marcelo Q. Hoexter, Hao Hu, Chaim Huyser, Neda Jahanshad, Anthony James, Norbert Kathmann, Christian Kaufmann, Kathrin Koch, Jun Soo Kwon, Luisa Lazaro, Christine Lochner, Rachel Marsh, Ignacio Martínez-Zalacaín, David Mataix-Cols, José M. Menchón, Luciano Minuzzi, Astrid Morer, Takashi Nakamae, Tomohiro Nakao, Janardhanan C. Narayanaswamy, Seiji Nishida, Erika Nurmi, Joseph O’Neill, John Piacentini, Fabrizio Piras, Federica Piras, Y. C. Janardhan Reddy, Tim J. Reess, Yuki Sakai, Joao R. Sato, H. Blair Simpson, Noam Soreni, Carles Soriano-Mas, Gianfranco Spalletta, Michael C. Stevens, Philip R. Szeszko, David F. Tolin, Guido A. van Wingen, Ganesan Venkatasubramanian, Susanne Walitza, Zhen Wang, Je-Yeon Yun, ENIGMA-OCD Working Group, Paul M. Thompson, Dan J. Stein, Odile A. van den Heuvel, and ENIGMA OCD Working Group. Cortical abnormalities associated with pediatric and adult obsessive-compulsive disorder: Findings from the enigma obsessive-compulsive disorder working group. The American Journal of Psychiatry, 175(5):453–462, 5 2018. ISSN 1535-7228. doi:10.1176/appi.ajp.2017.17050485. PMID: 29377733 PMCID: PMC7106947.

[199] Theo G. M. van Erp, Esther Walton, Derrek P. Hibar, Lianne Schmaal, Wenhao Jiang, David C. Glahn, Godfrey D. Pearlson, Nailin Yao, Masaki Fukunaga, Ryota Hashimoto, Naohiro Okada, Hidenaga Yamamori, Juan R. Bustillo, Vincent P. Clark, Ingrid Agartz, Bryon A. Mueller, Wiepke Cahn, Sonja M. C. de Zwarte, Hilleke E. Hulshoff Pol, René S. Kahn, Roel A. Ophoff, Neeltje E. M. van Haren, Ole A. Andreassen, Anders M. Dale, Nhat Trung Doan, Tiril P. Gurholt, Cecilie B. Hartberg, Unn K. Haukvik, Kjetil N. Jørgensen, Trine V. Lagerberg, Ingrid Melle, Lars T. Westlye, Oliver Gruber, Bernd Kraemer, Anja Richter, David Zilles, Vince D. Calhoun, Benedicto Crespo-Facorro, Roberto Roiz-Santiañez, Diana Tordesillas-Gutiérrez, Carmel Loughland, Vaughan J. Carr, Stanley Catts, Vanessa L. Cropley, Janice M. Fullerton, Melissa J. Green, Frans A. Henskens, Assen Jablensky, Rhoshel K. Lenroot, Bryan J. Mowry, Patricia T. Michie, Christos Pantelis, Yann Quidé, Ulrich Schall, Rodney J. Scott, Murray J. Cairns, Marc Seal, Paul A. Tooney, Paul E. Rasser, Gavin Cooper, Cynthia Shannon Weickert, Thomas W. Weickert, Derek W. Morris, Elliot Hong, Peter Kochunov, Lauren M. Beard, Raquel E. Gur, Ruben C. Gur, Theodore D. Satterthwaite, Daniel H. Wolf, Aysenil Belger, Gregory G. Brown, Judith M. Ford, Fabio Macciardi, Daniel H. Mathalon, Daniel S. O’Leary, Steven G. Potkin, Adrian Preda, James Voyvodic, Kelvin O. Lim, Sarah McEwen, Fude Yang, Yunlong Tan, Shuping Tan, Zhiren Wang, Fengmei Fan, Jingxu Chen, Hong Xiang, Shiyou Tang, Hua Guo, Ping Wan, Dong Wei, Henry J. Bockholt, Stefan Ehrlich, Rick P. F. Wolthusen, Margaret D. King, Jody M. Shoemaker, Scott R. Sponheim, Lieuwe De Haan, Laura Koenders, Marise W. Machielsen, Therese van Amelsvoort, Dick J. Veltman, Francesca Assogna, Nerisa Banaj, Pietro de Rossi, Mariangela Iorio, Fabrizio Piras, Gianfranco Spalletta, Peter J. McKenna, Edith Pomarol-Clotet, Raymond Salvador, Aiden Corvin, Gary Donohoe, Sinead Kelly, Christopher D. Whelan, Erin W. Dickie, David Rotenberg, Aristotle N. Voineskos, Simone Ciufolini, Joaquim Radua, Paola Dazzan, Robin Murray, Tiago Reis Marques, Andrew Simmons, Stefan Borgwardt, Laura Egloff, Fabienne Harrisberger, Anita Riecher-Rössler, Renata Smieskova, Kathryn I. Alpert, Lei Wang, Erik G. Jönsson, Sanne Koops, Iris E. C. Sommer, Alessandro Bertolino, Aurora Bonvino, Annabella Di Giorgio, Emma Neilson, Andrew R. Mayer, Julia M. Stephen, Jun Soo Kwon, Je-Yeon Yun, Dara M. Cannon, Colm McDonald, Irina Lebedeva, Alexander S. Tomyshev, Tolibjohn Akhadov, Vasily Kaleda, Helena Fatouros-Bergman, Lena Flyckt, Karolinska Schizophrenia Project, Geraldo F. Busatto, Pedro G. P. Rosa, Mauricio H. Serpa, Marcus V. Zanetti, Cyril Hoschl, Antonin Skoch, Filip Spaniel, David Tomecek, Saskia P. Hagenaars, Andrew M. McIntosh, Heather C. Whalley, Stephen M. Lawrie, Christian Knöchel, Viola Oertel-Knöchel, Michael Stäblein, Fleur M. Howells, Dan J. Stein, Henk S. Temmingh, Anne Uhlmann, Carlos Lopez-Jaramillo, Danai Dima, Agnes McMahon, Joshua I. Faskowitz, Boris A. Gutman, Neda Jahanshad, Paul M. Thompson, and Jessica A. Turner. Cortical brain abnormalities in 4474 individuals with schizophrenia and 5098 control subjects via the enhancing neuro imaging genetics through meta analysis (enigma) consortium. Biological Psychiatry, 84(9):644–654, 11 2018. ISSN 1873-2402. doi:10.1016/j.biopsych.2018.04.023. PMID: 29960671 PMCID: PMC6177304.

[200] D. P. Hibar, L. T. Westlye, N. T. Doan, N. Jahanshad, J. W. Cheung, C. R. K. Ching, A. Versace, A. C. Bilderbeck, A. Uhlmann, B. Mwangi, B. Krämer, B. Overs, C. B. Hartberg, C. Abé, D. Dima, D. Grotegerd, E. Sprooten, E. Bøen, E. Jimenez, F. M. Howells, G. Delvecchio, H. Temmingh, J. Starke, J. R. C. Almeida, J. M. Goikolea, J. Houenou, L. M. Beard, L. Rauer, L. Abramovic, M. Bonnin, M. F. Ponteduro, M. Keil, M. M. Rive, N. Yao, N. Yalin, P. Najt, P. G. Rosa, R. Redlich, S. Trost, S. Hagenaars, S. C. Fears, S. Alonso-Lana, T. G. M. van Erp, T. Nickson, T. M. Chaim-Avancini, T. B. Meier, T. Elvsåshagen, U. K. Haukvik, W. H. Lee, A. H. Schene, A. J. Lloyd, A. H. Young, A. Nugent, A. M. Dale, A. Pfennig, A. M. McIntosh, B. Lafer, B. T. Baune, C. J. Ekman, C. A. Zarate, C. E. Bearden, C. Henry, Simhandl, C. McDonald, C. Bourne, D. J. Stein, H. Wolf, D. M. Cannon, D. C. Glahn, D. J. Veltman, Pomarol-Clotet, E. Vieta, E. J. Canales-Rodriguez, G. Nery, F. L. S. Duran, G. F. Busatto, G. Roberts, G. D. Pearlson, G. M. Goodwin, H. Kugel, H. C. Whalley, H. G. Ruhe, J. C. Soares, J. M. Fullerton, J. K. Rybakowski, J. Savitz, K. T. Chaim, M. Fatjó-Vilas, M. G. Soeiro-de Souza, M. P. Boks, M. V. Zanetti, M. C. G. Otaduy, M. S. Schaufelberger, M. Alda, M. Ingvar, M. L. Phillips, J. Kempton, M. Bauer, M. Landén, N. S. Lawrence, E. M. van Haren, N. R. Horn, N. B. Freimer, O. Gruber, P. R. Schofield, P. B. Mitchell, R. S. Kahn, R. Lenroot, R. Machado-Vieira, R. A. Ophoff, S. Sarró, S. Frangou, T. D. Satterthwaite, T. Hajek, U. Dannlowski, U. F. Malt, V. Arolt, W. F. Gattaz, W. C. Drevets, X. Caseras, I. Agartz, P. M. Thompson, and O. A. Andreassen. Cortical abnormalities in bipolar disorder: an mri analysis of 6503 individuals from the enigma bipolar disorder working group. Molecular Psychiatry, 23(4):932–942, 4 2018. ISSN 1476-5578. doi:10.1038/mp.2017.73. PMID: 28461699 PMCID: PMC5668195.

[201] Max A. Laansma, Joanna K. Bright, Sarah Al-Bachari, Tim J. Anderson, Tyler Ard, Francesca Assogna, Katherine A. Baquero, Henk W. Berendse, Jamie Blair, Fernando Cendes, John C. Dalrymple-Alford, Rob M.A. de Bie, Ines Debove, Michiel F. Dirkx, Jason Druzgal, Hedley C.A. Emsley, Gäetan Garraux, Rachel P. Guimarães, Boris A. Gutman, Rick C. Helmich, Johannes C. Klein, Clare E. Mackay, Corey T. McMillan, Tracy R. Melzer, Laura M. Parkes, Fabrizio Piras, Toni L. Pitcher, Kathleen L. Poston, Mario Rango, Letícia F. Ribeiro, Cristiane S. Rocha, Christian Rummel, Lucas S.R. Santos, Reinhold Schmidt, Petra Schwingenschuh, Gianfranco Spalletta, Letizia Squarcina, Odile A. van den Heuvel, Chris Vriend, Jiun-Jie Wang, Daniel Weintraub, Roland Wiest, Clarissa L. Yasuda, Neda Jahanshad, Paul M. Thompson, Ysbrand D. van der Werf, and The ENIGMA-Parkinson’s Study. International multicenter analysis of brain structure across clinical stages of parkinson’s disease. Movement Disorders, 36(11):2583–2594, 2021. ISSN 1531-8257. doi:10.1002/mds.28706. preprint: https://onlinelibrary.wiley.com/doi/pdf/10.1002/mds.28706.

[202] Simon Kaller, Michael Rullmann, Marianne Patt, Georg Becker, Julia Luthardt, Johanna Girbardt, Philipp Meyer, Peter Werner, Henryk Barthel, Anke McLeod, Thomas Fritz, Swen Hesse, and Osama Sabri. Test–retest measurements of dopamine d1-type receptors using simultaneous pet/mri imaging. European Journal of Nuclear Medicine and Molecular Imaging, 44, 6 2017. doi:10.1007/s00259-017-3645-0.

[203] Christine M. Sandiego, Jean-Dominique Gallezot, Keunpoong Lim, Jim Ropchan, Shu-fei Lin, Hong Gao, Evan D. Morris, and Kelly P. Cosgrove. Reference region modeling approaches for amphetamine challenge studies with [11c]flb 457 and pet. Journal of Cerebral Blood Flow and Metabolism: Official Journal of the International Society of Cerebral Blood Flow and Metabolism, 35(4):623–629, 3 2015. ISSN 1559-7016. doi:10.1038/jcbfm.2014.237. PMID: 25564239 PMCID: PMC4420880.

[204] Mark Slifstein, Elsmarieke van de Giessen, Jared Van Snellenberg, Judy L. Thompson, Rajesh Narendran, Roberto Gil, Elizabeth Hackett, Ragy Girgis, Najate Ojeil, Holly Moore, Deepak D’Souza, Robert T. Malison, Yiyun Huang, Keunpoong Lim, Nabeel Nabulsi, Richard E. Carson, Jeffrey A. Lieberman, and Anissa Abi-Dargham. Deficits in prefrontal cortical and extrastriatal dopamine release in schizophrenia: a positron emission tomographic functional magnetic resonance imaging study. JAMA psychiatry, 72(4):316–324, 4 2015. ISSN 2168-6238. doi:10.1001/jamapsychiatry.2014.2414. PMID: 25651194 PMCID: PMC4768742.

[205] Christopher T. Smith, Jennifer L. Crawford, Linh C. Dang, Kendra L. Seaman, M. Danica San Juan, Aishwarya Vijay, Daniel T. Katz, David Matuskey, Ronald L. Cowan, Evan D. Morris, David H. Zald, and Gregory R. Samanez-Larkin. Partial-volume correction increases estimated dopamine d2-like receptor binding potential and reduces adult age differences. Journal of Cerebral Blood Flow and Metabolism: Official Journal of the International Society of Cerebral Blood Flow and Metabolism, 39(5):822–833, 5 2019. ISSN 1559-7016. doi:10.1177/0271678X17737693. PMID: 29090626 PMCID: PMC6498753.

[206] Yasmin Zakiniaeiz, Ansel T. Hillmer, David Matuskey, Nabeel Nabulsi, Jim Ropchan, Carolyn M. Mazure, Marina R. Picciotto, Yiyun Huang, Sherry A. McKee, Evan D. Morris, and Kelly P. Cosgrove. Sex differences in amphetamine-induced dopamine release in the dorsolateral prefrontal cortex of tobacco smokers. Neuropsychopharmacology: Official Publication of the American College of Neuropsychopharmacology, 44(13):2205– 2211, 12 2019. ISSN 1740-634X. doi:10.1038/s41386-019-0456-y. PMID: 31269510 PMCID: PMC6897943.

[207] Markus Savli, Andreas Bauer, Markus Mitterhauser, Yu-Shin Ding, Andreas Hahn, Tina Kroll, Alexander Neumeister, Daniela Haeusler, Johanna Ungersboeck, Shannan Henry, Sanaz Attaripour Isfahani, Frank Rattay, Wolfgang Wadsak, Siegfried Kasper, and Rupert Lanzenberger. Normative database of the serotonergic system in healthy subjects using multi-tracer pet. NeuroImage, 63(1):447–459, 10 2012. ISSN 1053-8119. doi: 10.1016/j.neuroimage.2012.07.001.

[208] Jean-Dominique Gallezot, Nabeel Nabulsi, Alexander Neumeister, Beata Planeta-Wilson, Wendol A. Williams, Tarun Singhal, Sunhee Kim, R. Paul Maguire, Timothy McCarthy, J. James Frost, Yiyun Huang, Yu-Shin Ding, and Richard E. Carson. Kinetic modeling of the serotonin 5-ht(1b) receptor radioligand [(11)c]p943 in humans. Journal of Cerebral Blood Flow and Metabolism: Official Journal of the International Society of Cerebral Blood Flow and Metabolism, 30(1):196–210, 1 2010. ISSN 1559-7016. doi:10.1038/jcbfm.2009.195. PMID: 19773803 PMCID: PMC2949107.

[209] David Matuskey, Zubin Bhagwagar, Beata Planeta, Brian Pittman, Jean-Dominique Gallezot, Jason Chen, Jane Wanyiri, Soheila Najafzadeh, Jim Ropchan, Paul Geha, Yiyun Huang, Marc N. Potenza, Alexander Neumeister, Richard E. Carson, and Robert T. Malison. Reductions in brain 5-ht1b receptor availability in primarily cocaine-dependent humans. Biological Psychiatry, 76(10):816–822, 11 2014. ISSN 1873-2402. doi:10.1016/j.biopsych.2013.11.022. PMID: 24433854 PMCID: PMC4037398.

[210] James W. Murrough, Christoph Czermak, Shannan Henry, Nabeel Nabulsi, Jean-Dominique Gallezot, Ralitza Gueorguieva, Beata Planeta-Wilson, John H. Krystal, John F. Neumaier, Yiyun Huang, Yu-Shin Ding, Richard E. Carson, and Alexander Neumeister. The effect of early trauma exposure on serotonin type 1b receptor expression revealed by reduced selective radioligand binding. Archives of General Psychiatry, 68(9):892–900, 9 2011. ISSN 1538-3636. doi:10.1001/archgenpsychiatry.2011.91. PMID: 21893657 PMCID: PMC3244836.

[211] Christopher Pittenger, Thomas G. Adams, Jean-Dominique Gallezot, Michael J. Crowley, Nabeel Nabulsi, null James Ropchan, Hong Gao, Stephen A. Kichuk, Ryan Simpson, Eileen Billingslea, Jonas Hannestad, Michael Bloch, Linda Mayes, Zubin Bhagwagar, and Richard E. Carson. Ocd is associated with an altered association between sensorimotor gating and cortical and subcortical 5-ht1b receptor binding. Journal of Affective Disorders, 196:87–96, 5 2016. ISSN 1573-2517. doi:10.1016/j.jad.2016.02.021. PMID: 26919057 PMCID: PMC4808438.

[212] Aybala Saricicek, Jason Chen, Beata Planeta, Barbara Ruf, Kalyani Subramanyam, Kathleen Maloney, David Matuskey, David Labaree, Lorenz Deserno, Alexander Neumeister, John H. Krystal, Jean-Dominique Gallezot, Yiyun Huang, Richard E. Carson, and Zubin Bhagwagar. Test-retest reliability of the novel 5-ht1b receptor pet radioligand [11c]p943. European Journal of Nuclear Medicine and Molecular Imaging, 42(3):468–477, 3 2015. ISSN 1619-7089. doi:10.1007/s00259-014-2958-5. PMID: 25427881.

[213] Vincent Beliveau, Melanie Ganz, Ling Feng, Brice Ozenne, Liselotte Højgaard, Patrick M. Fisher, Claus Svarer, Douglas N. Greve, and Gitte M. Knudsen. A high-resolution in vivo atlas of the human brain’s serotonin system. The Journal of Neuroscience, 37(1):120, 1 2017. doi:10.1523/JNEUROSCI.2830-16.2016. PMID: 28053035 publisher: Society for Neuroscience.

[214] Rajiv Radhakrishnan, Nabeel Nabulsi, Edward Gaiser, Jean-Dominique Gallezot, Shannan Henry, Beata Planeta, Shu-fei Lin, Jim Ropchan, Wendol Williams, Evan Morris, Deepak Cyril D’Souza, Yiyun Huang, Richard E. Carson, and David Matuskey. Age-related change in 5-ht6 receptor availability in healthy male volunteers measured with 11c-gsk215083 pet. Journal of Nuclear Medicine, 59(9):1445–1450, 9 2018. ISSN 0161-5505. doi:10.2967/jnumed.117.206516. PMID: 29626125 PMCID: PMC6126437.

[215] Rajiv Radhakrishnan, David Matuskey, Nabeel Nabulsi, Edward Gaiser, Jean-Dominique Gallezot, Shannan Henry, Beata Planeta, Shu-Fei Lin, Jim Ropchan, Yiyun Huang, Richard E. Carson, and Deepak Cyril D’Souza. In vivo 5-ht6 and 5-ht2a receptor availability in antipsychotic treated schizophrenia patients vs. unmedicated healthy humans measured with [11c]gsk215083 pet. Psychiatry Research. Neuroimaging, 295:111007, 1 2020. ISSN 1872-7506. doi: 10.1016/j.pscychresns.2019.111007. PMID: 31760336.

[216] Stephen R. Baldassarri, Ansel T. Hillmer, Jon Mikael Anderson, Peter Jatlow, Nabeel Nabulsi, David Labaree, Kelly P. Cosgrove, Stephanie S. O’Malley, Thomas Eissenberg, Suchitra Krishnan-Sarin, and Irina Esterlis. Use of electronic cigarettes leads to significant beta2-nicotinic acetylcholine receptor occupancy: Evidence from a pet imaging study. Nicotine and Tobacco Research: Official Journal of the Society for Research on Nicotine and Tobacco, 20(4):425–433, 3 2018. ISSN 1469-994X. doi:10.1093/ntr/ntx091. PMID: 28460123 PMCID: PMC5896427.

[217] A. T. Hillmer, I. Esterlis, J. D. Gallezot, F. Bois, M. Q. Zheng, N. Nabulsi, S. F. Lin, R. L. Papke, Y. Huang, Sabri, R. E. Carson, and K. P. Cosgrove. Imaging of cerebral alpha4beta2* nicotinic acetylcholine receptors with (-)-[(18)f]flubatine pet: Implementation of bolus plus constant infusion and sensitivity to acetylcholine in human brain. NeuroImage, 141:71–80, 11 2016. ISSN 1095-9572. doi:10.1016/j.neuroimage.2016.07.026. PMID: 27426839 PMCID: PMC5026941.

[218] Mika Naganawa, Nabeel Nabulsi, Shannan Henry, David Matuskey, Shu-Fei Lin, Lawrence Slieker, Adam J. Schwarz, Nancy Kant, Cynthia Jesudason, Kevin Ruley, Antonio Navarro, Hong Gao, Jim Ropchan, David Labaree, Richard E. Carson, and Yiyun Huang. Firstin-human assessment of 11c-lsn3172176, an m1 muscarinic acetylcholine receptor pet radiotracer. Journal of Nuclear Medicine: Official Publication, Society of Nuclear Medicine, 62(4):553–560, 4 2021. ISSN 1535-5667. doi: 10.2967/jnumed.120.246967. PMID: 32859711 PMCID: PMC8049371.

[219] Jonathan M. DuBois, Olivier G. Rousset, Jared Rowley, Manuel Porras-Betancourt, Andrew J. Reader, Aurelie Labbe, Gassan Massarweh, Jean-Paul Soucy, Pedro Rosa-Neto, and Eliane Kobayashi. Characterization of age/sex and the regional distribution of mglur5 availability in the healthy human brain measured by high-resolution [(11)c]abp688 pet. European Journal of Nuclear Medicine and Molecular Imaging, 43(1):152–162, 1 2016. ISSN 1619-7089. doi:10.1007/s00259-015-3167-6. PMID: 26290423.

[220] Kelly Smart, Sylvia M. L. Cox, Stephanie G. Scala, Maria Tippler, Natalia Jaworska, Michel Boivin, Jean R. Séguin, Chawki Benkelfat, and Marco Leyton. Sex differences in [11c]abp688 binding: a positron emission tomography study of mglu5 receptors. European Journal of Nuclear Medicine and Molecular Imaging, 46(5):1179–1183, 2019. ISSN 1619-7070. doi:10.1007/s00259-018-4252-4. PMID: 30627817 PMCID: PMC6451701.

[221] Marian Galovic, Adam Al-Diwani, Umesh Vivekananda, Francisco Torrealdea, Kjell Erlandsson, Tim D. Fryer, Young T. Hong, Benjamin A. Thomas, Colm J. McGinnity, Evan Edmond, Kerstin Sander, Erik Årstad, Ilijas Jelcic, Franklin I. Aigbirhio, Ashley M. Groves, Kris Thielemans, Brian Hutton, Alexander Hammers, John S. Duncan, Jonathan P. Coles, Anna Barnes, Charlotte J. Stagg, Matthew C. Walker, Sarosh R. Irani, Matthias J. Koepp, and for the NEST Investigators. In vivo nmda receptor function in people with nmda receptor antibody encephalitis. medRxiv, 12 2021. doi:10.1101/2021.12.04.21267226. URL https://www.medrxiv.org/content/10.1101/2021.12.04.21267226v1. page: 2021.12.04.21267226.

[222] Marian Galovic, Kjell Erlandsson, Tim D. Fryer, Young T. Hong, Roido Manavaki, Hasan Sari, Sarah Chetcuti, Benjamin A. Thomas, Martin Fisher, Selena Sephton, Roberto Canales, Joseph J. Russell, Kerstin Sander, Erik Årstad, Franklin I. Aigbirhio, Ashley M. Groves, John S. Duncan, Kris Thielemans, Brian F. Hutton, Jonathan P. Coles, Matthias J. Koepp, and NEST investigators. Validation of a combined image derived input function and venous sampling approach for the quantification of [18f]ge-179 pet binding in the brain. NeuroImage, 237:118194, 8 2021. ISSN 1095-9572. doi:10.1016/j.neuroimage.2021.118194. PMID: 34023451.

[223] Martin Nørgaard, Vincent Beliveau, Melanie Ganz, Claus Svarer, Lars Pinborg, Sune Keller, Peter Jensen, Douglas Greve, and Gitte Knudsen. A high-resolution in vivo atlas of the human brain’s benzodiazepine binding site of gaba a receptors. NeuroImage, 232:117878, 2021. doi:10.1101/2020.04.10.035352. ISBN: 4535456720.

[224] Jean-Dominique Gallezot, Beata Planeta, Nabeel Nabulsi, Donna Palumbo, Xiaoxi Li, Jing Liu, Carolyn Rowinski, Kristin Chidsey, David Labaree, Jim Ropchan, Shu-Fei Lin, Aarti Sawant-Basak, Timothy J. McCarthy, Anne W. Schmidt, Yiyun Huang, and Richard E. Carson. Determination of receptor occupancy in the presence of mass dose: [11c]gsk189254 pet imaging of histamine h3 receptor occupancy by pf-03654746. Journal of Cerebral Blood Flow and Metabolism: Official Journal of the International Society of Cerebral Blood Flow and Metabolism, 37(3):1095–1107, 3 2017. ISSN 1559-7016. doi:10.1177/0271678X16650697. PMID: 27207170 PMCID: PMC5363483.

[225] Deepak Cyril D’Souza, Jose A. Cortes-Briones, Mohini Ranganathan, Halle Thurnauer, Gina Creatura, Toral Surti, Beata Planeta, Alexander Neumeister, Brian Pittman, Marc Normandin, Michael Kapinos, Jim Ropchan, Yiyun Huang, Richard E. Carson, and Patrick D. Skosnik. Rapid changes in cb1 receptor availability in cannabis dependent males after abstinence from cannabis. Biological Psychiatry. Cognitive Neuroscience and Neuroimaging, 1(1):60–67, 1 2016. ISSN 2451-9022. doi:10.1016/j.bpsc.2015.09.008. PMID: 26858993 PMCID: PMC4742341.

[226] Alexander Neumeister, Marc D. Normandin, James W. Murrough, Shannan Henry, Christopher R. Bailey, David A. Luckenbaugh, Keri Tuit, Ming-Qiang Zheng, Isaac R. Galatzer-Levy, Rajita Sinha, Richard E. Carson, Marc N. Potenza, and Yiyun Huang. Positron emission tomography shows elevated cannabinoid cb1 receptor binding in men with alcohol dependence. Alcoholism, Clinical and Experimental Research, 36 (12):2104–2109, 12 2012. ISSN 1530-0277. doi: 10.1111/j.1530-0277.2012.01815.x. PMID: 22551199 PMCID: PMC3418442.

[227] Marc D. Normandin, Ming-Qiang Zheng, Kuo-Shyan Lin, N. Scott Mason, Shu-Fei Lin, Jim Ropchan, David Labaree, Shannan Henry, Wendol A. Williams, Richard E. Carson, Alexander Neumeister, and Yiyun Huang. Imaging the cannabinoid cb1 receptor in humans with [11c]omar: assessment of kinetic analysis methods, testretest reproducibility, and gender differences. Journal of Cerebral Blood Flow and Metabolism: Official Journal of the International Society of Cerebral Blood Flow and Metabolism, 35(8):1313–1322, 8 2015. ISSN 1559-7016. doi:10.1038/jcbfm.2015.46. PMID: 25833345 PMCID: PMC4528005.

[228] Mohini Ranganathan, Jose Cortes-Briones, Rajiv Radhakrishnan, Halle Thurnauer, Beata Planeta, Patrick Skosnik, Hong Gao, David Labaree, Alexander Neumeister, Brian Pittman, Toral Surti, Yiyun Huang, Richard E. Carson, and Deepak Cyril D’Souza. Reduced brain cannabinoid receptor availability in schizophrenia. Biological Psychiatry, 79(12):997–1005, 6 2016. ISSN 1873-2402. doi:10.1016/j.biopsych.2015.08.021. PMID: 26432420 PMCID: PMC4884543.

[229] Tatu Kantonen, Tomi Karjalainen, Janne Isojärvi, Pirjo Nuutila, Jouni Tuisku, Juha Rinne, Jarmo Hietala, Valtteri Kaasinen, Kari Kalliokoski, Harry Scheinin, Jussi Hirvonen, Aki Vehtari, and Lauri Nummenmaa. Interindividual variability and lateralization of mu-opioid receptors in the human brain. NeuroImage, 217:116922, 8 2020. ISSN 1053-8119. doi: 10.1016/j.neuroimage.2020.116922.

[230] Paul L Williams and Randall D Beer. Nonnegative Decomposition of Multivariate Information. arXiv, 2010. doi:10.48550/arXiv.1004.2515. URL http://arxiv.org/abs/1004.2515. 1004.2515.

[231] Erfan Nozari, Maxwell A. Bertolero, Jennifer Stiso, Lorenzo Caciagli, Eli J. Cornblath, Xiaosong He, Arun S. Mahadevan, George J. Pappas, and Dani S. Bassett. Macroscopic resting-state brain dynamics are best described by linear models. Nature Biomedical Engineering, pages 1–17, dec 11 2023. ISSN 2157-846X. doi:10.1038/s41551-023-01117-y. URL https://www.nature.com/articles/s41551-023-01117-y. publisher: Nature Publishing Group.

[232] Adam B Barrett. Exploration of synergistic and redundant information sharing in static and dynamical Gaussian systems. Physical Review E, 91:52802, 2015. doi: 10.1103/PhysRevE.91.052802. [Online; accessed 2020-09-02].

[233] David Balduzzi and Giulio Tononi. Integrated information in discrete dynamical systems: motivation and theoretical framework. PLoS Computational Biology, 4(6): e1000091, 2008. doi:10.1371/journal.pcbi.1000091.

[234] Adam B. Barrett and Anil K. Seth. Practical measures of integrated information for time-series data. PLoS Computational Biology, 7(1):e1001052, 2011. doi:10.1371/journal.pcbi.1001052.

[235] Masafumi Oizumi, Naotsugu Tsuchiya, and Shun-ichi Amari. Unified framework for information integration based on information geometry. Proceedings of the National Academy of Sciences, 113(51):14817–14822, 2016. doi:10.1073/pnas.1603583113.

[236] Pedro A. M. Mediano, Anil K. Seth, and Adam B. Barrett. Measuring integrated information: Comparison of candidate measures in theory and simulation. Entropy, 21 (1):17, 2019. doi:10.3390/e21010017.

[237] Max Tegmark. Improved measures of integrated information. PLoS Computational Biology, 12(11):e1005123, 2016. doi:10.1371/journal.pcbi.1005123.

[238] Joseph T. Lizier, Mikhail Prokopenko, and Albert Y. Zomaya. Information modification and particle collisions in distributed computation. Chaos, 20(3):037109, 2010. doi:10.1063/1.3486801.

[239] Joseph T Lizier, Mikhail Prokopenko, and Albert Y Zomaya. Local measures of information storage in complex distributed computation. Information Sciences, 208: 39–54, 2012.

[240] Thomas Schreiber. Measuring information transfer. Physical review letters, 85(2):461, 2000.

[241] James Massey et al. Causality, feedback and directed information. In Proc. Int. Symp. Inf. Theory Applic.(ISITA-90), volume 2, 1990.

[242] M. Wibral, R. Vicente, and M. Lindner. Transfer entropy in neuroscience. In M. Wibral, R. Vicente, and J. Lizier, editors, Directed Information Measures in Neuroscience, Understanding Complex Systems, pages 1–24. Springer, Berlin, Heidelberg, 2014. doi:10.1007/978-3-642-54474-3_1.

[243] Clive WJ Granger. Investigating causal relations by econometric models and cross-spectral methods. Econometrica: Journal of the Econometric Society, pages 424–438, 1969.

[244] Norbert Wiener. Nonlinear prediction and dynamics. In Jerzy Neyman, editor, Berkeley Symposium on Mathematical Statistics and Probability, Volume 3, pages 247–252. University of California Press, Berkeley, 1956.

[245] Lionel Barnett, Adam B. Barrett, and Anil K. Seth. Granger causality and transfer entropy are equivalent for gaussian variables. Physical Review Letters, 103(23): 238701, 2009. doi:10.1103/PhysRevLett.103.238701.

[246] David Balduzzi and Giulio Tononi. Integrated information in discrete dynamical systems: Motivation and theoretical framework. PLoS Computational Biology, 4(6): 1–18, 06 2008. 10.1371/journal.pcbi.1000091.

[247] John D Murray, Alberto Bernacchia, David J Freedman, Ranulfo Romo, Jonathan D Wallis, Xinying Cai, Camillo Padoa-Schioppa, Tatiana Pasternak, Hyojung Seo, Daeyeol Lee, et al. A hierarchy of intrinsic timescales across primate cortex. Nature neuroscience, 17(12):1661–1663, 2014.

[248] M. E. J. Newman. Modularity and community structure in networks. Proceedings of the National Academy of Sciences of the United States of America, 103(23):8577– 8582, 2006. doi:10.1073/pnas.0601602103.

[249] Mark D. Humphries and Kevin Gurney. Network ‘small-world-ness’: A quantitative method for determining canonical network equivalence. PLOS ONE, 3(4):e2051, 2008. doi:10.1371/journal.pone.0002051.

[250] Enzo Tagliazucchi, Pablo Balenzuela, Daniel Fraiman, and Dante R. Chialvo. Criticality in large-scale brain fmri dynamics unveiled by a novel point process analysis. Frontiers in Physiology, 3 FEB:1–12, 2012. ISSN 1664042X. doi:10.3389/fphys.2012.00015. PMID: 22347863 Citation Key: Tagliazucchi2012 ISBN: 1664-042X.

[251] Mikail Rubinov and Olaff Sporns. Complex network measures of brain connectivity: Uses and interpretations. NeuroImage, 52(3):1059–1069, 2010. ISSN 10538119. doi:10.1016/j.neuroimage.2009.10.003. URL http://dx.doi.org/10.1016/j.neuroimage.2009.10.003. PMID: 19819337 Citation Key: Rubinov2010 ISBN: 1095-9572 (Electronic)\r1053-8119 (Linking).

[252] Reinder Vos de Wael, Oualid Benkarim, Casey Paquola, Sara Lariviere, Jessica Royer, Shahin Tavakol, Ting Xu, Seok-Jun Hong, Georg Langs, Sofie Valk, et al. Brainspace: a toolbox for the analysis of macroscale gradients in neuroimaging and connectomics datasets. Communications biology, 3(1):103, 2020.

[253] Andrea I. Luppi, Fernando E. Rosas, Pedro A. M. Mediano, Athena Demertzi, David K. Menon, and Emmanuel A. Stamatakis. Unravelling consciousness and brain function through the lens of time, space, and information. Trends in Neurosciences, 47(7):551–568, jul 1 2024. ISSN 0166-2236, 1878-108X. doi:10.1016/j.tins.2024.05.007. URL https://www.cell.com/trends/neurosciences/abstract/S0166-2236(24)00087-0. publisher: Elsevier PMID: 38824075.

[254] Ronald R Coifman, Stephane Lafon, Ann B Lee, Mauro Maggioni, Boaz Nadler, Frederick Warner, and Steven W Zucker. Geometric diffusions as a tool for harmonic analysis and structure definition of data: Diffusion maps. Proc Natl Acad Sci USA, 102(21):7426–7431, 2005.

[255] Larry V Hedges. Distribution theory for glass’s estimator of effect size and related estimators. journal of Educational Statistics, 6(2):107–128, 1981.

[256] Harald Hentschke and Maik C Stüttgen. Computation of measures of effect size for neuroscience data sets. European Journal of Neuroscience, 34(12):1887–1894, 2011.

[257] Ross D Markello and Bratislav Misic. Comparing spatial null models for brain maps. NeuroImage, 236:118052, 2021.

[258] Russell A Poldrack, Aniket Kittur, Donald Kalar, Eric Miller, Christian Seppa, Yolanda Gil, D Stott Parker, Fred W Sabb, and Robert M Bilder. The cognitive atlas: toward a knowledge foundation for cognitive neuroscience. Frontiers in neuroinformatics, 5:17, 2011.

[259] Aaron F Alexander-Bloch, Haochang Shou, Siyuan Liu, Theodore D Satterthwaite, David C Glahn, Russell T Shinohara, Simon N Vandekar, and Armin Raznahan. On testing for spatial correspondence between maps of human brain structure and function. NeuroImage, 178: 540–551, 2018.

[260] Justine Y. Hansen, Ross D. Markello, Jacob W. Vogel, Jakob Seidlitz, Danilo Bzdok, and Bratislav Misic. Mapping gene transcription and neurocognition across human neocortex. Nature Human Behaviour, pages 1–11, 3 2021. ISSN 2397-3374. doi:10.1038/s41562-021-01082-z. publisher: Nature Publishing Group.

[261] Yoav Benjamini and Yosef Hochberg. Controlling the false discovery rate: a practical and powerful approach to multiple testing. J Roy Stat Soc B, 57(1):289–300, 1995.

[262] B.T. Thomas Yeo, Fenna M. Krienen, Jorge Sepulcre, Mert R. Sabuncu, D. Lashkari, Marisa Hollinshead, Joshua L. Roffman, Jordan W. Smoller, Lilla Zollei, Jonathan R. Polimeni, Bruce Fischl, Hesheng Liu, and Randy L. Buckner. The organization of the human cerebral cortex estimated by intrinsic functional connectivity. Journal of neurophysiology, 106:1125–1165, 2011. ISSN 1522-1598. doi:10.1152/jn.00338.2011. PMID: 21653723 Citation Key: Yeo2011 ISBN: 1522-1598 (Electronic)\r0022-3077 (Linking).

[263] Jason Z Kim, Jonathan M. Soffer, Ari E Kahn, Jean M Vettel, Fabio Pasqualetti, and Danielle S Bassett. Role of graph architecture in controlling dynamical networks with applications to neural systems. Nature Physics, 14(1):91–98, 2018. ISSN 17452481. doi:10.1038/NPHYS4268.

[264] Ross D Markello, Justine Y Hansen, Zhen-Qi Liu, Vincent Bazinet, Golia Shafiei, Laura E Suárez, Nadia Blostein, Jakob Seidlitz, Sylvain Baillet, Theodore D Satterthwaite, et al. Neuromaps: structural and functional interpretation of brain maps. Nature Methods, 19(11):1472–1479, 2022.

