## Supplementary Figures for "An engine for systematic discovery of cause-effect relationships between brain structure and function"

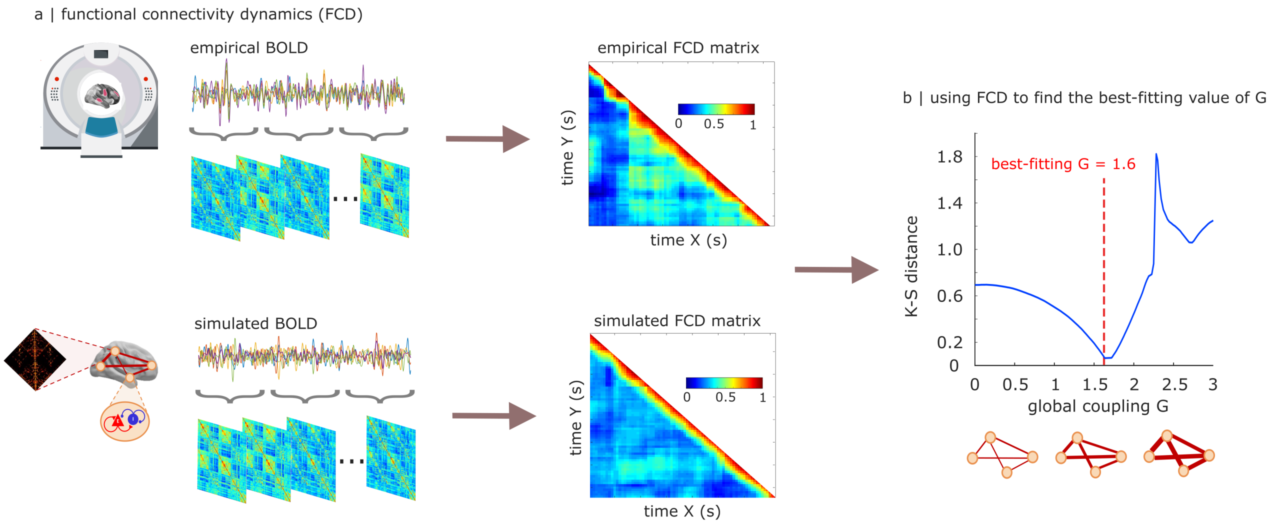


Figure S1. **Finding the optimal value of the dynamic mean-field model's global effective coupling parameter, *G* |** (**a**) For both empirical and simulated BOLD signals, a time-versus-time matrix of functional connectivity dynamics (FCD) is computed by correlating the time-dependent FC matrices centred at each timepoint. (**b**) Across values of the global effective coupling parameter $G$, we compute the KS-distance between the empirical and simulated FCD. A Bayesian optimiser [64] is used to sample different values of $G$ and identify the point where the KS distance between empirical and simulated FCD is minimised, representing the point of best fit. The resulting value (here: $G = 1.6$) is used to scale the structural connectome for generating simulated time-series for the tuned model. Abscissa: $G$ values. Ordinate: KS distance (lower is better).

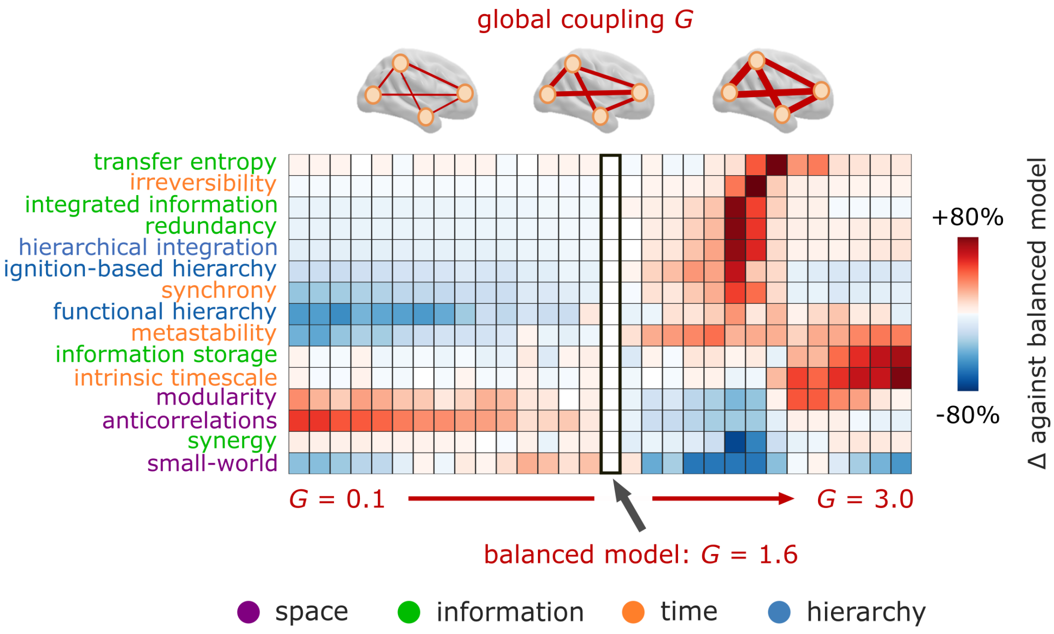


Figure S2. **Functional consequences of scaling the global effective coupling |** The global level of effective coupling has drastically variable effects across different aspects of functional brain organisation, with measures such as metastability and integrated information peaking around the same level of effective coupling. We compute our battery of functional measures of brain organisation using values of effective coupling $G$ ranging from 0.1 to 3.0, in increments of 0.1. Heatmap shows percentage changes in each functional measure, using as reference the value of each measure obtained for $G = 1.6$, corresponding to the balanced model based on minimal KS distance between empirical and simulated FCD (Fig. S1). Different dimensions of brain function are optimised for different values of effective coupling, revealing the existence of complex trade-offs.

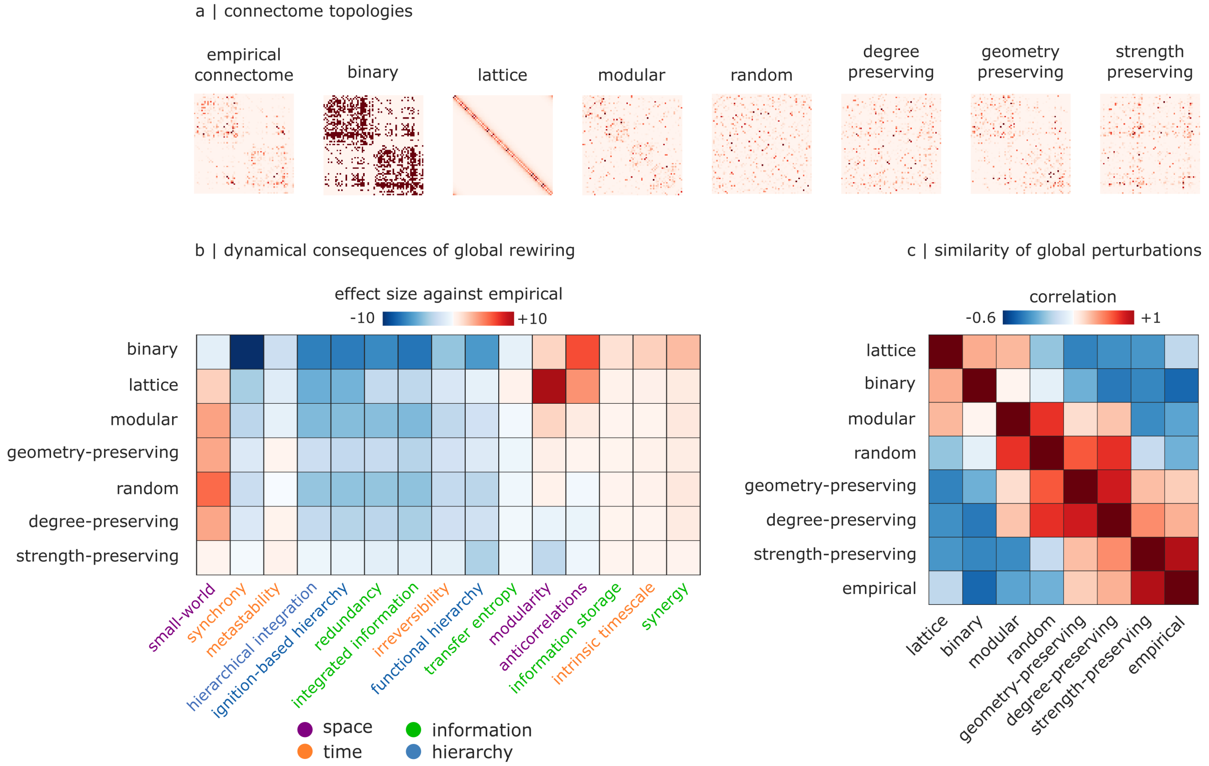


Figure S3. **Functional consequences of perturbing global network topology of the structural connectome |** (**a**) The empirical human structural connectome is compared against a binary network, a lattice network, a modular network (stochastic block model), and four types of random network: fully random, degree-preserving, degree- and strength-preserving, and geometry-preserving (i.e., preserving degree and connection length to respect the spatial embedding of the empirical connectome). (**b**) Heatmap shows the effect size (Hedge's $g$) of each perturbation of global network topology, on each measure of functional brain organisation. (**c**) Increasingly severe global perturbations induce increasing deviations from the dynamical properties of the empirical human connectome.

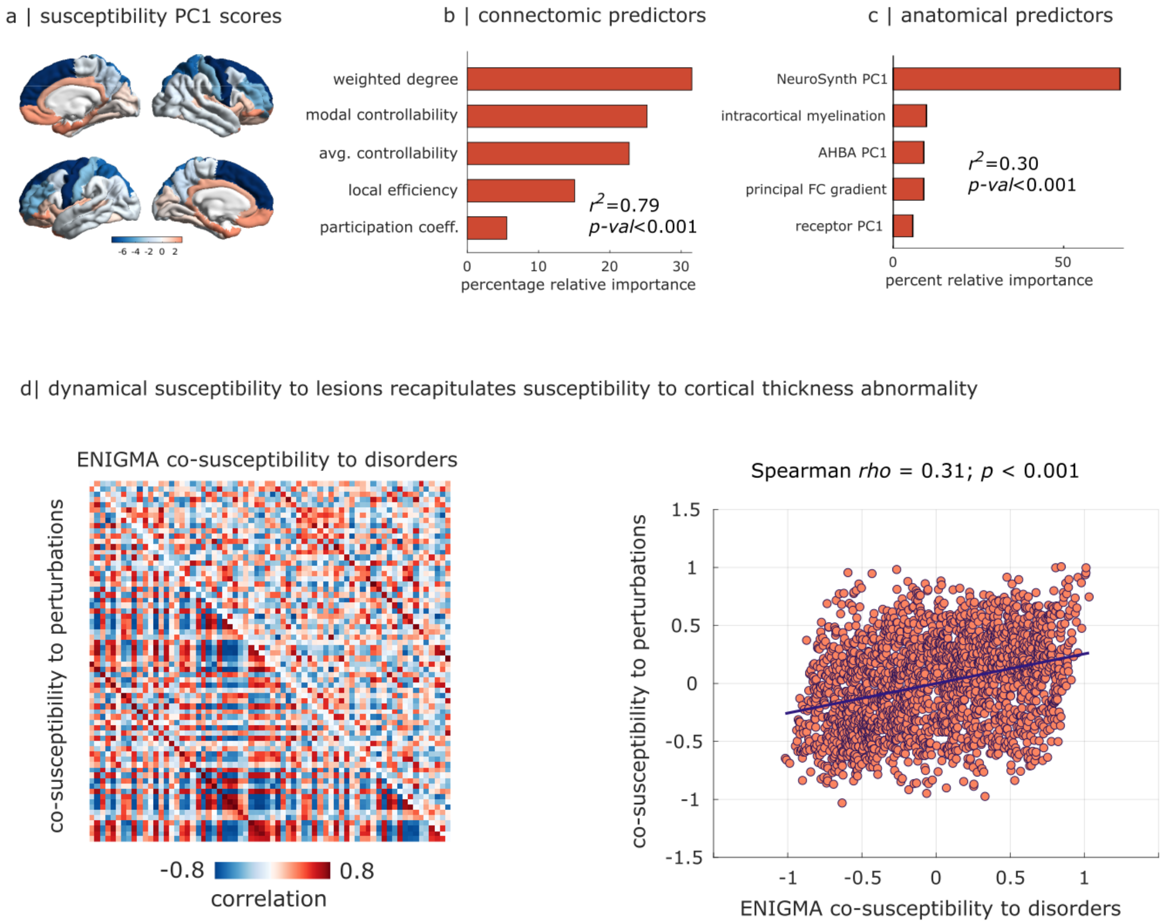


Figure S4. **Biological predictors of functional susceptibility to local perturbations |** (**a**) Principal component of dynamical susceptibility to local structural lesions. (**b**) Dominance analysis predicting the principal component of dynamical co-susceptibility to structural lesions from (a) based on local features of the empirical structural connectome (weighted degree, participation coefficient, local efficiency, and average and modal controllability), expressed in terms of percentage of relative importance. The variance explained is significantly greater than what would be explained based on null maps with preserved spatial autocorrelation ($R^2 = 0.79, p < 0.001$). (**c**) Dominance analysis predicting the principal component of dynamical co-susceptibility to structural lesions from (a) based on macroscale features of cortical functional and anatomical organisation: intracortical myelination (T1w:T2w ratio); cortical thickness; principal component of gene expression from the Allen Human Brain Atlas database (AHBA PC1); principal component of meta-analytic activation from the NeuroSynth database (NeuroSynth PC1); and principal gradient of functional connectivity. The variance explained ($R^2$) is significantly greater than what would be explained based on null maps with preserved spatial autocorrelation ($R^2 = 0.30, p < 0.001$). (**d**) Lower triangular: regional co-susceptibility to lesion-induced dynamical alterations, computed as the correlation between each pair of regions across all dynamical alterations. Upper triangular: co-susceptibility to disorders from the ENIGMA consortium, computed as the correlation between each pair of regions across all disorder-related changes in cortical thickness. Scatter plot: significant correlation between the two co-susceptibility matrices (Spearman's $rho = 0.31; p < 0.001$), after removing exponential distance-dependence.

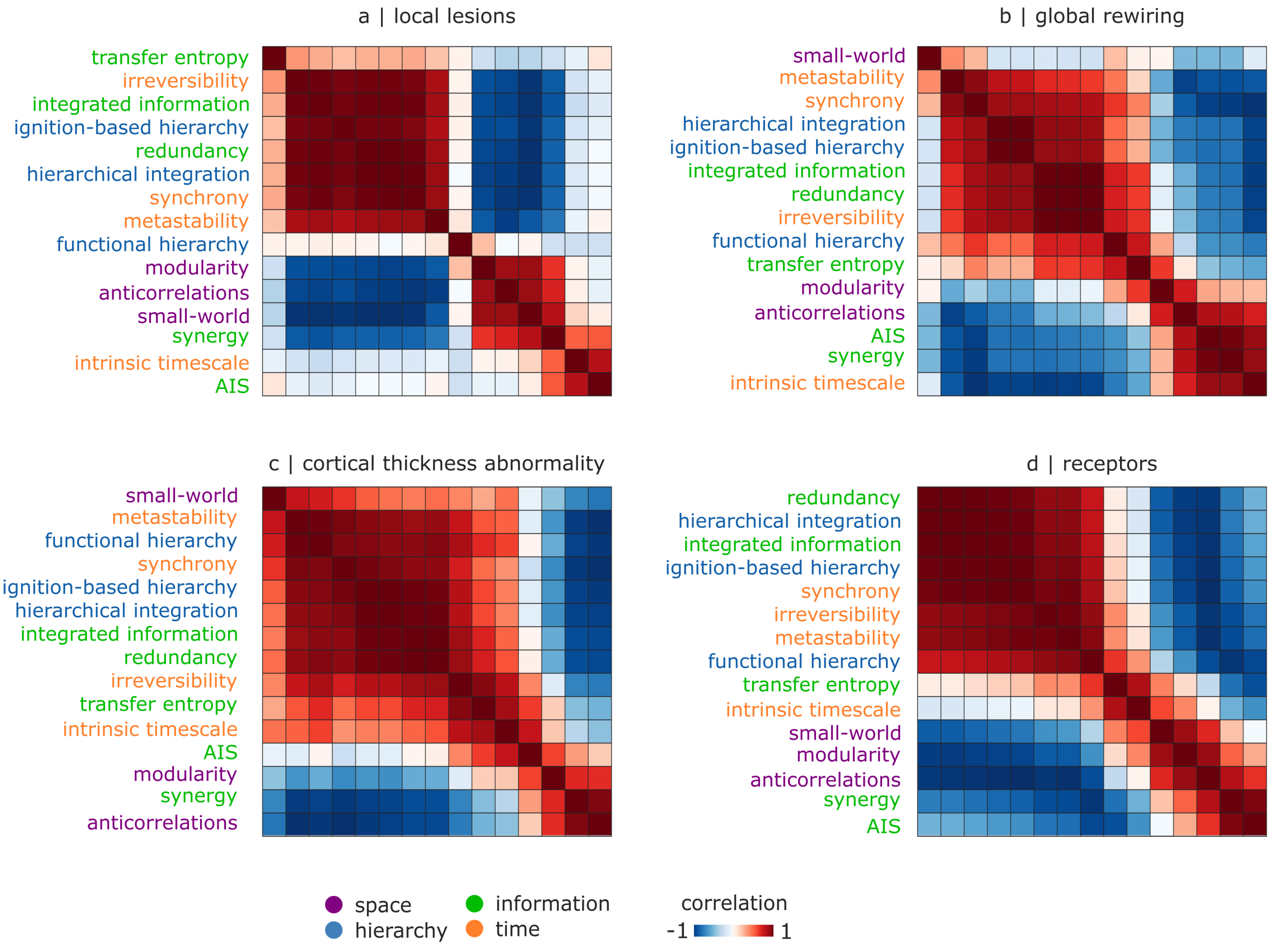


Figure S5. **Two broad patterns of brain function |** **(a)** Similarity of functional measures' response to local structural lesions. **(b)** Similarity of functional measures' response to global rewirings of the structural connectome. **(c)** Similarity of functional measures' response to regional changes in local biophysics according to gradients of cortical thickness abnormality. **(d)** Similarity of functional measures' response to neuromodulation according to empirical receptor density from *in vivo* PET.

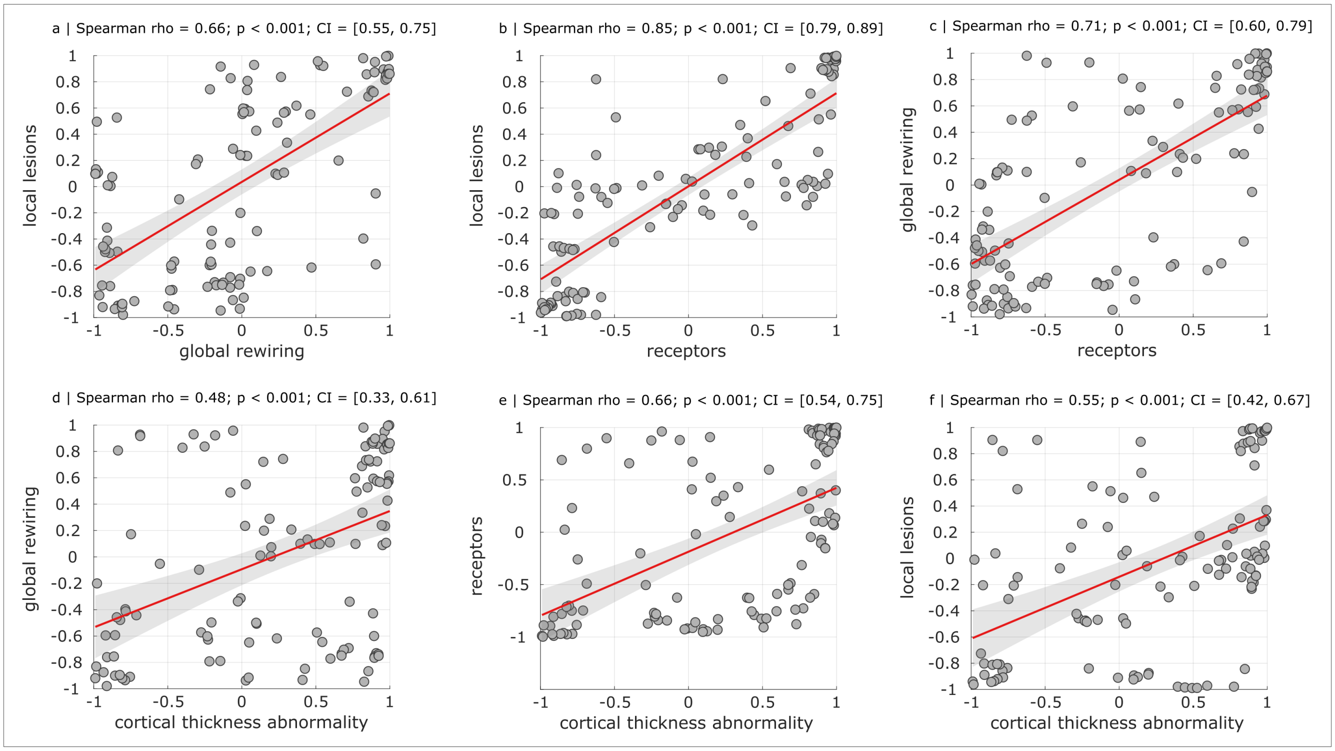


Figure S6. **Clustering of measures' responses to perturbations is consistent across perturbation types|** **(a)** Similarity of the clustering of functional measures' changes in response to local lesions and global rewiring. **(b)** Similarity of the clustering of functional measures' changes in response to local lesions and neuromodulation according to empirical receptor density from *in vivo* PET. **(c)** Similarity of the clustering of functional measures' changes in response to global rewiring and neuromodulation according to empirical receptor density from *in vivo* PET. **(d)** Similarity of the clustering of functional measures' changes in response to global rewiring and cortical thickness abnormality. **(e)** Similarity of the clustering of functional measures' changes in response to neuromodulation and cortical thickness abnormality. **(d)** Similarity of the clustering of functional measures' changes in response to local lesions and cortical thickness abnormality.

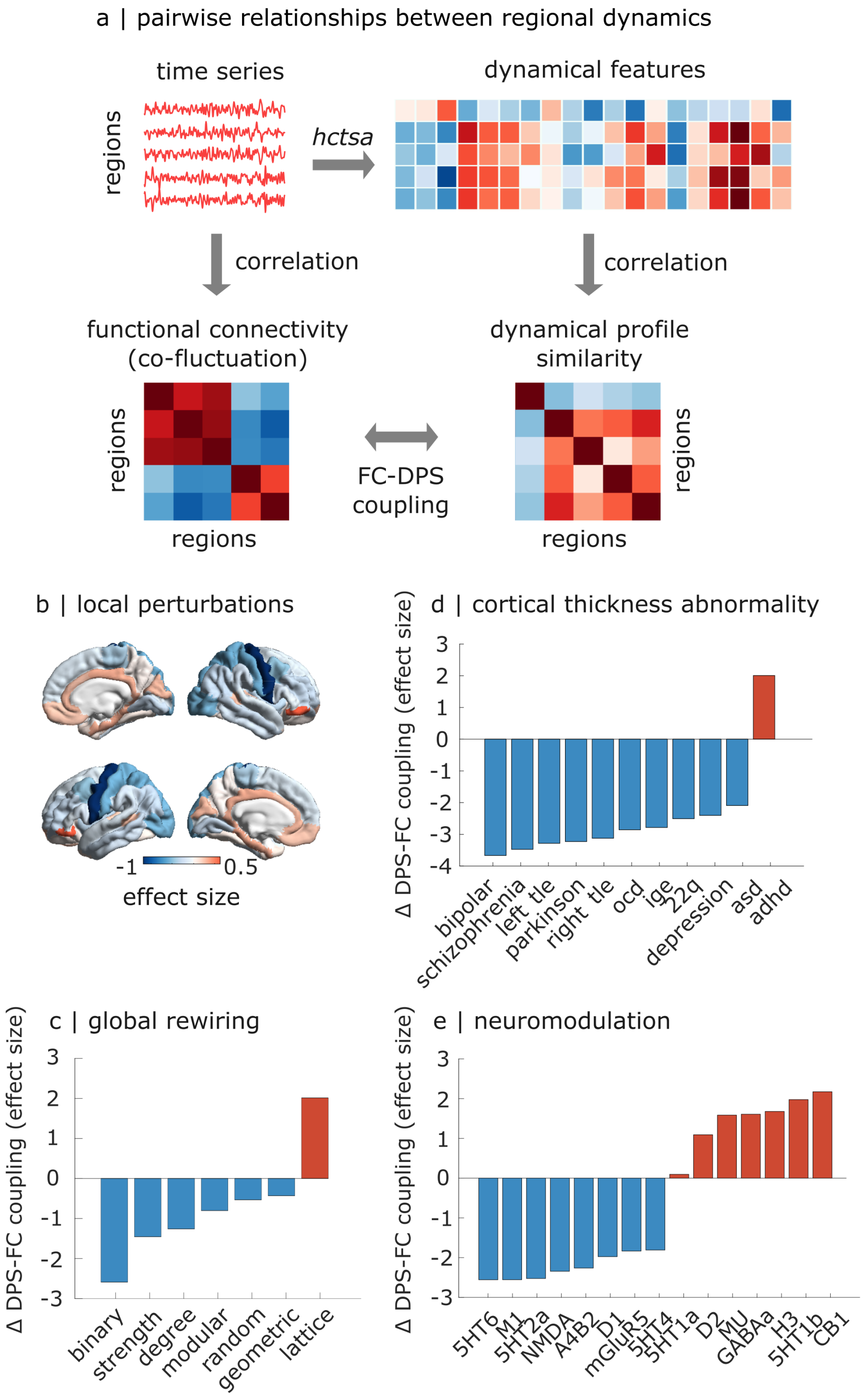


Figure S7. **Modulating the coupling between temporal and feature-based representations of inter-regional interactions |** (**a**) Different ways of characterising pairwise similarity between regional time-series. Functional connectivity (FC) is the correlation between two regions' time-series over time, reflecting co-fluctuation. Dynamical profile similarity (DPS) is the correlation between the dynamical features of two regions' time-series. For both FC and DPS, the result is a region-by-region matrix of pairwise similarities. (**b**) For each cortical region, we show the change in FC-DPS coupling that is observed after lesioning that region's structural connectivity. (**c**) Change in FC-DPS coupling as a function of global network rewiring. (**d**) Change in FC-DPS coupling as a function of reshaping intrinsic regional excitation according to empirical patterns of cortical thickness abnormality. (**e**) Change in FC-DPS coupling as a function of modulating the regional excitatory or inhibitory gain according to empirical patterns of cortical receptor density.
